# Isosteric Engineering of Enzymes: Overcoming Activity–Stability Trade-offs by Site-Selective CH → N Substitutions

**DOI:** 10.64898/2026.02.24.707619

**Authors:** Elwy H. Abdelkader, Haocheng Qianzhu, Gottfried Otting, Thomas Huber

## Abstract

Enzymes used on industrial scale are routinely engineered for best performance. However, exhaustive mutagenesis campaigns using the twenty canonical proteinogenic amino acids rapidly reach an evolutionary ceiling, where gains in activity compromise other critical properties such as thermal endurance. Although non-canonical amino acids (ncAA) expand the chemical space, most are costly for use on an industrial scale and significantly perturb structure. Here, we demonstrate that the evolutionary ceiling of highly optimized polyethylene terephthalate (PET) hydrolases (PETases) can be broken with azatryptophans that (i) differ minimally from their canonical tryptophan, (ii) are genetically encoded, and (iii) are produced in high yield by enzymatic biosynthesis from inexpensive precursors. The first genetic encoding systems are described for 4-azatryptophan, 5-azatryptophan, and 6-azatryptophan, achieving single, site-selective isosteric CH → N substitutions that enhancing the catalytic activity while preserving thermal stability. The fluorescence of 6AW provides a uniquely sensitive reporter of side-chain solvent exposure, which is critical for PETase activity and shown to vary between five different PETases. Furthermore, Azatryptophan-bearing enzymes are inexpensive to produce. To benchmark PETase activity, a rapid fluorescence-based kinetic assay, **PETra**, is introduced, which delivers consistency and reproducibility by using a soluble substrate yet correlates strongly with the hydrolysis of solid PET.

## Introduction

Polyethylene terephthalate (PET) waste is a major contributor to global environmental pollution due to its high production volume and resistance to biodegradation. Since the discovery of *Ideonella sakaiensis* PET hydrolase (*Is*PETase) in 2016, intensive efforts have sought to identify and engineer PET-degrading enzymes with improved activity, particularly at elevated temperatures.^[1,2]^ A persistent barrier to progress is the activity–stability trade-off, where mutations that accelerate catalysis often increase conformational flexibility and reduce thermal stability, whereas highly thermostable variants can be too rigid to support efficient turnover.^[3-5]^ Trp185 is a highly conserved residue that lines the PET hydrolase substrate-binding cleft and exhibits pronounced conformational flexibility that is believed to support substrate recognition, transition-state stabilization, and product release. However, this mobility, which has been documented across multiple crystal structures, also contributes to the activity–stability trade-off in PET hydrolase optimization.^[6,7]^

To further improve the performance of highly engineered PET hydrolases currently available and to transcend the limits imposed by canonical amino acids, we turned to isosteric substitutions at Trp185. Isosterism posits that compounds sharing the same number and arrangement of electrons display similar physical properties despite different nuclei,^[8]^ and it forms a cornerstone of lead optimization in medicinal chemistry.^[9]^ Here we invert the classical ligand-centric use of isosteric replacement by applying the concept to the enzyme itself. Using genetic code expansion (GCE) to circumvent the constraints of the canonical amino acid repertoire,^[10]^ we introduce isosteric azatryptophans, 4-azatryptophan (4AW), 5-azatryptophan (5AW), 6-azatryptophan (6AW), and 7-azatryptophan (7AW), at position 185 across multiple PET hydrolase scaffolds. Expanding on our recent demonstration of 7AW incorporation,^[11]^ we here present genetic encoding systems for 4AW, 5AW, and 6AW, describe their spectroscopic properties, and explore their potential to enhance PET hydrolase function. We also present biosynthesis protocols for these azatryptophan isosteres, which renders this engineering strategy affordable and feasible on a large scale. To quantitatively assess enzyme performance, we further introduce **PETra**, a fluorescence-based kinetic assay using a soluble PET analogue that enables reliable quantification of intrinsic catalytic parameters and correlates strongly with PET film degradation.

We refer to the PETases containing azatryptophan instead of tryptophan as azaPETases. The azaPETases provide site-selective insight into side-chain flexibility, yield measurable gains in PET degradation, improve substrate affinity, and produce favourable shifts in product distribution, all while maintaining thermal stability and thereby mitigating the activity–stability trade-off.

## Results and Discussion

### Scalable Biosynthesis and Site-specific Genetic Encoding of Azatryptophans

In previous work, we achieved the site-specific incorporation of 7AW in response to an amber stop codon using a mutant pyrrolysyl-tRNA synthetase from the methanogenic archaeon ISO4-G1 (G1PylRS).^[11]^ To extend this platform to the 4AW, 5AW, and 6AW isomers in a cost-effective manner (Figure 1A), we used the engineered tryptophan synthase β-subunit *Tm*9D8* mutant from *Thermotoga maritima*, which catalyses the formation of diverse tryptophan analogues from serine and indole precursors (Figure 1B).^[11-13]^ By replacing commercially available, chemically synthesised azatryptophans (USD 70,000–480,000 per mol) with their low-cost biosynthetic precursors (USD 30–330 per mol), this approach reduces azatryptophan raw-material costs by ∼1000-fold and enables multigram-scale protein production (Figures S1 and S2 and Tables S1 and S2).

**Figure 1.**
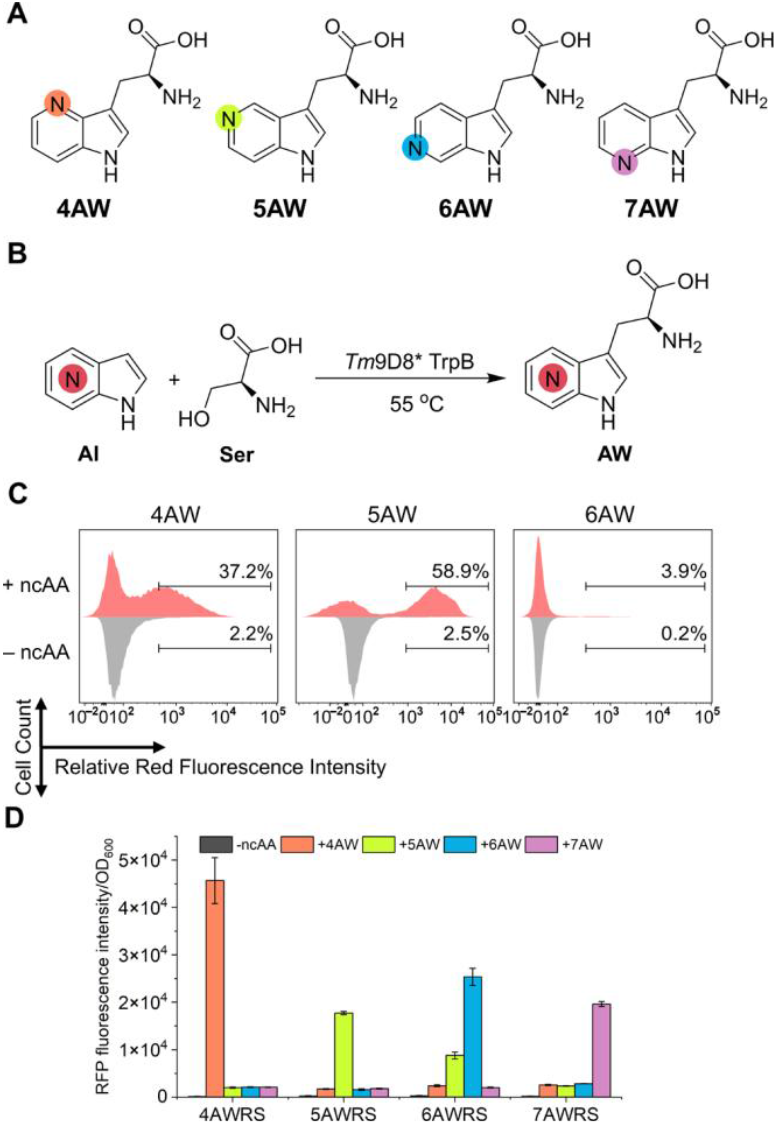
Enhancing PET hydrolase activities via genetically encoded azatryptophans. (A) Chemical structures of 4AW, 5AW, 6AW, and 7AW. The photophysical properties are reported in Figure S2 and Table S2. (B) Enzymatic synthesis of azatryptophans. The *Tm*9D8* TrpB enzyme catalyzes the conversion of azaindole (AI) and serine (Ser) into the corresponding azatryptophan (AW). (C) Identification of G1PylRS mutants specific for 4AW, 5AW, and 6AW. The final-round FACS histograms show enrichment of active, isomer-specific mutants. The x- and y-axes are RFP fluorescence from an amber-interrupted RFP reporter and cell count, respectively. Traces with ncAA (red, plotted upward) and without ncAA (grey, plotted downward) indicate ncAA-dependent amber suppression, confirming RS specificity. (D) Cross-selectivity of 4AWRS, 5AWRS, 6AWRS, and 7AWRS.^[11]^ across azatryptophan isomers. OD_600_-normalized RFP fluorescence from an amber reporter after growth with or without 1 mM of the indicated azatryptophan. Data are shown as the mean of biological triplicates ± standard deviation.

With the azatryptophans in hand, we screened an established G1PylRS mutant library by fluorescence-activated cell sorting (FACS) to identify functional variants selective for each isomer.^[11,14-17]^ The selection strategy alternated positive rounds (in the presence of the noncanonical amino acid (ncAA)) to enrich functional synthetases with negative rounds (in the absence of ncAA) to eliminate non-orthogonal variants. *E. coli* DH10B cells co-expressing the pBK-G1RS library and a His_6_-TAG-RFP reporter plasmid were iteratively sorted for high fluorescence with ncAA and minimal signal without ncAA, producing the expected ncAA-dependent amber suppression phenotype (Figures 1C, S3-S5, and Table S3).

Following several rounds of selection, sixty high-fluorescence colonies per azatryptophan were isolated and sequenced, revealing the best-performing variants: 4AWRS (N165A, V167F, Y204W, A221H, W237T), 5AWRS (N165D, V167F, Y204W, A221S, W237S), and 6AWRS (L124T, Y125L, N165D, V167A, Y204W, A221S, W237W). Each tRNA synthetase exhibited strong discrimination against canonical amino acids, including tryptophan, and high specificity for its cognate azatryptophan isomer (Figure 1D).

To validate incorporation fidelity, we expressed the peptide ENLYFQGDX (X = azatryptophan) fused to the C-terminus of the NT* solubility tag,^[18]^ with the azatryptophan encoded by an amber stop codon.^[11,17]^ After expression, purification, and TEV protease cleavage, the resulting GDX tripeptides were isolated. 1D ^1^H-NMR spectra confirmed clean and exclusive incorporation of the intended azatryptophan isomer, with no detectable misincorporation of canonical tryptophan (Figure S6). To assess potential azatryptophan mischarging by the endogenous *E. coli* tryptophanyl-tRNA synthetase, we expressed the NT* construct with the amber stop codon at the 3′ end replaced by the canonical tryptophan codon in cultures supplemented with 1 mM azatryptophan. The resulting GDW tripeptide contained no azatryptophan signal in the NMR spectra, demonstrating that the azatryptophan isomers are not readily misincorporated at native tryptophan codons under rich-media conditions (Figure S7).

### A Conformational Latch Controlling Trp185 Mobility Revealed by Azatryptophan Probes

FAST-PETase (FAST-P),^[19]^ Depo-PETase (Depo-P),^[23]^ Hot-PETase (Hot-P),^[20]^ LCC-ICCG,^[24]^ and Kubu-PETase (Kubu-P)^[22]^ represent some of the most effective PET hydrolases identified to date for the depolymerization of PET into its constituent monomers bis(2-hydroxyethyl) terephthalate (BHET), mono(2-hydroxyethyl)terephthalic acid (MHET), terephthalic acid (TPA), and ethylene glycol (Figure 2A). The first PETase was discovered in the bacterium *Ideonella sakaiensis* (*Is*PETase) and exhibits a temperature optimum of 40 °C, above which its activity rapidly declines owing to limited thermal stability.^[1]^ FAST-PETase differs from *Is*PETase by five mutations that increase the thermal stability at elevated temperatures and enhance activity at lower temperatures, achieving a temperature optimum of 50 °C.^[19]^ Depo-PETase contains seven mutations that emerged from high-throughput screening of *Is*PETase libraries.^[23]^ HotP-ETase carries a more extensive set of mutations that elevate its temperature optimum to 60 °C.^[20]^ LCC-ICCG shares a similar fold with *Is*PETase but was derived from leaf-branch compost cutinase (LCC) and was engineered for high activity and thermal stability, with a temperature optimum of 70–80 °C.^[24]^ Kubu-PETase was recently identified through large-scale landscape profiling of nearly 2,000 putative PET hydrolases and was subsequently engineered to achieve an optimum of 60–70 °C.^[22]^

**Figure 2.**
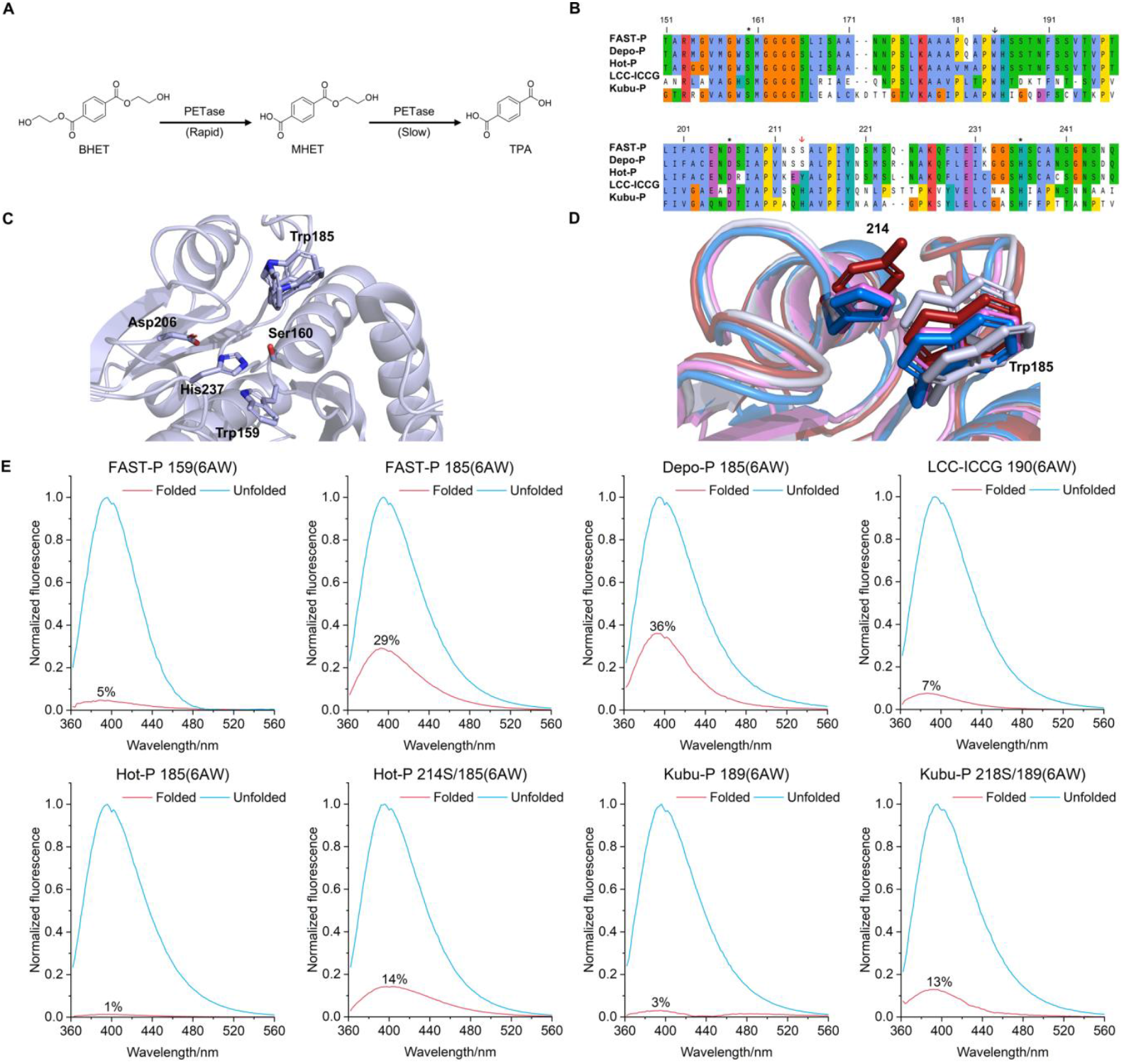
Probing Trp185 flexibility in PET hydrolases using 6AW fluorescence. (A) Hydrolysis pathway of BHET by PET hydrolases. Sequence alignment of the key residues in the PET hydrolases investigated in this study, highlighting the wobbling tryptophan (black arrow) and residue 214 (red arrow). The catalytic Ser-Asp-His triad is marked with asterisks. Full sequence alignments with native numbering are shown in Figure S8. (C) Crystal structure of wild-type *Is*PETase (PDB ID: 5XG0),^[3]^ highlighting the catalytic triad and the positions of Trp159 and Trp185. (D) Structural alignment of FAST-PETase (grey, PDB ID: 7SH6),^[19]^ Hot-PETase (red, PDB ID: 7QVH),^[20]^ LCC-ICCG (blue, PDB ID: 8OTA),^[21]^ and Kubu-PETase (violet, PDB ID: 8YTW).^[22]^ FAST-PETase shows Trp185 in two of the conformations found in wild-type *Is*PETase, whereas the other enzymes show a single conformation, consistent with bulkier residues at position 214 restricting Trp185 mobility. (E) Fluorescence emission spectra of PET hydrolases in which Trp185 is replaced by 6AW. Samples were excited at 330 nm and spectra are normalized to the maximum fluorescence intensity of the urea-unfolded proteins. The flexibility factors (percent) report the relative fluorescence intensity in the folded state.

Co-crystal structures with PET fragments and analogues reveal a conserved catalytic triad comprising Ser160, Asp206, and His237 in *Is*PETase variants; the equivalent residues in LCC-ICCG and Kubu-PETase are Ser165/Asp210/His242 and Ser162/Asp210/His240, respectively. For consistency, all PET hydrolases in this study are described using the sequence numbering of *Is*PETase (Figure 2B; see Figure S8 for the sequence alignment with the original sequence numbering). Apo-structures show a distinct substrate binding cleft (Figure 2C).^[19,25]^ In *Is*PETase-derived enzymes, two tryptophan residues, Trp159 and Trp185, line the substrate binding cleft. Trp185 is solvent-accessible and highly conserved among different PET hydrolases, whereas Trp159 is partially buried and substituted by His164 in LCC-ICCG.

Trp185 is essential for catalysis, as evidenced by the W185A variant of *Is*PETase, which exhibits severely reduced activity.^[26]^ Crystal structures of *Is*PETase (Figure 2C) and FAST-PETase (Figure 2D) show that the solvent exposure of Trp185 confers substantial conformational flexibility, giving rise to the term “wobbling tryptophan”.^[6,26-29]^ Computational work suggests that this mobility is important for substrate positioning, transition-state stabilization, and product release.^[7,30-32]^ Trp185 also contributes to substrate binding via edge-on contacts with the aromatic ring of PET fragments, as observed in a co-crystal structure of *Is*PETase with an MHET analogue (PDB ID 5XH3)^[26]^ and of LCC-ICCG with MHET (PDB ID 7VVE).^[25]^

To probe Trp185 mobility experimentally, we used 6AW as a fluorescent probe and defined a “flexibility factor” as the ratio of emission intensities (measured at λ_em_ = 400 nm with excitation at λ_ex_ = 330 nm) for the folded versus urea-denatured protein at the same concentration. Emission of 6AW at 400 nm increases with solvent exposure and polarity,^[33]^ thus low flexibility factors indicate a restricted, buried environment, whereas high values reflect solvent exposure and conformational freedom (Figure 2E). In FAST-PETase, 6AW at position 159 gave a flexibility factor of 5%, consistent with limited exposure, while 6AW at position 185 produced a flexibility factor of 29%, indicating greater solvent exposure and conformational freedom. Depo-PETase showed an even higher flexibility factor at position 185 (36%), suggesting larger-amplitude motions than in FAST-PETase, despite both enzymes containing a small serine residue at position 214. In contrast, Hot-PETase exhibited a markedly low flexibility factor (1%). The Hot-PETase crystal structure (Figure 2E) shows the side chain of Trp185 packed against Tyr214, explaining the restricted mobility. Replacing Tyr214 with serine (Hot-PETase 214S) released this constraint, increasing the flexibility factor ∼14-fold to 14% (Figure 2E).

LCC-ICC and Kubu-PETase, in which residue 214 is a histidine, also exhibited low flexibility at position 185 (7% and 3%, respectively). Substituting His214 with serine in Kubu-PETase (Kubu-PETase 218S) increased the flexibility factor more than four-fold, consistent with partial relaxation of the wobbling constraint. Together, these results identify residue 214 as a conformational latch that modulates Trp185 mobility across PET hydrolase scaffolds, with aromatic residues (Tyr/His) restricting motion and serine permitting broader conformational sampling.

### PETra: a Quantitative Fluorescence-based Assay for PET Hydrolase Kinetics

The kinetic analysis of PET hydrolases is challenging because PET is insoluble and structurally heterogeneous, making assays with solid PET substrates laborious and difficult to reproduce.^[34]^ Common surrogate substrates, such as *p*-nitrophenyl esters, enable rapid measurements but are poor mimics of the hydrophobic PET backbone and correlate only weakly with true PET degradation.^[35]^ To overcome these limitations, we developed **PETra** (**PET** hydrolase **r**apid **a**ssay), a continuous fluorescence assay based on the soluble monomeric PET fluorescent analogue bis(2-hydroxyethyl) 2-hydroxyterephthalate (BHET-OH; Figure 3A).^[23]^ **PETra** allows high-sensitivity measurements in microtiter plates and yields accurate kinetic parameters, including the Michaelis constant (*K*_m_) and the catalytic turnover number (*k*_cat_), which are otherwise difficult to obtain with solid PET substrates.

**Figure 3.**
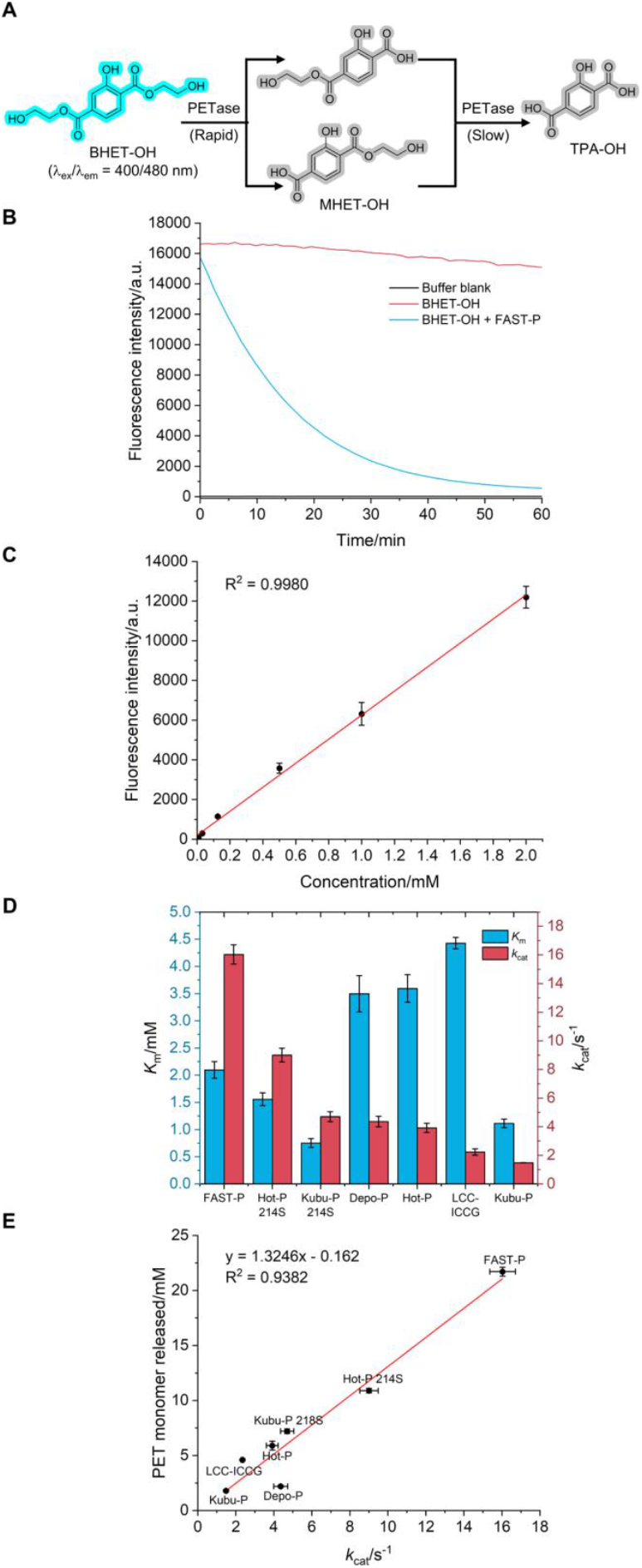
PETra: a fluorescence-based assay for PET hydrolase activity. (A) Hydrolysis pathway of BHET-OH by PET hydrolases. Hydrolysis of BHET-OH is selectively monitored by fluorescence at λ_ex_/λ_em_ = 400/480 nm, as its products MHET-OH and TPA-OH show negligible fluorescence. (B) Kinetic fluorescence traces of BHET-OH hydrolysis by FAST-PETase. Samples (1 mM BHET-OH in 100 mM potassium phosphate buffer, pH 8.0) were incubated with 500 nM FAST-PETase (blue) or buffer alone (red) at 25 °C, and fluorescence was recorded at λ_ex_/λ_em_ = 400/480 nm. (C) Calibration curve of BHET-OH (0.002–2 mM) used in **PETra**, measured at λ_ex_/λ_em_ = 400/480 nm. (D) Michaelis constant (*K*_m_) and catalytic turnover number (*k*_cat_) at 30 °C determined by **PETra** for the PET hydrolases evaluated in this study. (E) Correlation between the PET monomers released from hydrolysing amorphous gf-PET films (6 mm diameter, 7.1 ± 0.1 mg) in 24 h at 50 °C and the *k*_cat_ values measured at 30 °C. Data represent the mean of three independent replicates ± standard deviation.

1D ^1^H-NMR analysis of BHET and BHET-OH hydrolysis by FAST-PETase revealed a two-step pathway (Figures 3A, S10, and S11). The first, rapid cleavage produces MHET and mono(2-hydroxyethyl) 2-hydroxyterephthalate (MHET-OH), respectively. The second, slower step generates TPA and 2-hydroxyterephthalic acid (TPA-OH), consistent with the weak MHETase activity common to PETases.^[36]^ Fluorescence monitoring of BHET-OH hydrolysis by FAST-PETase (λ_ex_/λ_em_ = 400/480 nm) showed a rapid fluorescence loss corresponding to the first ester bond cleavage (Figures 3B), as both MHET-OH and TPA-OH exhibit negligible fluorescence at these wavelengths (Figure S6). Importantly, BHET-OH displayed a linear fluorescence response across concentrations from 0.002–2 mM (R^2^= 0.9980), enabling accurate quantification of its hydrolysis (Figures 3C).

Using **PETra**, we measured initial velocities (*V*_0_) across BHET-OH concentrations and fitted the data to the Michaelis– Menten equation (Figure S7 and S8), yielding *K*_m_ and *V*_max_ values and allowing calculation of *k*_cat_ (*V*_max_/[E]; Table S4). FAST-PETase showed the highest turnover rate (*k*_cat_ ≈ 16 s^-1^), nearly twice that of the next most active PET hydrolase (Figures 3D), and a moderate substrate affinity (*K*_m_ ≈ 2 mM). By contrast, Kubu-PETase displayed stronger substrate affinity (*K*_m_ ≈ 1.1 mM) but much lower catalytic activity (*k*_cat_ ≈ 1.5 s^-1^). Mutations that increase Trp185 flexibility markedly enhanced performance. In Kubu-PETase, introducing the H218S mutation lowered *K*_m_ by approximately one-third and increased *k*_cat_ nearly threefold. In Hot-PETase, the Y214S mutation reduced *K*_m_ by 57% and more than doubled *k*_cat_. These results demonstrate that enhancing Trp185 mobility simultaneously improves substrate affinity and catalytic turnover, yielding substantial activity gains.

To test whether **PETra** faithfully captures the hydrolytic performance of PET hydrolases on solid PET substrates, we quantified the degradation of amorphous Goodfellow PET film (gf-PET) by the same panel of PET hydrolases. The release of PET monomers, defined as the combined amounts of BHET, MHET, and TPA, was measured by complementary spectrophotometric and NMR methods.^[37,38]^ **PETra**-derived *k*_cat_ values correlated strongly with gf-PET film hydrolysis (R^2^ = 0.9382; Figures 3E and S15), whereas *K*_m_ showed no meaningful correlation, producing only a moderate correlation with catalytic efficiency (*k*_cat_/*K*_m_). FAST-PETase proved the most active enzyme both in **PETra** (30 °C) and in gf-PET film hydrolysis (50 °C).

### Trp185 Dynamics Govern the Activity–Stability Trade-off in PET Hydrolases

Thermal stability measurements reveal a clear relationship between Trp185 mobility and structural robustness in PET hydrolases. Variants with rigid Trp185, Hot-PETase and Kubu-PETase, showed substantially higher melting temperatures (Tm) than FAST-PETase and Depo-PETase, which contain more mobile Trp185 (Figure 4A and Table S5). Increasing Trp185 flexibility through Y214S or H218S substitutions in Hot-PETase and Kubu-PETase, respectively, markedly reduced Tm, demonstrating that enhanced wobbling incurs a stability penalty. Although FAST-PETase and Depo-PETase have similar Tm values, their performance on gf-PET at 50 °C diverged substantially. Depo-PETase showed a rapid initial hydrolysis but activity declined after ∼6 h, plateauing by 24 h (Figure 4B). This limited thermal endurance indicates marginal stability at 50 °C, approximately 18 °C below its Tm, and may stem from its markedly elevated Trp185 mobility, as reflected by the higher flexibility factor relative to FAST-PETase. Consequently, its 24 h activity falls below the general correlation with **PETra**-derived *k*_cat_ values measured at 30 °C (Figure 3E), even though the initial linear phase aligns well with **PETra** kinetics (Figure S16).

**Figure4.**
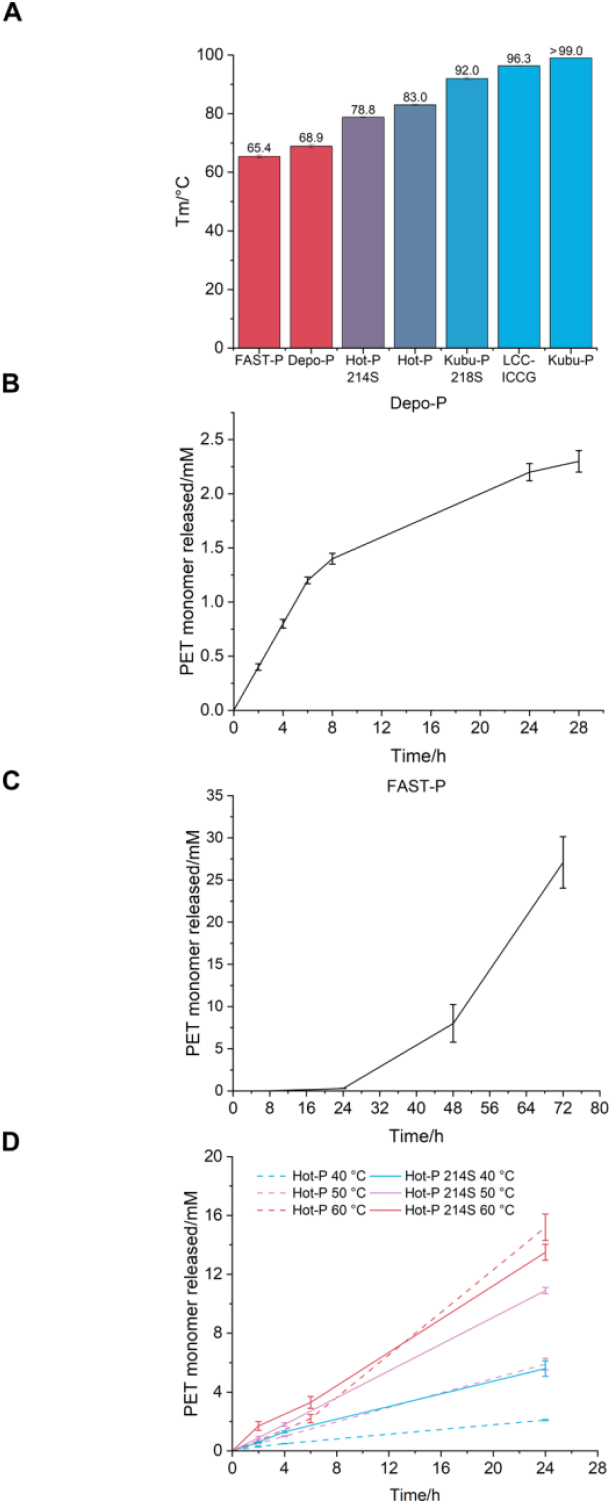
Effect of Trp185 flexibility on the thermal stability and endurance of PET hydrolases. (A) Melting temperatures (Tm) of the PET hydrolase variants determined by thermal denaturation. Time-course hydrolysis of amorphous gf-PET film by Depo-PETase at 50 °C, showing a sharp decline in activity after ∼7 h due to limited thermal stability and endurance. (C) Time-course hydrolysis of amorphous gf-PET film by FAST-PETase at 50 °C, revealing a ∼24 h lag phase followed by sustained product accumulation, indicating extended thermal stability. (D) Comparative analysis of amorphous gf-PET hydrolysis by Hot-PETase (dashed lines) and its Y214S variant (solid lines) at 40 °C, 50 °C, and 60 °C. At lower temperatures (40 °C and 50 °C), the Y214S mutation enhances activity through increased Trp185 flexibility, but this advantage is lost at 60 °C due to reduced thermal endurance. Data shown are the average of three independent replicates ± the standard deviation.

In contrast, FAST-PETase showed a pronounced lag phase, releasing few PET monomers in the first 24 h,^[39-41]^ but then sustained robust hydrolysis over the following 48 h (Figure 4C). Despite having a lower Tm value than Depo-PETase, FAST-PETase therefore exhibits superior thermal endurance at 50 °C, highlighting the distinction between intrinsic thermal stability and functional thermal endurance.^[42]^ These results align with simulation studies suggesting that its superior high-temperature performance derives from enhanced thermal endurance rather than reduced flexibility.^[5]^

To further probe the impact of Trp185 wobbling on temperature-dependent catalysis, we compared the activity of Hot-PETase with its Y214S variant on gf-PET film at 40 °C, 50 °C, and 60 °C (Figure 4D). At 40 °C and 50 °C, both enzymes showed linear release of PET monomers over 24 h, with Hot-PETase Y214S outperforming the parent enzyme by 167% and 85%, respectively. At 60 °C, however, the trend was reversed, with the parent enzyme releasing 13% more monomers over 24 h, indicating that the stability cost of the Y214S mutation manifests at higher temperatures. These results show that Trp185 wobbling is a central determinant of the activity–stability trade-off in PET hydrolases. Aromatic residues at position 214 restrict Trp185 dynamics and increase stability, whereas serine releases the conformational latch, enabling higher catalytic rates but reducing thermal robustness. This behavior reflects a general principle in enzyme engineering: catalytic gains driven by increased conformational plasticity often incur penalties in thermal resilience, necessitating careful balancing of stability and activity in PET hydrolase optimization.^[4,43,44]^

### Isosteric Editing of Trp185 Enhances PET Hydrolase Activity

To examine how isosteric substitutions of the wobbling tryptophan influence PET hydrolase activity, we measured the degradation of amorphous gf-PET films by the Trp185 6AW- and 7AW-substituted azaPETases and compared their performance to the corresponding tryptophan-containing wild-type enzymes (Figure 5A). Across all PET hydrolases tested, replacing Trp185 with 6AW or 7AW increased PET hydrolysis, with 7AW consistently outperforming 6AW. The largest enhancements occurred in scaffolds with higher Trp185 mobility, such as FAST-PETase and Hot-PETase Y214S. In FAST-PETase, 6AW and 7AW substitutions produced sustained increases in PET monomer release over 72 h, yielding 65% and 90% greater hydrolysis, respectively, at pH 8.0 (Figure 5B). Both isosteres also markedly shortened the characteristic lag phase of FAST-PETase at higher enzyme concentrations (Figure 5C). The activity improvements were pH dependent. At pH 7.0, the enhancement by 7AW was reduced, and 6AW incorporation resulted in lower activity relative to the parent enzyme (Figure 5D).

**Figure 5.**
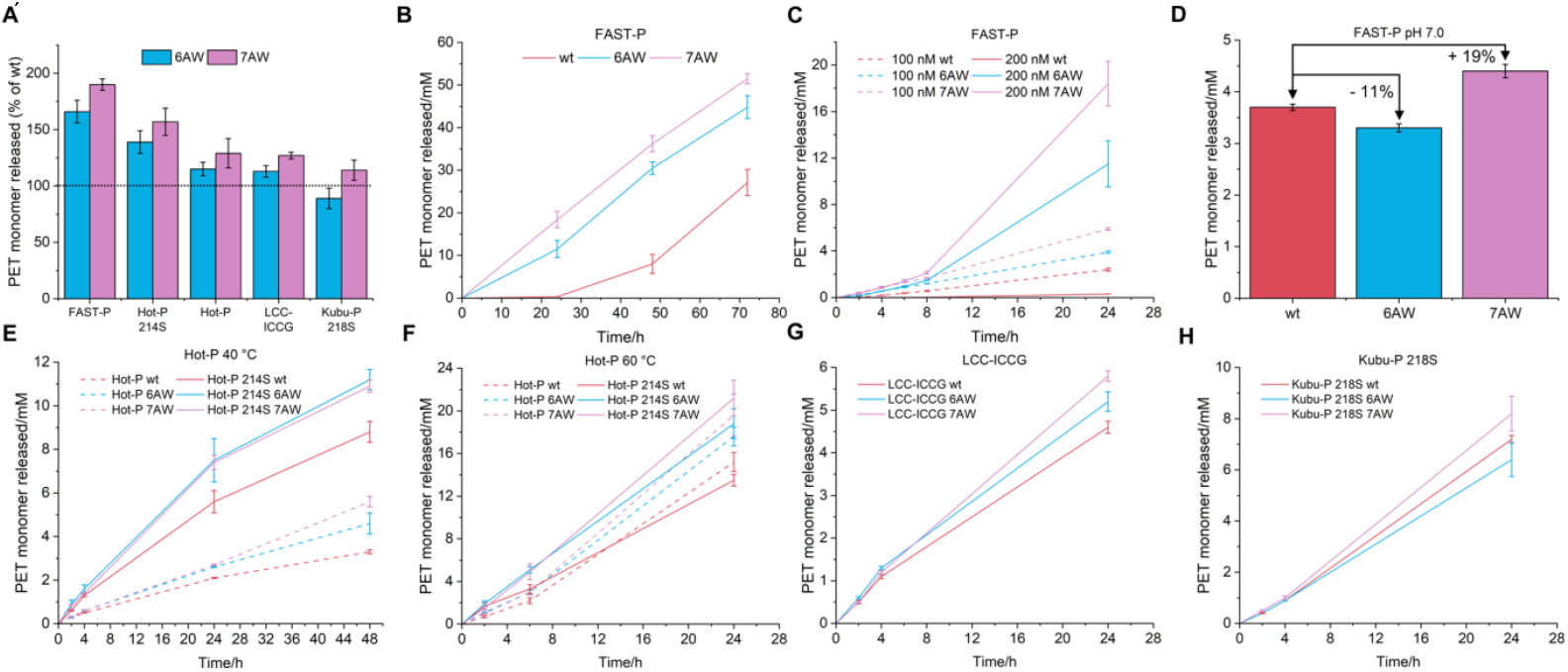
Isosteric replacement of the wobbling tryptophan enhances performance. (A) Comparison of PET degradation activities of the 6AW- and 7AW-substituted azaPETases, measured as the percentage of PET monomers released from gf-PET film hydrolysis after 24 h, relative to their respective tryptophan-containing enzymes tested at 200 nM concentration. Incubation conditions were 50 °C for FAST-PETase, Kubu-PETase 218S, and LCC-ICCG or 60 °C for Hot-PETase and its 214S mutant. (B) Time-course analysis of amorphous gf-PET film hydrolysis by 200 nM FAST-PETase and its Trp185 azaPETases at pH 8.0 and 50 °C, showing faster PET degradation and suppression of the lag phase. (C) Concentration dependence of the lag phase of FAST-PETase and its Trp185 azaPETases at pH 8.0 and 50 °C. (D) Effect of pH on the activity of 100 nM FAST-PETase and its Trp185 azaPETases at 40 °C. (E) Time-course analysis of amorphous gf-PET film hydrolysis by 200 nM Hot-PETase, its azaPETases (dashed lines) compared to Hot-PETase 214S and its azaPETases (solid lines) at pH 9.2 and 60 °C. (F) Same as panel E, but performed at 60 °C. (G) Time-course analysis of amorphous gf-PET hydrolysis by 200 nM LCC-ICCG and its Trp190 azaPETases at pH 9.2 and 50 °C. (H) Time-course analysis of amorphous gf-PET hydrolysis by 200 nM Kubu-PETase 218S and its Trp189 azaPETases at pH 9.2 and 50 °C. All data in the figure represent the mean of three independent replicates ± standard deviation.

Isosteric substitution also enhanced PET hydrolysis in Hot-PETase and Hot-PETase Y214S across multiple temperatures (Figure 5E, F). Remarkably, at 60 °C, both 6AW and 7AW improved the thermal endurance of Hot-PETase Y214S despite exhibiting no significant effect on the Tm values (Table S5). This enhanced thermal endurance allowed the Hot-PETase Y214S azaPETases to outperform the parental Hot-PETase after 24 h (Figure 5F). Finally, 7AW incorporation improved PET degradation by 27% in LCC-ICCG and 14% in Kubu-PETase 218S (Figure 5G, H).

Next, we determined *K*_m_ and *k*_cat_ values for the 6AW- and 7AW-substituted azaPETases using **PETra** (Table S4). Changes in substrate affinity were generally modest (Figure 6A). The most pronounced improvement occurred in LCC-ICCG, where *K*_m_ decreased by 14% with 6AW and 35% with 7AW. Effects on *k*_cat_ showed no uniform trend (Figure 6B). The largest changes were observed in FAST-PETase, where 7AW increased *k*_cat_ by 27%, while 6AW decreased it by 32%. Other variants exhibited smaller changes, within experimental uncertainty. Importantly, azatryptophan substitution had no significant effect on melting temperatures, which suggests that the isosteric replacement preserves the steric profile of the tryptophan side chain. (Table S5). Plotting the thermal stability (Tm) against PET hydrolysis performance (Figure 6C) across the PET hydrolase scaffold panel revealed an inverse correlation consistent with a strong activity– stability trade-off (linear regression, R^2^ = 0.9568). Stable scaffolds such as LCC-ICCG and Kubu-P 218S exhibited high Tm values but low monomer release, whereas FAST-P showed the highest monomer release at comparatively low Tm. The regression line can be viewed as an arbitrary evolutionary ceiling, reflecting the tendency of accessible sequence space to couple stability and activity. Against this background, incorporation of azatryptophans (6AW or 7AW) generally shifted variants toward higher monomer release with only modest changes in Tm relative to the corresponding wild-type enzymes, indicating partial decoupling of activity from stability and, in several cases, movement beyond the constraint implied by the fitted trade-off.

**Figure 6.**
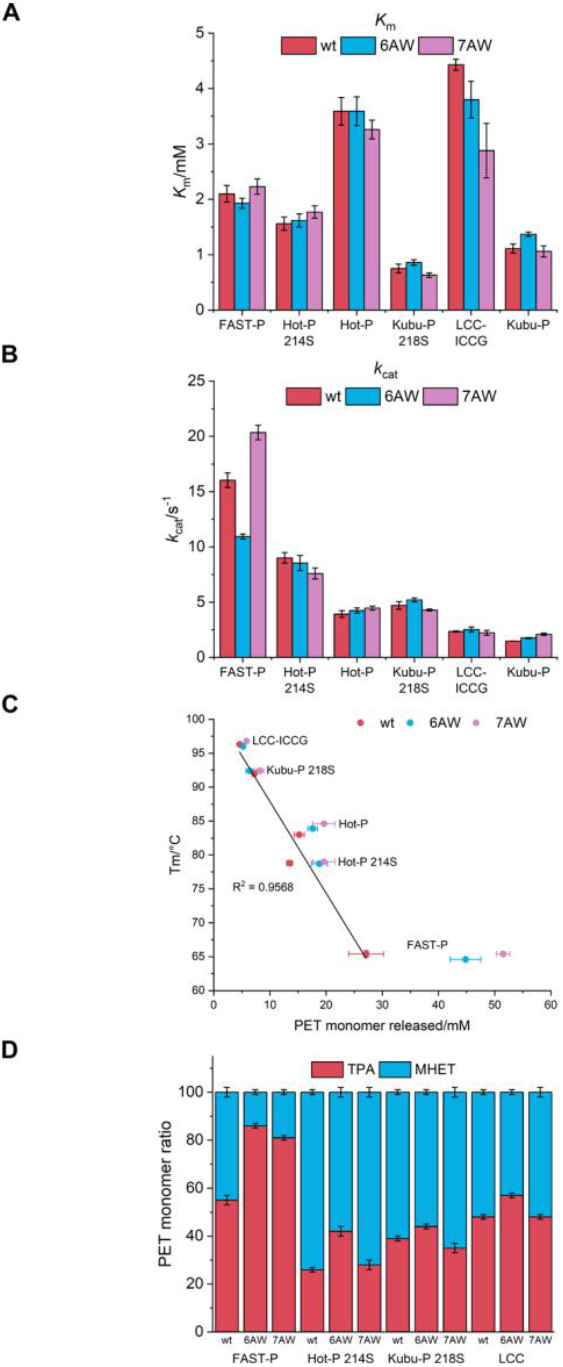
Activity–stability trade-off across PET hydrolase scaffolds and its modulation by azatryptophan substitution. (A) Michaelis constants (*K*_m_) and (B) catalytic turnover number (*k*_cat_) for the parent PET hydrolases and their azaPETases determined by **PETra** at 30 °C. (C) Correlation between melting temperature (Tm) and PET monomer release in 24 h (mM) for wild-type enzymes (red) or azatryptophan substitutions (6AW, blue; 7AW, purple). The black line shows the linear regression for wild-type enzymes. (D) Product profile (TPA versus MHET) during steady-state hydrolysis of amorphous gf-PET films at 50 °C (FAST-PETase, Kubu-PETase 218S, and LCC-ICCG) or 60 °C (Hot-PETase and its 214S mutant). The data represent the mean of three independent replicates ± standard deviation.

We also analyzed how replacing Trp185 with 6AW or 7AW affects product distribution of TPA versus MHET during steady-state PET hydrolysis (Figure 6D). For all PET hydrolases tested, 6AW increased the TPA fraction, with the strongest effect observed in FAST-PETase (86% TPA of the total aromatic products). In contrast, 7AW produced smaller shifts, except in FAST-PETase, where TPA reached 81%. This shift toward TPA is advantageous, as the accumulation of MHET requires additional downstream processing for conversion to TPA, increasing cost and complexity.^[45]^

The enhanced catalytic engagement of AW-substituted enzymes may arise from additional pH-dependent hydrogen-bonding interactions. Introducing a nitrogen atom into the indole ring creates a hydrogen-bond acceptor capable of interacting with carboxylic acid groups on the PET surface.^[46,47]^ Such interactions may improve substrate capture and orient PET for productive catalysis, particularly in scaffolds with intrinsically higher Trp185 mobility.^[41]^

## Conclusion

Enhancing enzyme activities by substituting ncAAs into substrate binding sites is particularly challenging, especially when the ncAA differs substantially from the amino acid it replaces, and to date has been successful only in isolated cases.^[48-52]^ Furthermore, the cost of chemical synthesis of these ncAAs made large scale applications unattractive. In contrast, the protein-centric isosterism introduced in the present work employs ncAAs that are readily made by biosynthesis from inexpensive precursors, readily installed by genetic encoding, and successful at enhancing the activities of an entire class of enzymes of industrial significance. Importantly, the azatryptophans afford single CH-group editing at functionally sensitive sites with maximal preservation of the steric and structural framework of the target proteins. In the context of PET hydrolases, this approach yields azaPETases that simultaneously enhance catalytic efficiency and thermal robustness, breaking a long-standing activity–stability trade-off.

Tryptophan plays central roles in the molecular recognition, catalysis, and electron transfer of many proteins, including carbohydrate-binding proteins and cytochrome P450 enzymes. The concept of isosteric amino acid substitution can thus be applied to many other systems. Furthermore, genetic encoding systems have become available also for a wide range of fluoroaromatic amino acids^[53-56]^ which, by delivering single-atom substitutions, open further opportunities for precision tuning of protein function. Combined with straightforward biosynthetic protocols of these ncAAs from inexpensive precursors, these advances establish outstanding potential for next-generation biocatalysts of expanded scope, efficiency, and design flexibility.

To encourage the uptake of this technology, the plasmids for the site-specific incorporations of AW isomers have been deposited at Addgene (Watertown, MA) to support distribution and applications: pRSF-G1(4AW)RS (plasmid #207617), pRSF-G1(5AW)RS (plasmid #207618), pRSF-G1(6AW)RS (plasmid #207619), and pRSF-G1(7AW)RS (plasmid #207516)

## Supporting Information

The authors have cited additional references within the Supporting Information.^[53,57-63]^

Detailed experimental procedures can be found in the Supporting Information.

## Acknowledgements

We thank Dr. Harpreet Vohra and Michael Devoy at the John Curtin School of Medical Research, Australian National University, for technical support on FACS experiments. Financial support by the Australian Research Council, including the Centre of Excellence for Innovations in Peptide and Protein Science (CE200100012) and Discovery Projects (DP230100079, DP260100191), is gratefully acknowledged.

## Data Availability Statement

The data that support the findings of this study are available from the corresponding author upon reasonable request.

## Entry for the Table of Contents

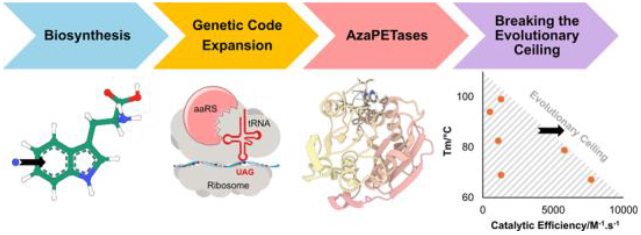

Coupling biosynthetic noncanonical amino acid production with genetic code expansion enables site-specific incorporation of azatryptophans into PET-degrading enzymes (PETases). By isosteric single-atom editing of a conserved tryptophan, AzaPETases break the activity–stability trade-off, delivering higher catalytic efficiency at elevated temperature and revealing new structure–activity relationships for enzyme design.

