## Supporting Information for "Isosteric Engineering of Enzymes: Overcoming Activity–Stability Trade-offs by Site-Selective CH → N Substitutions"

Table of Contents

Methods

a) Selection of functional G1PylRS enzymes recognising azatryptophans
b) Protein expression and purification
c) Multigram enzymatic synthesis of azatryptophans
d) Measuring PET hydrolase kinetics using PETra
e) PET hydrolase activity assay by degradation of amorphous Goodfellow PET film (gf-PET)

Figure S1: Enzymatic synthesis of azatryptophans using TrpB enzyme

Figure S2: 3D fluorescence emission spectra of azatryptophans

Figure S3: FACS experiments for selecting functional G1PylRS enzymes for 4AW

Figure S4: FACS experiments for selecting functional G1PylRS enzymes for 5AW

Figure S5: FACS experiments for selecting functional G1PylRS enzymes for 6AW

Figure S6: Confirmation of the site-specific, genetically encoded incorporation of azatryptophans into proteins

Figure S7: Assessment of background incorporation of azatryptophans at the tryptophan codon by the E. coli endogenous tryptophanyl-tRNA synthetase

Figure S8: Sequence alignment of the PET hydrolases investigated in this study

Figure S9: Fluorescence emission spectra of folded and unfolded wild-type and 4AW-substituted azaPETases

Figure S10: 1D 1H-NMR spectra of BHET, BHET-OH and their FAST-PETase hydrolysis products

Figure S11: 1D $^1$H-NMR kinetic analysis of the hydrolysis of BHET

Figure S12: Fluorescence characterisation of BHET-OH and its PET hydrolase hydrolysis products

Figure S13: Kinetic fluorescence spectra of BHET-OH hydrolysis by FAST-PETase

Figure S14: Michaelis–Menten analysis of FAST-PETase

Figure S15: Correlation between PETra-derived kinetic parameters and hydrolysis of amorphous gf-PET film

Figure S16: Analysis of PET degradation by Depo-PETase on gf-PET film

Figure S17: Reducing SDS-PAGE analysis of purified proteins following Ni–NTA affinity chromatography

Figure S18: Intact protein mass spectrometry analysis of purified proteins

Figure S19: Standardisation of gf-PET film by annealing for reproducible enzymatic degradation

Table S1: List of chemicals and their suppliers used in this study

Table S2: Photophysical properties of azatryptophan isomers

Table S3: Mutations in the G1PylRS variants selected for activity with 4AW, 5AW, and 6AW

Table S4: PETra-derived kinetic parameters for the PET hydrolases tested in this study

Table S5: Melting temperatures (Tm) of the PET hydrolases used in this study

Table S6: DNA and corresponding amino acid sequences of the proteins used in this study

References

### Methods

#### a) Selection of functional G1PylRS enzymes recognising azatryptophans

The selection of functional G1PylRS enzymes to specifically recognise 4AW, 5AW, and 6AW followed a previously established protocol, using the previously established library of G1PylRS mutants encoded on the pBK-G1RS plasmid.[53] The plasmid library was transformed into *E. coli* DH10B cells harboring the pBAD-H6RFP reporter plasmid encoding mRFP1 red fluorescent protein (RFP) with an amber stop codon following an N-terminal $His_6$ tag. After transformation, the culture was directly inoculated into 25 mL LB medium supplemented with 100 mg/L carbenicillin, 50 mg/L kanamycin, 0.4% L-arabinose, and 1 mM target ncAA. This culture served as the sample for the first round of positive selection (**1P+**). The same cells grown without ncAA supplementation were used as a control sample (**1P–**). Overnight expression at 37 °C led to readily detectable level of RFP expression, and then, the cells were harvested, resuspended in 5 mL of PBS buffer (137 mM NaCl, 2.7 mM KCl, 10 mM $Na_2HPO_4$, 1.8 mM $KH_2PO_4$, pH 7.4), and diluted 20-fold to a concentration suitable for fluorescence-activated cell sorting (FACS). FACS was performed using a FACSAria Fusion cell sorter (BD Biosciences, USA; Figures S1–3a). Cells with high red fluorescence levels were collected from the **1P+** sample (indicated by violet shades in Figures S1–3a) and subjected to a subsequent round of negative selection (**2N–**) in the absence of ncAA. Cells exhibiting low RFP expression were collected from the **2N–** sample and aliquoted to inoculate media under positive (**3P+**, with ncAA) and negative (**3P–**, without ncAA) conditions. The selection experiments continued with iterative positive and negative rounds following the same strategy.

The **5P+** sample for 4AW demonstrated a clear response to the presence of 4AW, and the top 3.1% of RFP-fluorescent cells from the **5P+** sample were collected. The selection experiments for 5AW and 6AW continued to the seventh round to reach enrichment on the target population. Approximately the top 3% (3.2% for 5AW and 3.9% for 6AW) of RFP-fluorescent cells from the **7P+** samples were collected.

About 2,000 cells collected from each final round were recovered by plating on LB agar plates containing 100 mg/L carbenicillin and 50 mg/L kanamycin. Isolated colonies were analysed in 96-well plates. Sixty enzyme candidates for each selection experiment were inoculated into media under both positive (with 1 mM ncAA) and negative (without ncAA) conditions. The red fluorescence intensity was measured as an indicator of RFP expression and normalised to the $OD_{600}$ of the cell culture using a TECAN Infinite 200 Pro M Plex plate reader (Tecan, Switzerland). DNA sequence analysis of the pBK-G1RS plasmids identified three, two, and two individually different candidates to incorporate 4AW, 5AW, and 6AW, respectively. The amino acid mutations of these candidates are listed in **Table S3**.

**b) Protein expression and purification**

**1. Optimisation of the expression of *Tm*9D8* TrpB**

The low expression yield of *Tm*9D8* TrpB (10 mg/L cell culture) prompted us to reengineer the construct to increase the protein yield. In the optimised protein *Tm*9D8* TrpB-opt, the $His_6$-tag was relocated from the N-terminus to the C-terminus. The expression vector carried kanamycin resistance (**Table S6**). *Tm*9D8* TrpB-opt was expressed in *E. coli* BL21(DE3) cells transformed with pET-24(+)*Tm*9D8* TrpB-opt (Twist Bioscience, USA). The transformed cells were grown at 37 °C in LB medium containing 50 mg/L kanamycin. A 10 mL aliquot of the overnight culture was used to inoculate 1 L LB medium supplemented with 50 mg/L kanamycin. Cultures were allowed to grow at 37 °C. At $OD_{600}$ of 0.6–1.0, the temperature was reduced to 25 °C, and protein expression was induced by the addition of 1 mM IPTG.

After 16 h, the cells were harvested by centrifugation (4,000 *g*, 4 °C, 15 min). Following resuspension in buffer A (50 mM Tris-HCl pH 7.5, 300 mM NaCl, 5% glycerol, 10 mM imidazole), 100 µM pyridoxal 5'-phosphate (PLP) was added, and the cells were lysed by sonication (ultrasonic homogeniser Omni-Ruptor 4000, Omni International, USA) on ice (50% power and 50% pulse length for 12 min). The cell lysate was then heated at 75 °C for 30 min in a water bath. The heat-treated lysate was clarified by centrifugation (30,000 *g*, 4 °C, 1 h). The clear supernatant was loaded onto a 5 mL Ni-NTA HisTrap column connected to an ÄKTA pure 25 chromatography system (Cytiva, USA). The column was washed with 20 column volumes buffer A and the protein was eluted with 5 column volumes buffer B (same as buffer A but with 500 mM imidazole). Afterwards, the buffer of the eluted protein was exchanged to 50 mM potassium phosphate buffer, pH 8.0, using an Amicon ultrafiltration centrifugal tube with a molecular weight cut-off of 10 kDa. The yield of purified *Tm*9D8* TrpB-opt was ~40 mg/L cell culture.

**2. Expression and purification of azaPETases**

*E. coli* B-95.ΔA cells[57] were co-transformed with the pRSF-G1 plasmid carrying the evolved azatryptophan G1PylRS/$tRNA^{CUA}$ pair and the pCDF plasmid containing the PET hydrolase gene, where the target Trp codon was replaced by a TAG codon (Twist Bioscience, USA; **Table S6**). The transformed cells were grown at 37 °C in LB medium containing 25 mg/L kanamycin and 25 mg/L spectinomycin. A 1 mL aliquot of the overnight culture was used to inoculate 100 mL LB medium supplemented with 25 mg/L kanamycin, 25 mg/L spectinomycin, and 1 mM of the corresponding azatryptophan. The cells were grown at 37 °C to an $OD_{600}$ of 0.6–1. At this point, the temperature was reduced to 25 °C and protein expression was induced by the addition of 1 mM IPTG.

Wild-type PET hydrolases were expressed in a similar way, but from cells transformed with the pCDF plasmid containing the wild-type gene (Twist Bioscience, USA) grown in LB medium supplemented with 25 mg/L spectinomycin.

After expression for 16 h, the cells were harvested by centrifugation (4,000 *g*, 4 °C, 15 min). Following resuspension in buffer A, the cells were lysed by sonication as described above. The cell lysate was centrifuged (30,000 *g*, 4 °C, 1 h) and the cleared supernatant was loaded onto a 1 mL His GraviTrap column (Cytiva, USA). The column was washed with 30 column volumes buffer A and the protein was eluted with 5 column volumes buffer B. Afterwards, the buffer was exchanged to PBS buffer using an Amicon ultrafiltration centrifugal tube (Merck Millipore, Germany) with a molecular weight cut-off of 10 kDa.

Protein concentrations were determined from absorbance measurement at 280 nm using extinction coefficients calculated with the Expasy server[58] and adjusted for each azatryptophan using the values listed in **Table S2**. The purity of the expressed enzymes was confirmed by SDS–PAGE (**Figure S17**) and the ncAA incorporation fidelity was verified by intact protein mass spectrometry (**Figure S18**).

#### 3. Expression, purification, and cleavage of NT* domain with C-terminal ENLYFQGDX motif

NT* domain with the C-terminal ENLYFQGDX motif (X standing for a non-canonical amino acid encoded by the amber stop codon) was produced as described for the PET hydrolases above. To release the GDX tripeptide (X = azatryptophan), 0.5 mL NT* fusion was incubated with $His_6$-TEV protease (10:1 by mass) at 25 °C for 4 h. Next, 0.5 mL Ni Sepharose 6 Fast Flow resin (Cytiva, USA) was added to capture the $His_6$-tagged proteins, leaving the GDX tripeptide in the supernatant, which was collected by centrifugation at 21,000 *g* for 5 min. This step was repeated three more times. $D_2O$ was added to the supernatant to a final concentration of 10% for analysis by 1D $^1$H-NMR.

#### 4. Intact protein mass spectrometry

Intact protein analysis was performed on an Orbitrap Fusion Tribrid mass spectrometer (Thermo Fisher Scientific, USA) connected to a Thermo Fisher Scientific UltiMate 3000 HPLC system equipped with ZORBAX 300SB-C3, 3.5 µm, 4.6 x 50 mm HPLC column (Agilent Technologies, USA). Approximately 50 pmol of sample was injected using a 500 µL/min linear gradient of solvent A (0.1% (v/v) formic acid in water) and solvent B (0.1% (v/v) formic acid in acetonitrile), ramping solvent B from 5% solvent B at the start to 80% after 12 min. Data were collected using an electrospray ionisation (ESI) source in positive ion mode. Protein intact mass was determined by deconvolution using the program Xcalibur 3.0.63 (Thermo Fisher Scientific, USA).

### c) Multigram enzymatic synthesis of azatryptophans

1 g L-serine (9.516 mmol) and 48 mg pyridoxal 5'-phosphate (0.2 mmol) were dissolved in 50 mL 100 mM potassium phosphate buffer, pH 8.0. The solution was heated to 55 °C. Next, 960 mg azaindole (8.126 mmol) dissolved in 2 mL ethanol was added. The

enzymatic reaction was initiated by the addition of *Tm*9D8* TrpB (20 μM final concentration) and the reaction volume adjusted to 60 mL with the reaction buffer.

The reaction mixture was shaken during incubation at 55 °C. To monitor the reaction progress, 20 μL samples of the reaction mixture were removed, diluted to 0.5 mL with PBS buffer, and analysed by 1D $^{1}$H-NMR (**Figure S1**). After completion, the reaction mixture was cooled to room temperature and the pH adjusted to 1 using concentrated HCl to ensure complete dissolution of the synthesised azatryptophan. Milli-Q water was added to adjust the volume to 80 mL to produce a 100 mM stock solution of the azatryptophan. This stock solution was used directly for the *in vivo* expression experiments without further processing.

**d) Measuring PET hydrolase kinetics using PETra**

The assay was performed in a 96-well black polystyrene plate, where 50 μL BHET-OH (from 100 mM stock in absolute ethanol) in 100 mM potassium phosphate buffer (pH 8.0) was added to column wells as 2-fold serial dilutions ranging from 4 mM to 0.031 mM. Subsequently, the reaction was started by adding 50 μL of 200 nM PET hydrolase solution prepared in 100 mM potassium phosphate buffer (pH 8.0) to each well. The plate was immediately placed in a TECAN Infinite 200 Pro M Plex plate reader (Tecan, Switzerland) adjusted to 30 °C to measure fluorescence kinetics ($\lambda_{ex}/\lambda_{em}$ = 400/480 nm) using kinetic cycles (**Figure S13**). For data analysis, calibration curves of BHET-OH were generated at each time point from control wells containing BHET-OH and buffer only (no enzyme), and these curves were used to calculate the remaining concentration of BHET-OH at every measured time point. Finally, enzyme kinetics were determined by calculating initial velocities ($V_0$) across the varying BHET-OH concentrations and fitting the data to the Michaelis–Menten equation (**Figure S14**) using OriginPro 2024 (64-bit) SR1 (OriginLab Corporation, MA, USA).

**e) PET hydrolase activity assay by degradation of amorphous Goodfellow PET film (gf-PET)**

**Standardisation of gf-PET film by annealing.** This is a critical step for assay consistency, as it removes the enthalpy relaxation caused by aging of the polymer, a major source of variability in degradation results.[59] Prior to the assay, gf-PET discs (6 mm diameter) were prepared from Goodfellow 252-144-75 sheets (0.25 mm thickness; Merck KGaA, Darmstadt, Germany) using a standard hole punch, washed once with 70% ethanol and three times with Milli-Q water, and stored in Milli-Q water. Immediately before use, the discs were transferred to1.5 mL low protein binding polypropylene microfuge tubes containing 900 μL of the assay buffer and annealed by heating the tubes in a shaking water bath (180 rpm) at 85 °C for 5 min and then quickly quenched in an ice-water bath for 15 min (**Figure S19**).[60]

Following the annealing step, 100 µL of a 2 µM PET hydrolase solution was added to each 1.5 mL microfuge tube containing the gf-PET disc and the assay buffer (100 mM potassium phosphate buffer (pH 8.0) for FAST-PETase or 50 mM glycine buffer (pH 9.2) for the other enzymes). The tubes were then incubated with shaking (180 rpm) in a water bath at the specified temperature. To measure the total PET monomer released using the spectrophotometric method, 5 µL samples were taken out of the reaction tubes at defined time points, centrifuged at 20,000 *g* for 5 min to remove insoluble particles and then $A_{260}$ was measured using a NanoDrop OneC UV-Vis spectrophotometer (Thermo Fisher Scientific Inc., Delaware, USA). To determine the product profile (ratio of BHET, MHET and TPA), 50 µL samples were taken out of the reaction tubes and mixed with 450 µL PBS buffer and 55 µL $D_2O$ and analysed by 1D $^1$H-NMR spectroscopy. The percentage of each monomer type was determined form the integration of the aromatic protons. The exact concentration of the PET monomer was calculated using the following equation:

$$PET\ monomer\ concentration\ (mM) = \frac{1000(A_{260})}{\left(\frac{\%\mathrm{BHET} * 6100}{100}\right) + \left(\frac{\%\mathrm{MHET} * 5400}{100}\right) + \left(\frac{\%\mathrm{TPA} * 4000}{100}\right)}$$

Where 6100, 54100, and 4000 are the molar extinction coefficients ($M^{-1}.cm^{-1}$) of BHET, MHET and TPA at 260 nm, respectively.

### Supplementary Figures

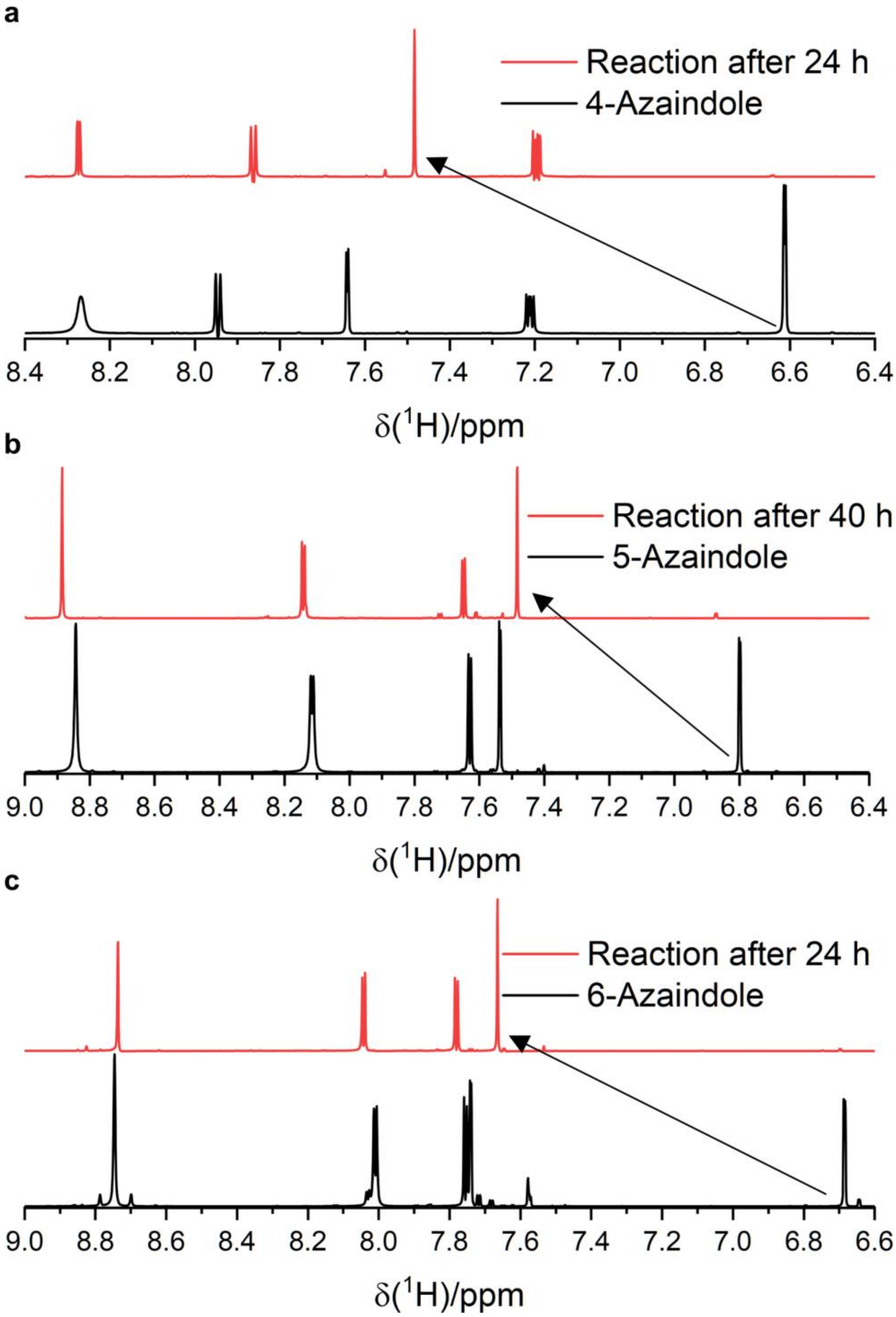

**Figure S1:** Enzymatic synthesis of azatryptophans using TrpB enzyme. **a-c,** Monitoring the progress of enzymatic synthesis of 4AW, 5AW, and 6AW by 1D $^{1}$H-NMR spectroscopy following incubation of serine (11 mM) and the respective azaindoles (10 mM) with *Tm*9D8* TrpB (20 µM) at 55 °C in 100 mM potassium phosphate buffer (pH 8.0). The spectral region shown comprises the NMR signals of the aromatic protons. Conversion yields were calculated by comparing the peak integrals of the H2 protons of the indole rings of the azaindole and corresponding azatryptophan (connected by an arrow). The conversion yields obtained are 98%, 95%, and 97% for 4AW, 5AW and 6AW, respectively.

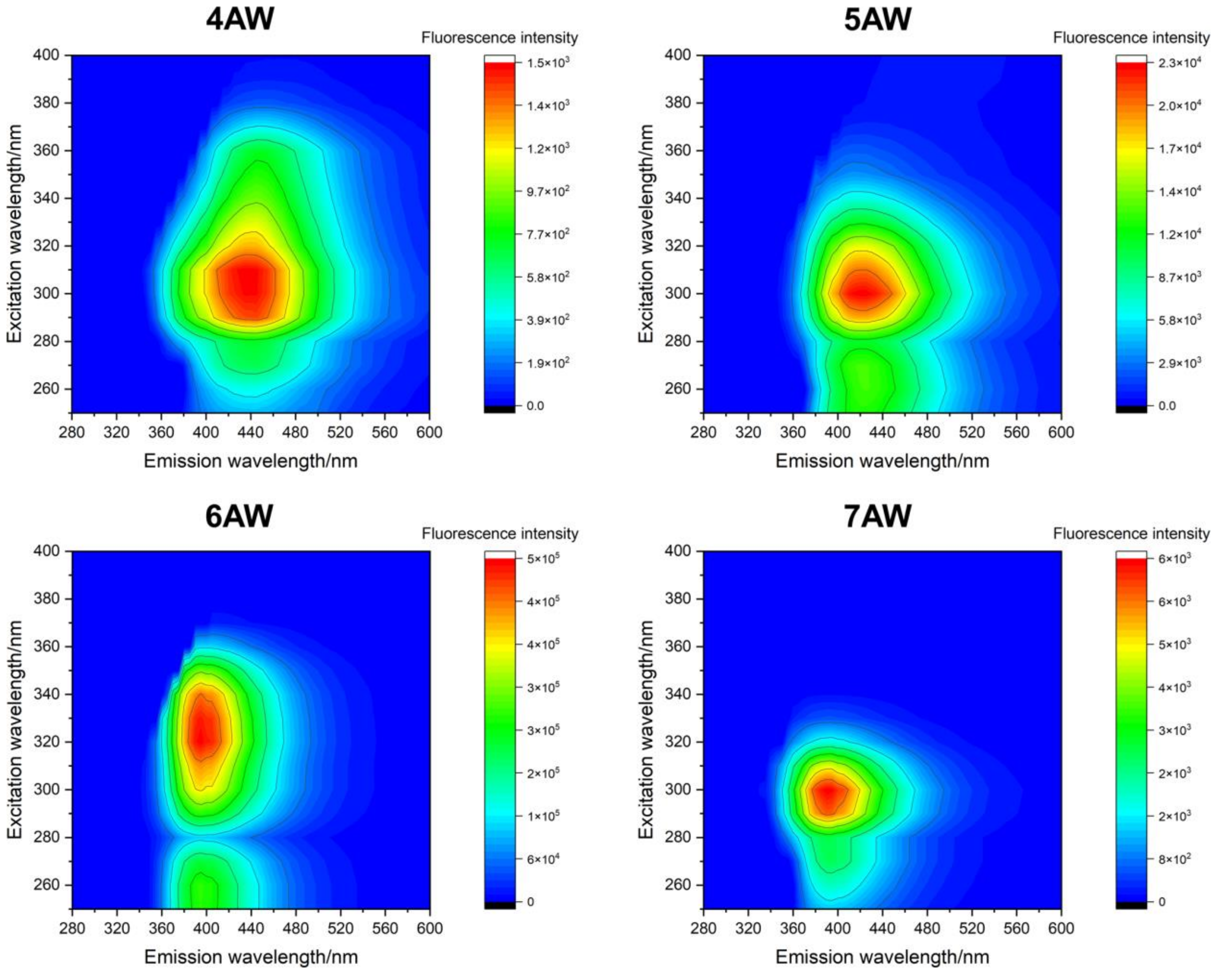

**Figure S2:** 3D fluorescence emission spectra of 10 μM aqueous solutions of azatryptophans in PBS buffer at 25 °C.

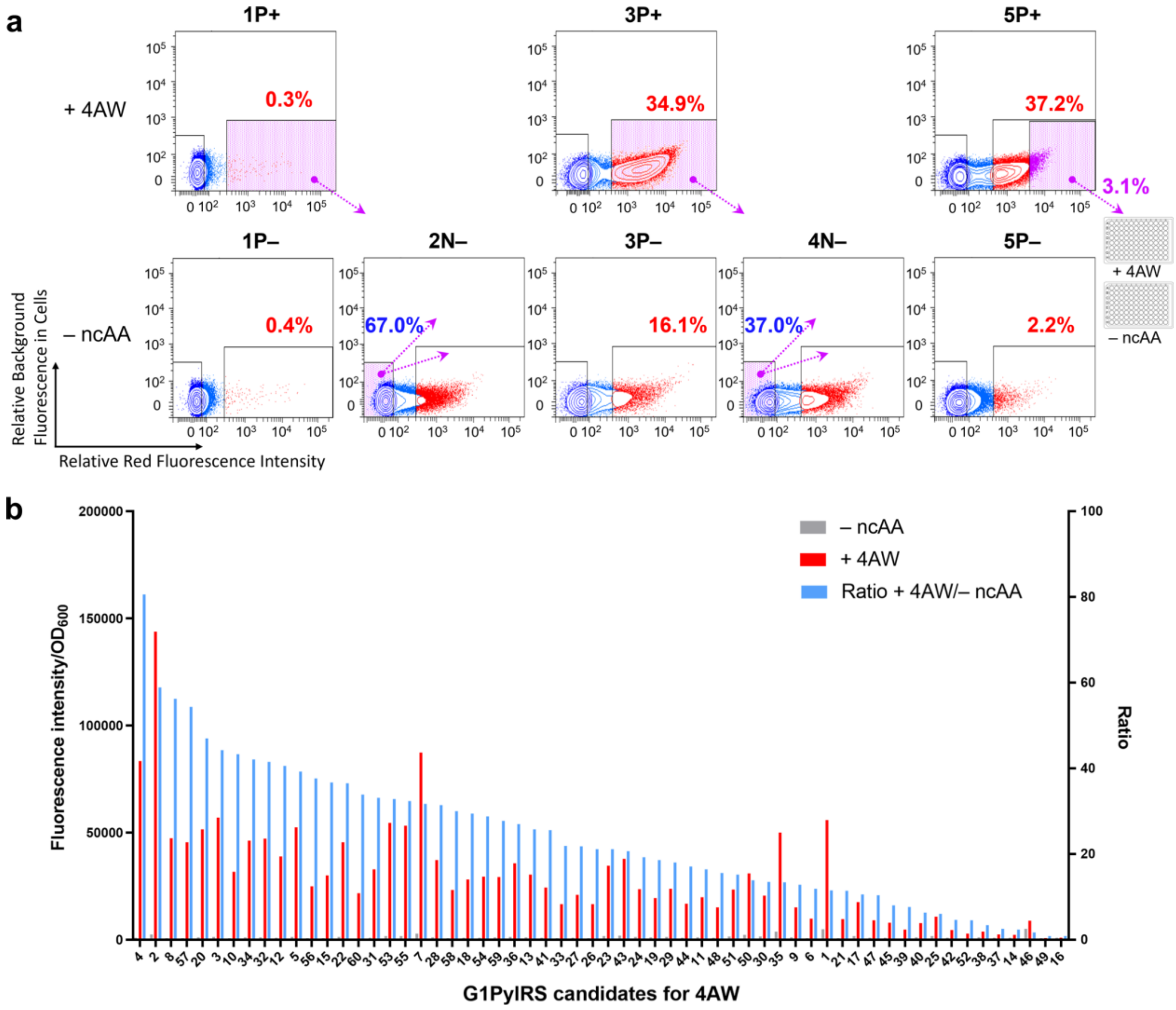

**Figure S3:** FACS experiments for selecting functional G1PylRS enzymes for 4AW. **a,** FACS screening of G1PylRS variants for activity and specificity in recognising 4AW. The horizontal axis of the scatter plots represents red fluorescence intensity (excitation at 560 nm), while the vertical axis indicates background fluorescence in cells excited at 488 nm. Violet-shaded regions identify the collected cell populations. Arrows illustrate the subsequent selection strategy applied after amplification by culturing. The preparation conditions of each sample are indicated with "+", denoting growth conditions with 1 mM ncAA, and "–" for conditions without ncAA. Positive and negative selection rounds are labelled as "**P**" and "**N**". **b,** Activity and specificity screen of G1PylRS variants for 4AW incorporation. Cells from the 3.1% fraction with the highest red fluorescence in the final selection round were cultured in 96-well plates with and without 1 mM 4AW. Red fluorescence intensity indicative of the readthrough efficiency of the amber-interrupted reporter gene was then measured. The plot presents the colonies ranked in descending order based on the ratio of red fluorescence in the + ncAA wells compared to the – ncAA wells. This ranking highlights the candidates with the highest activity and specificity for 4AW incorporation.

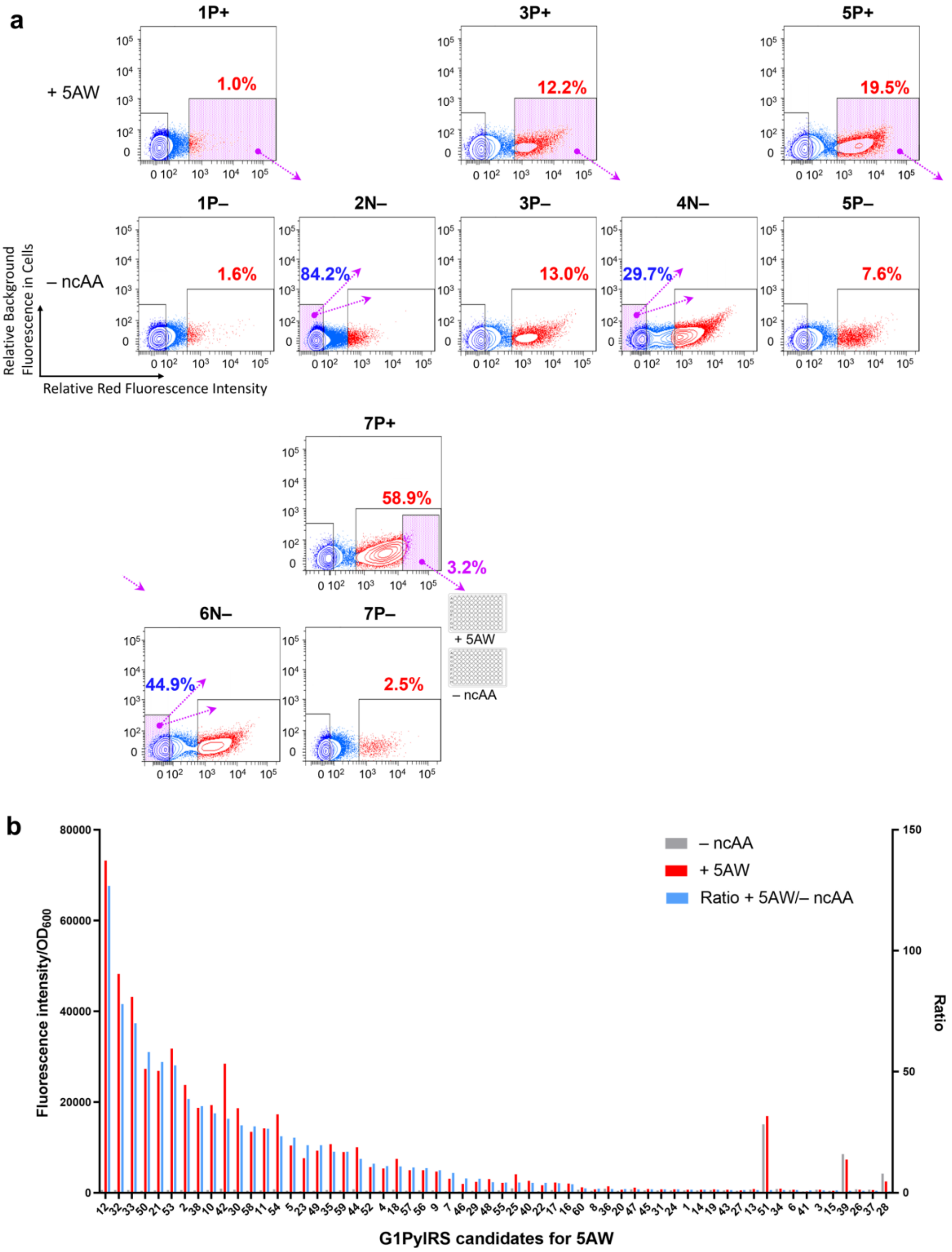

**Figure S4:** FACS experiments for selecting functional G1PylRS enzymes for 5AW. **a,** FACS screening of G1PylRS variants for activity and specificity in recognising 5AW. Annotations are the same as in **Figure S3**. **b,** Activity and specificity screen of G1PylRS variants for 5AW incorporation. Cells from the 3.2% fraction with the highest red fluorescence in the final selection round were cultured in 96-well plates with and without 1 mM 5AW. As in **Figure S3**, the plot presents the colonies ranked in a descending order based on the ratio of red fluorescence in the + ncAA wells compared to the – ncAA wells, highlighting the candidates with the highest activity and specificity for 5AW incorporation.

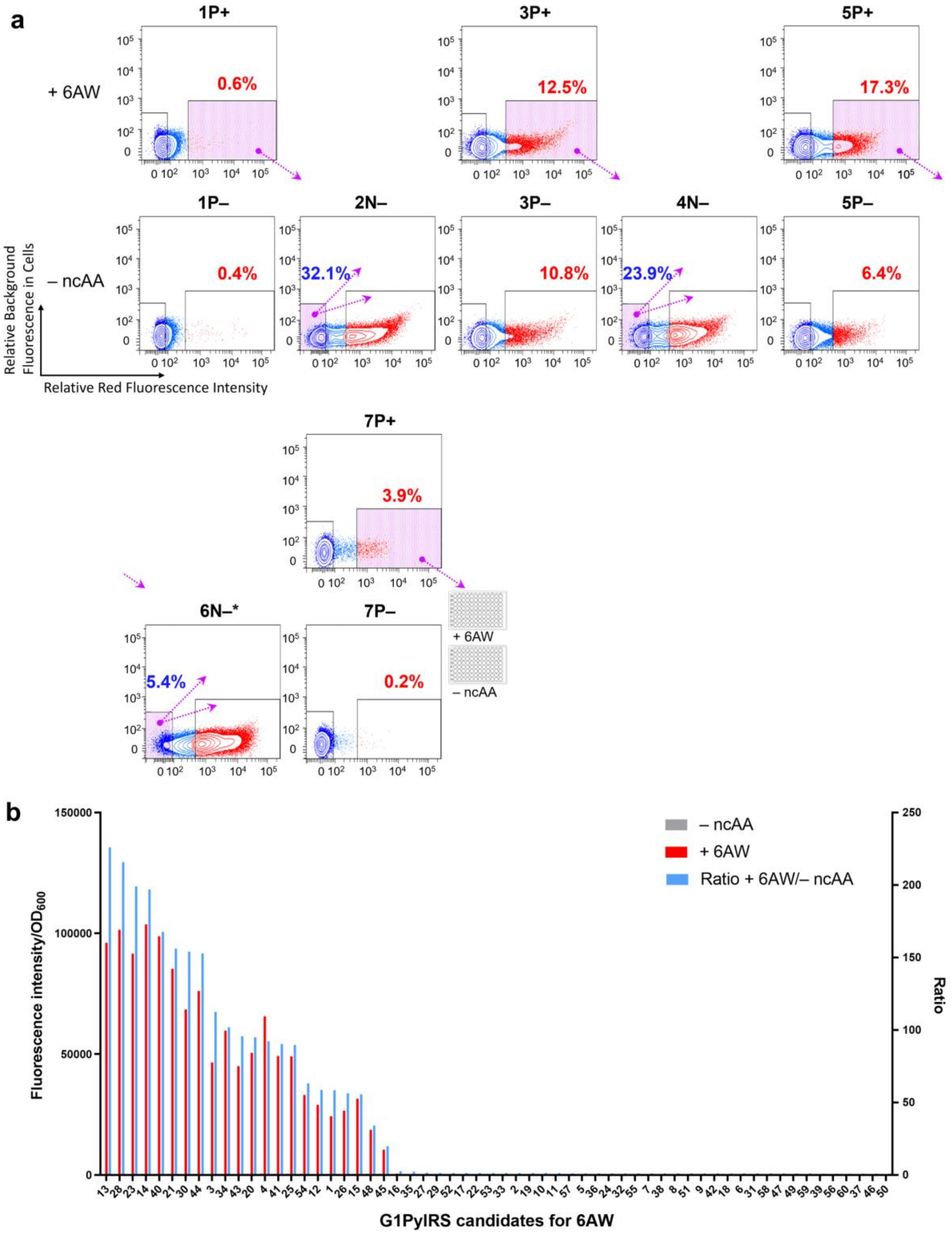

**Figure S5:** FACS experiments for selecting functional G1PylRS enzymes for 6AW. **a,** FACS screening of G1PylRS variants for activity and specificity in recognising 6AW. Annotations are the same as in **Figure S3**. **b,** Activity and specificity screen of G1PylRS variants for 6AW incorporation. Cells from the 3.9% fraction with the highest red fluorescence in the final selection round were cultured in 96-well plates with and without 1 mM 6AW. As in **Figure S3**, the plot presents the colonies ranked in a descending order based on the ratio of red fluorescence in the + ncAA wells compared to the – ncAA wells, highlighting the candidates with the highest activity and specificity for 6AW incorporation.

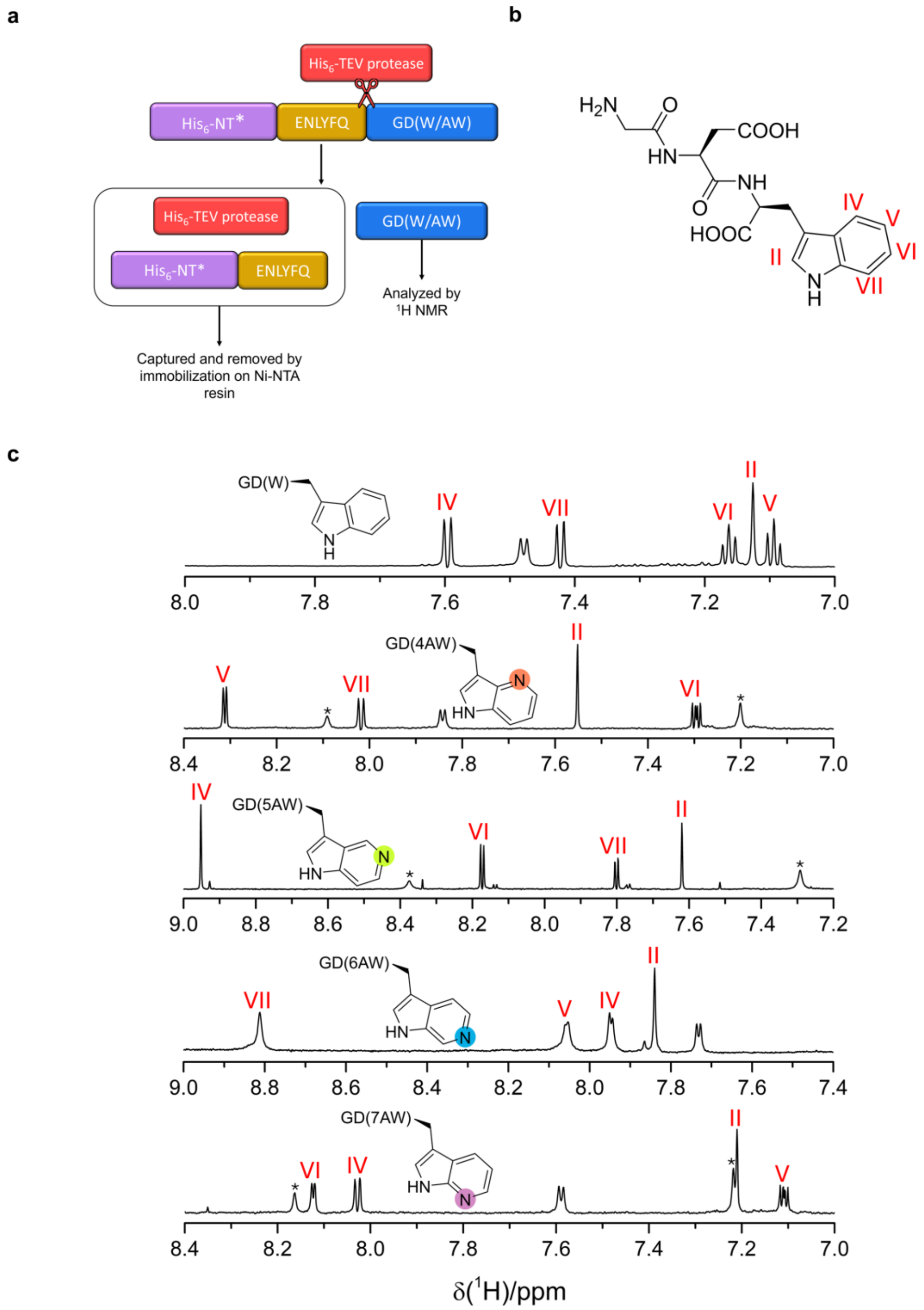

**Figure S6: Confirmation of the site-specific, genetically encoded incorporation of azatryptophans into proteins. a,** Diagram of the NT*-ENLYFQGDX construct used to produce the GD(W) or GD(AW) tripeptides and the approach for their isolation. **b,** Chemical structure of the GD(Trp) tripeptide, including numbering of the indole positions. **c,** 1D $^1$H-NMR spectra of GD(Trp) and the four different GD(AW) tripeptides recorded of 0.5–0.75 mM solutions in PBS buffer containing 10% $D_2O$. The spectra were recorded using a Bruker 800 MHz NMR spectrometer at 25 °C. Asterisks identify peaks of residual imidazole remaining after Ni–NTA purification.

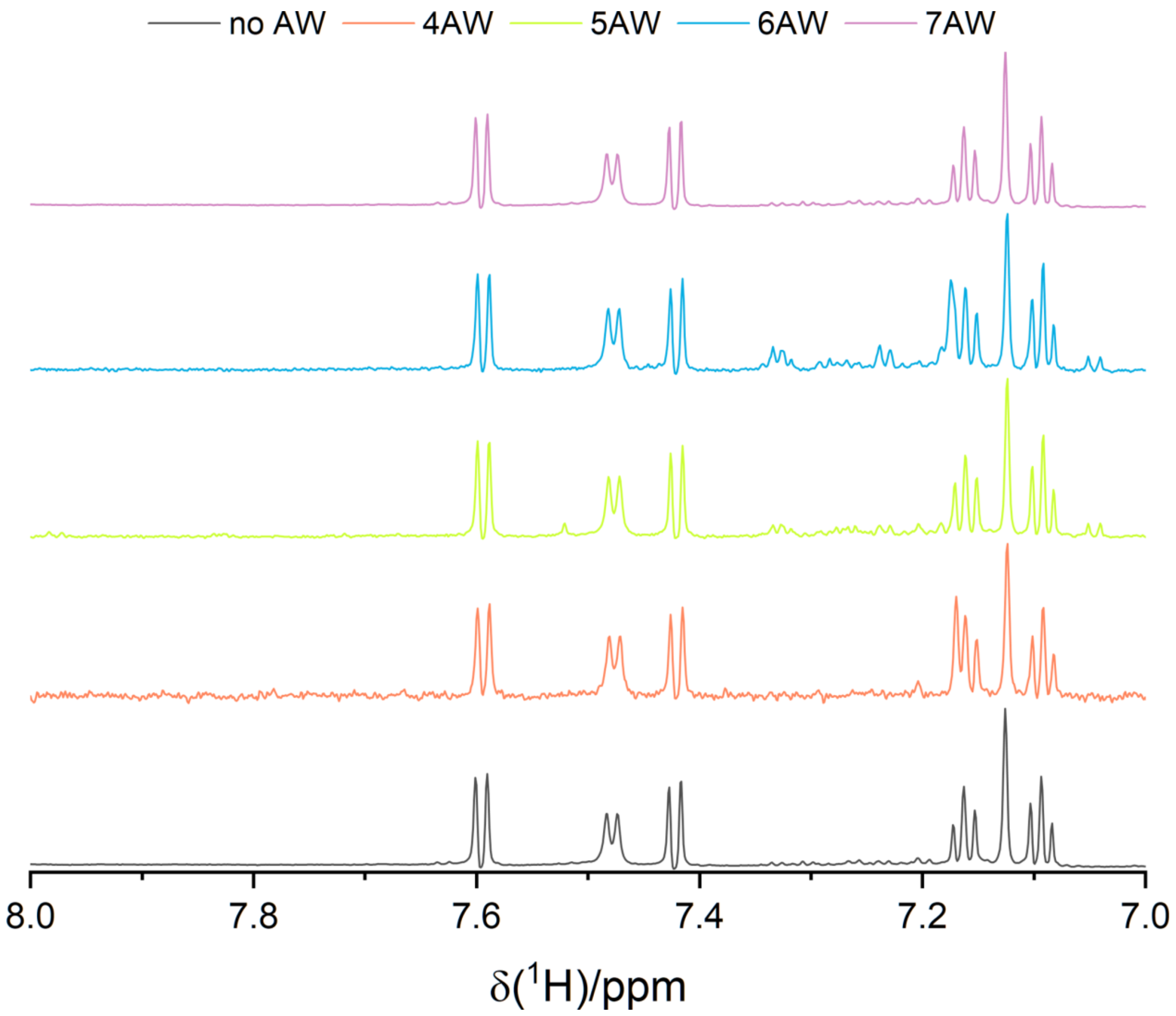

**Figure S7: Assessment of background incorporation of azatryptophans at the tryptophan codon by the *E. coli* endogenous tryptophanyl-tRNA synthetase.** The tripeptide GD(W) was purified from cultures supplemented with 1 mM azatryptophan and compared with GD(W) purified from media without azatryptophan supplementation. 1D $^{1}$H-NMR spectra of the GD(W) tripeptides show no evidence of azatryptophan incorporation at the tryptophan codon.

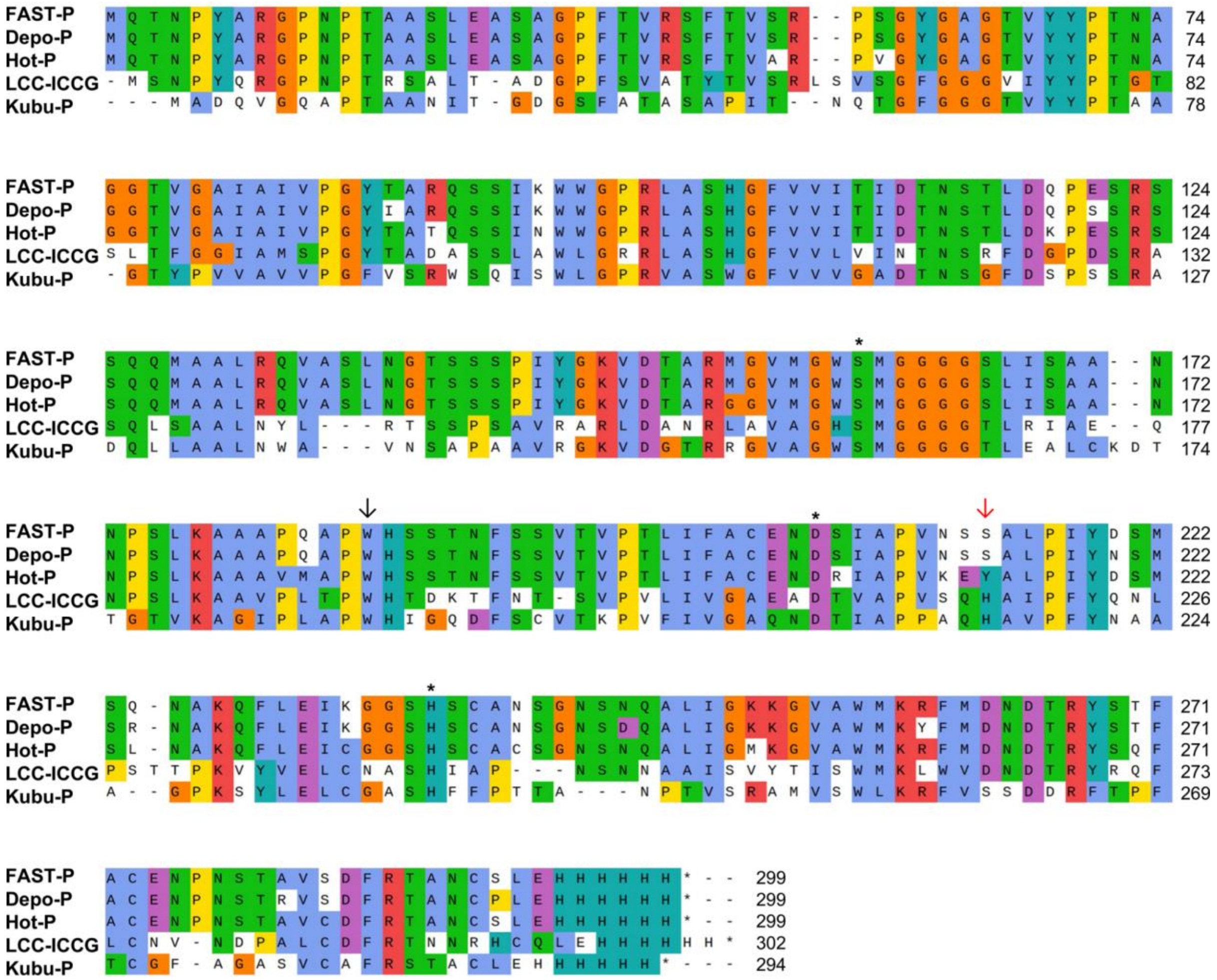

**Figure S8: Sequence alignment of the PET hydrolases investigated in this study using Clustal Omega 1.2.4 algorithm in SnapGene software (www.snapgene.com).** The wobbling tryptophan and the residue acting as the conformational latch are highlighted by black and red arrows, respectively. The catalytic Ser-Asp-His triad residues are denoted with the asterisks.

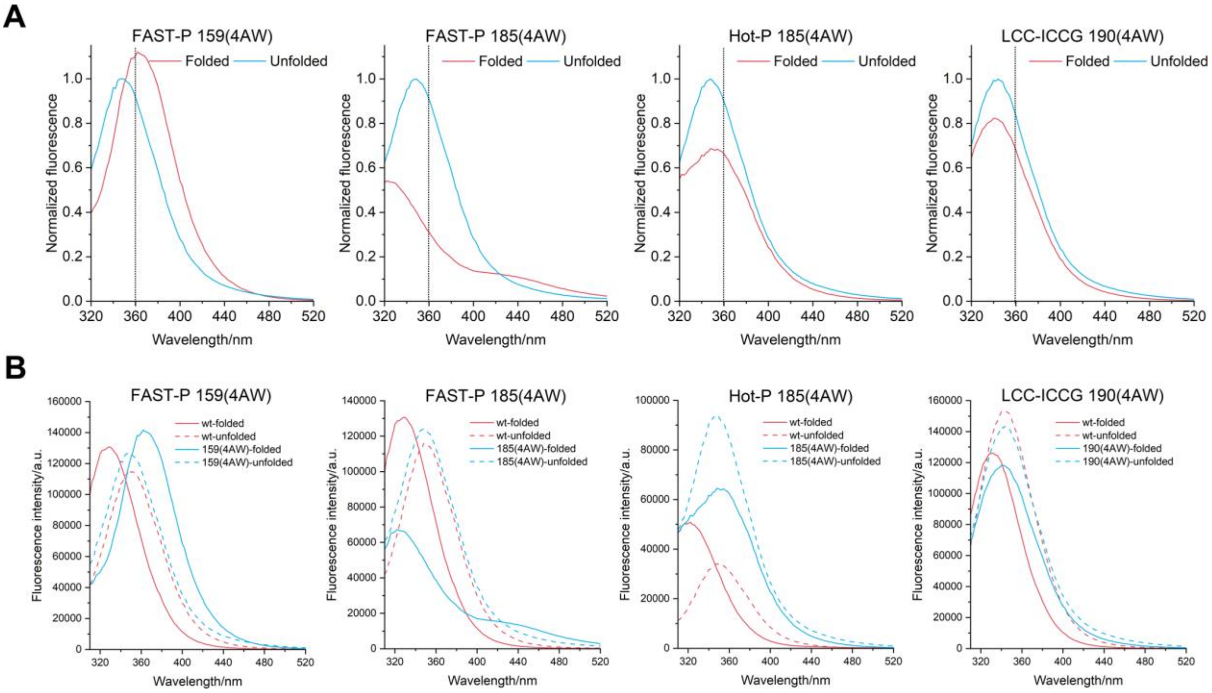

**Figure S9:** Fluorescence emission spectra of folded and unfolded wild-type and 4AW-substituted azaPETases excited at 280 nm. The data show that the fluorescence of 4AW also reports on solvent exposure more sensitively than tryptophan (compare the solvent-accessible mutant 159(4AW) with the buried 185(4AW) mutant of FAST-PETase) but the fluorescence of the four remaining tryptophan residues in the protein is dominant. In LCC-ICCG, 4AW at the wobbling tryptophan position produces a clear change in emission intensity and $\lambda_{em}$ (340 nm versus 330 nm in the wild-type), indicating reduced solvent exposure relative to FAST-PETase. In Hot-PETase, 4AW at position 185 yielded $\lambda_{em}$ = 360 nm, indicating relatively limited solvent exposure.

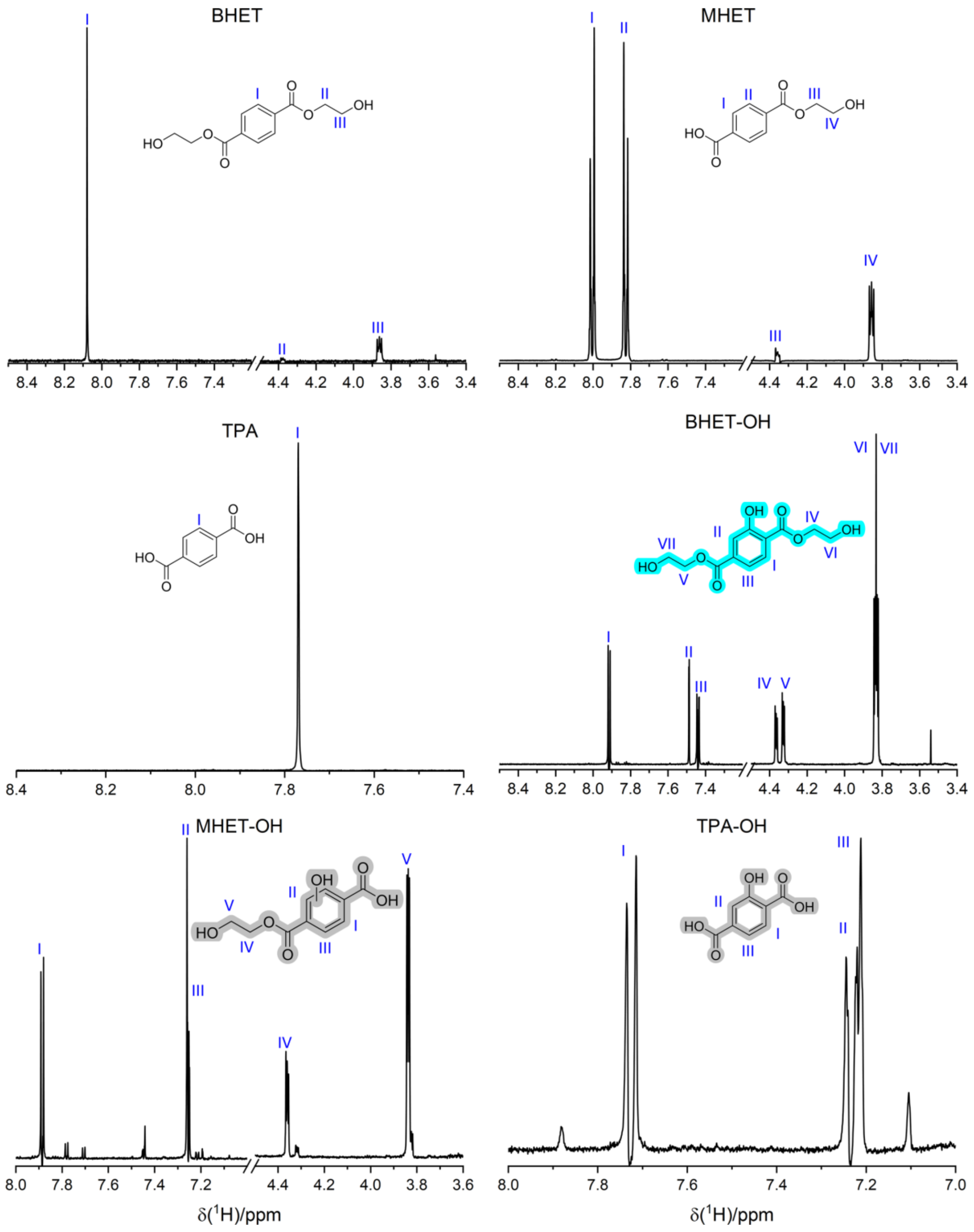

**Figure S10: 1D $^1$H-NMR spectra of BHET, BHET-OH and their FAST-PETase hydrolysis products.** Data were recorded on either a Bruker 400 MHz NMR spectrometer with a room-temperature probe or a Bruker 800 MHz NMR spectrometer with a cryoprobe at 25 °C, using a double spin-echo for solvent suppression. Samples (1–4 mM) were prepared in 100 mM potassium phosphate buffer (pH 8.0) containing 10% $D_2O$. Spectra of MHET-OH and TPA-OH were obtained by incubating 1 mM BHET-OH with 500 nM and 10 μM FAST-PETase, respectively, for 1 h at 25 °C. Roman numerals indicate the proton assignments.

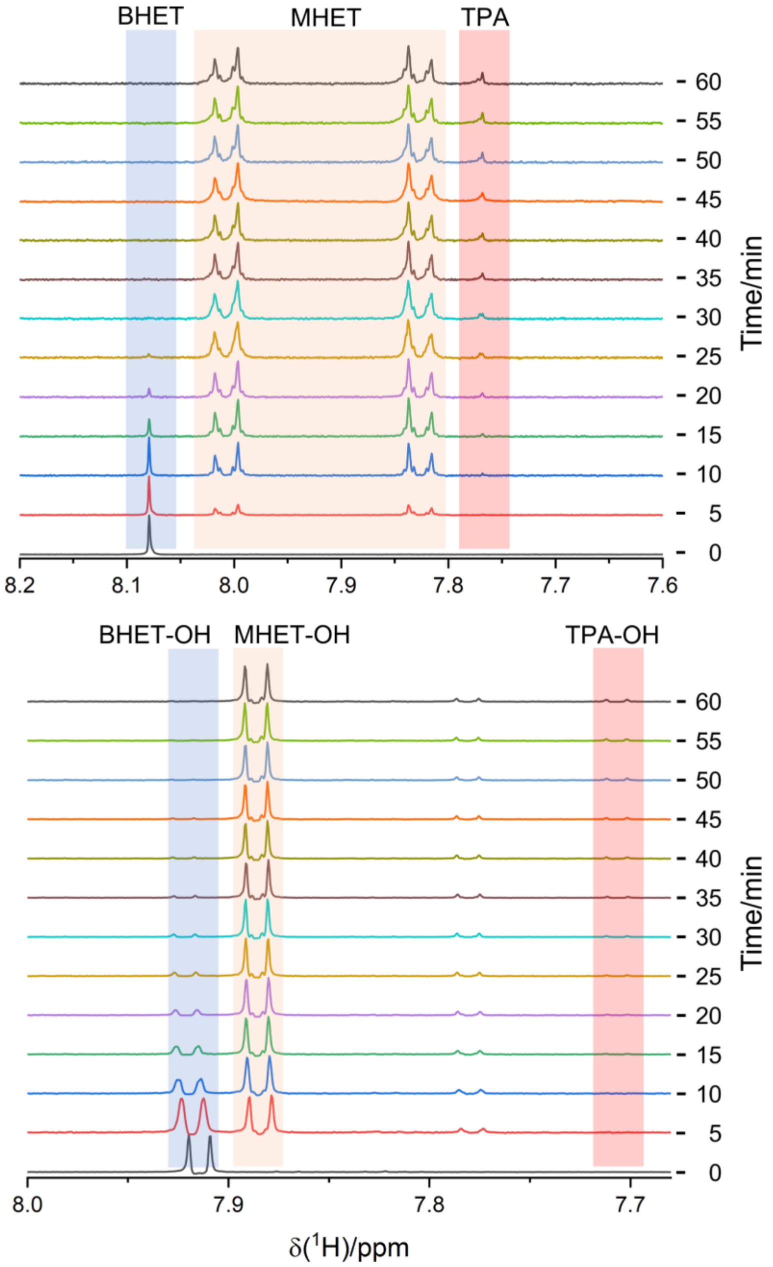

**Figure S11:** 1D $^{1}$H-NMR kinetic analysis of the hydrolysis of 1 mM BHET (recorded on a Bruker 400 MHz NMR spectrometer) and 1 mM BHET-OH (recorded on a Bruker 800 MHz NMR spectrometer) by 500 nM FAST-PETase at 25 °C, using a double spin-echo for solvent suppression. Samples were prepared in 100 mM potassium phosphate buffer (pH 8.0) containing 10% $D_2O$, and reactions were initiated by adding FAST-PETase directly to the 5 mm NMR tubes. One-minute spectra were acquired at the indicated time points, showing near-complete consumption of BHET and BHET-OH within 30 min.

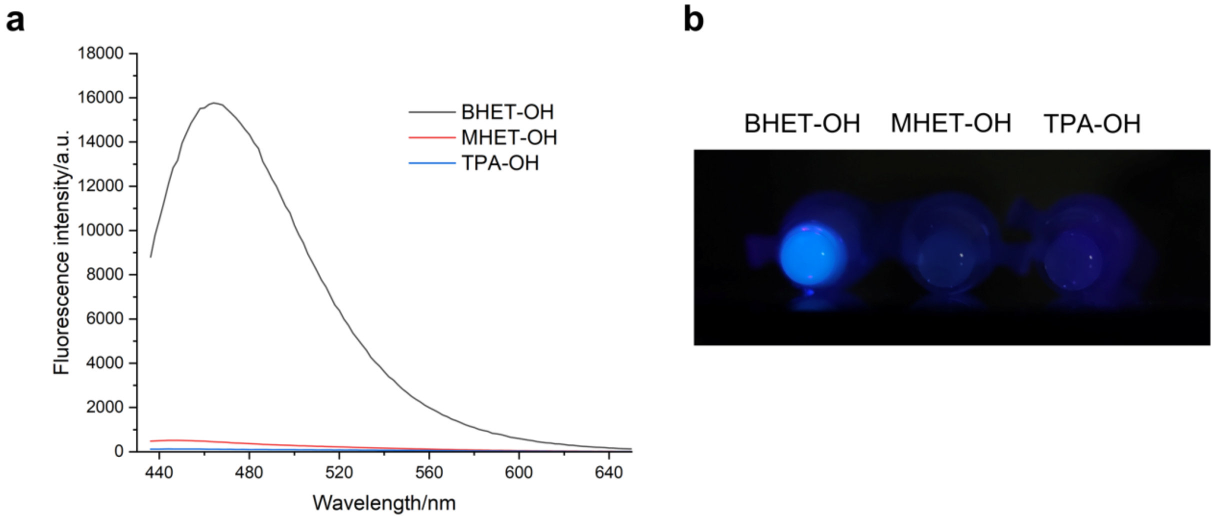

**Figure S12:** Fluorescence characterisation of BHET-OH and its PET hydrolase hydrolysis products. **a,** Fluorescence emission spectra ($\lambda_{ex}$ = 400 nm) of 1 mM BHET-OH alone (black) or after incubation with FAST-PETase at 500 nM (red) or 10 µM (blue) for 1 h at 25 °C, showing loss of fluorescence upon enzymatic hydrolysis. **b,** Visible fluorescence of the same samples shown in panel **a**, photographed with a mobile phone camera under 390 nm LED illumination. All samples were prepared in 100 mM potassium phosphate buffer (pH 8.0), and fluorescence spectra were recorded using a 96-well plate reader at 25 °C.

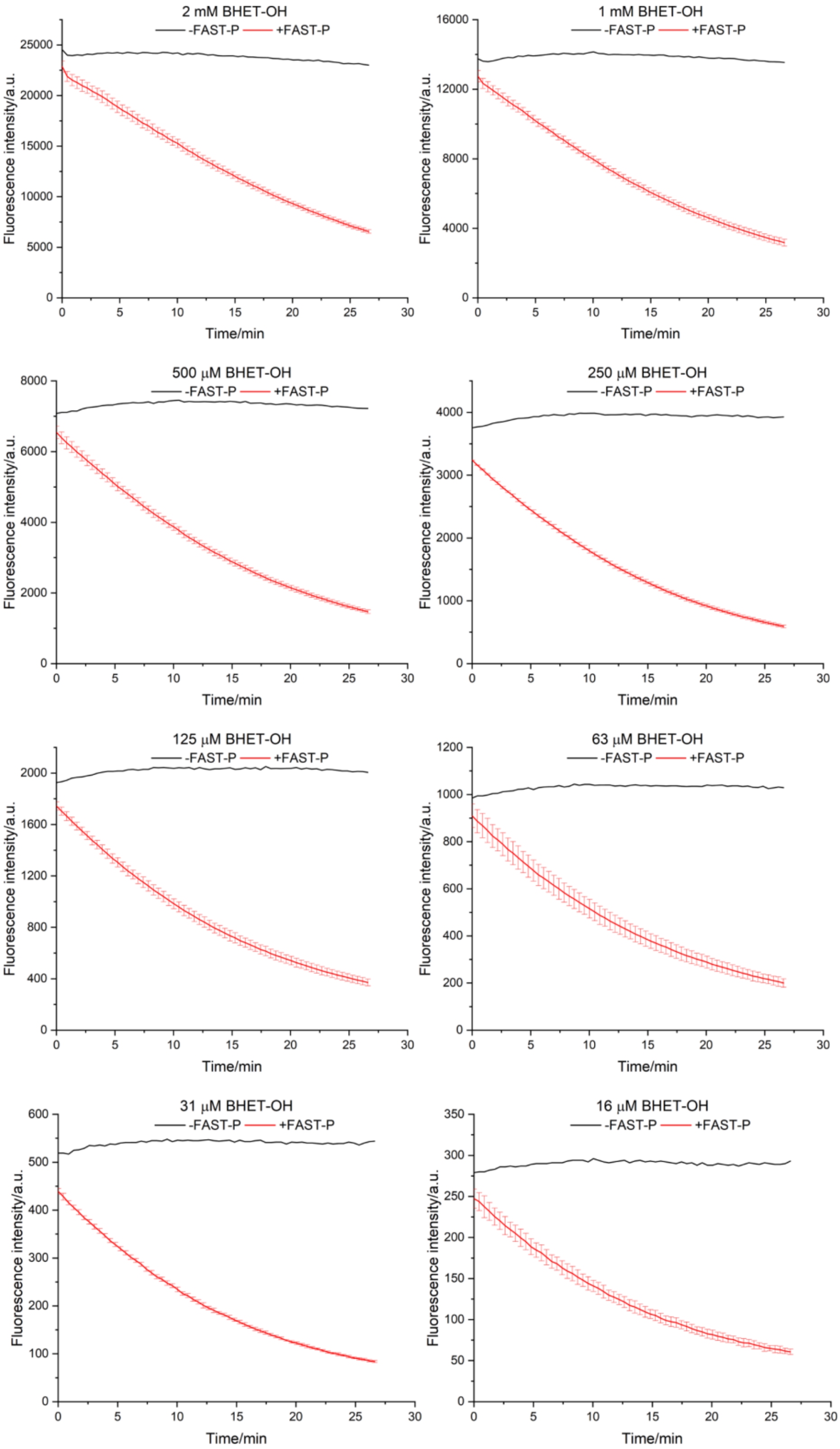

**Figure S13:** Kinetic fluorescence spectra of BHET-OH hydrolysis by FAST-PETase. Fluorescence traces of BHET-OH (2–0.016 mM) incubated with 100 nM FAST-PETase (red) measured at $\lambda_{ex}/\lambda_{em}$ = 400/480 nm. Black traces represent BHET-OH controls without enzyme. All reactions were performed in 100 mM potassium phosphate buffer (pH 8.0) at 30 °C, and fluorescence was recorded using a 96-well plate reader. Data represent the mean of three independent replicates ± standard deviation.

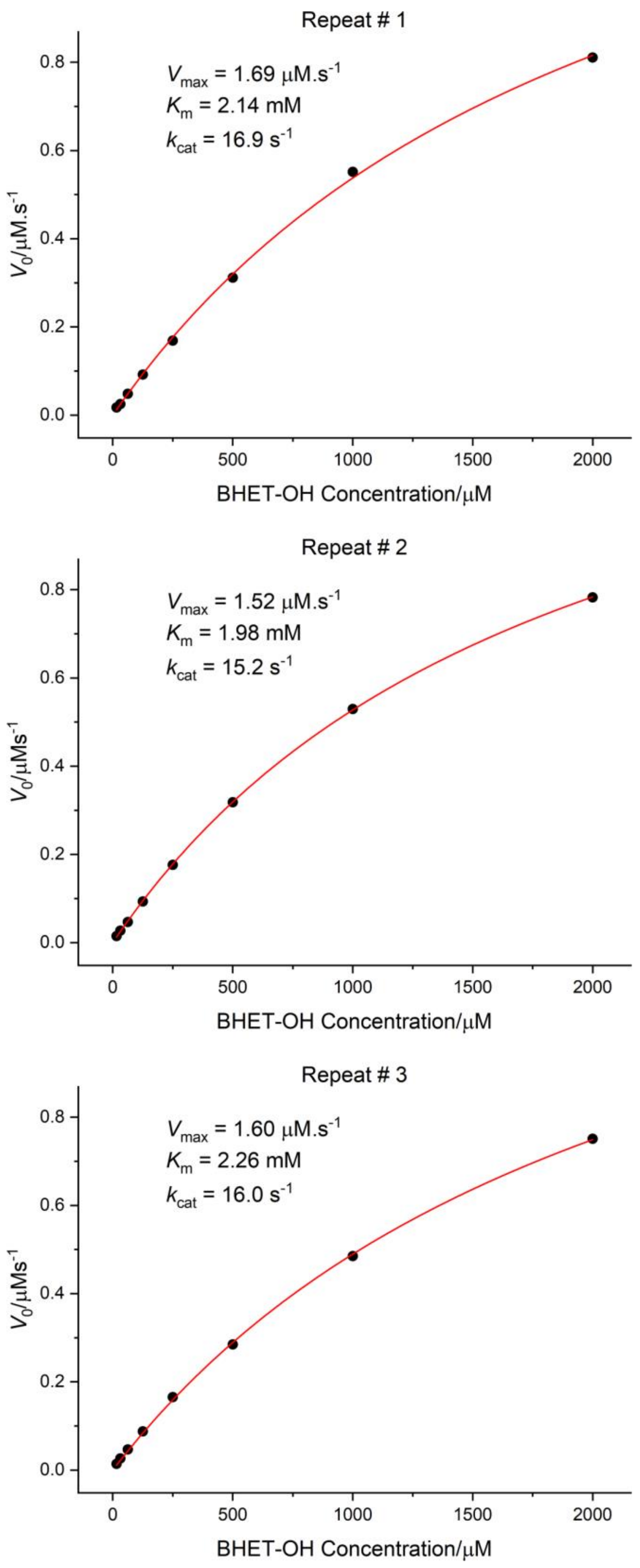

**Figure S14:** Michaelis–Menten analysis of FAST-PETase. Michaelis–Menten plots derived from the fluorescence kinetic data shown in **Figure S7**. Initial velocities were obtained by converting fluorescence changes into substrate concentrations using the BHET-OH calibration curve and fitting the data for each replicate individually to the Michaelis–Menten equation. The plots illustrate variability among replicates and the consistency of the derived kinetic parameters

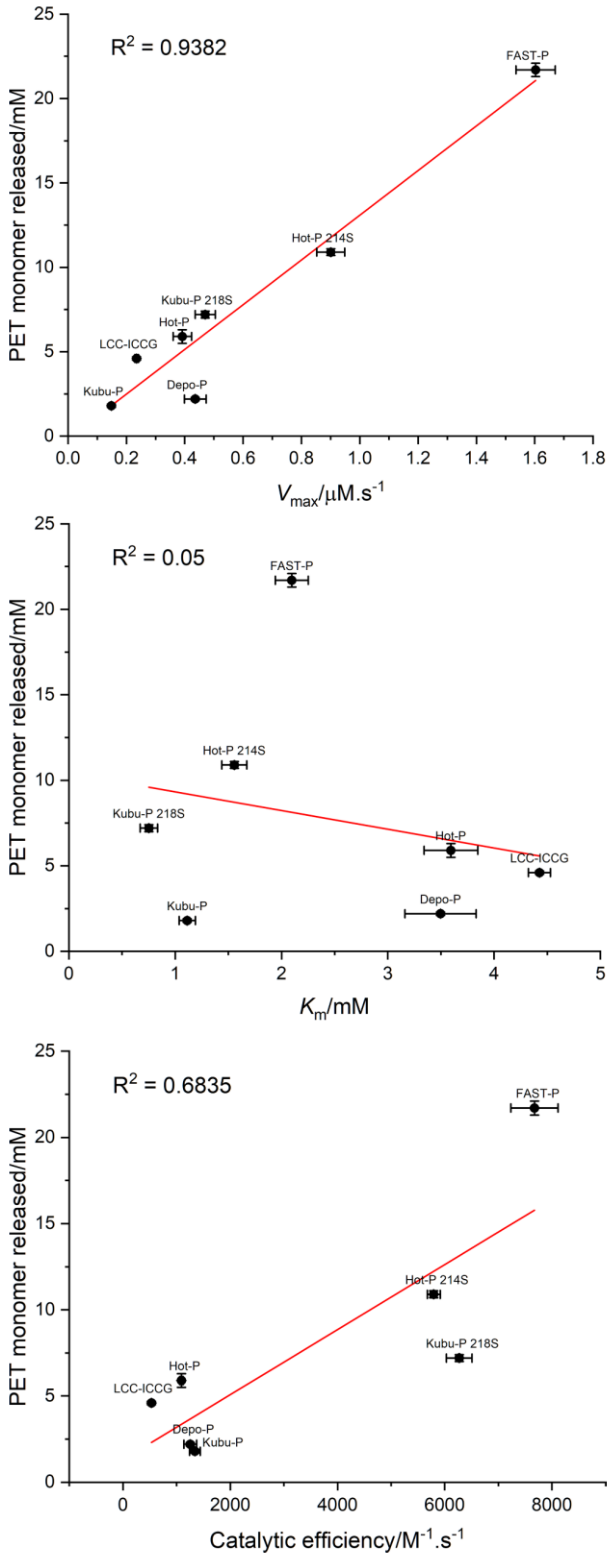

**Figure S15: Correlation between PETra-derived kinetic parameters and hydrolysis of amorphous gf-PET film.** The amount of PET monomers released after 24 h shows a strong positive correlation with $V_{max}$, and thus $k_{cat}$, given the fixed enzyme concentration of 100 nM ($R^2 = 0.94$), and a moderate correlation with catalytic efficiency ($k_{cat}/K_m$, $R^2 = 0.68$). By contrast, no meaningful correlation was observed with $K_m$ alone ($R^2 = 0.05$). Error bars represent the standard deviation of three independent replicates.

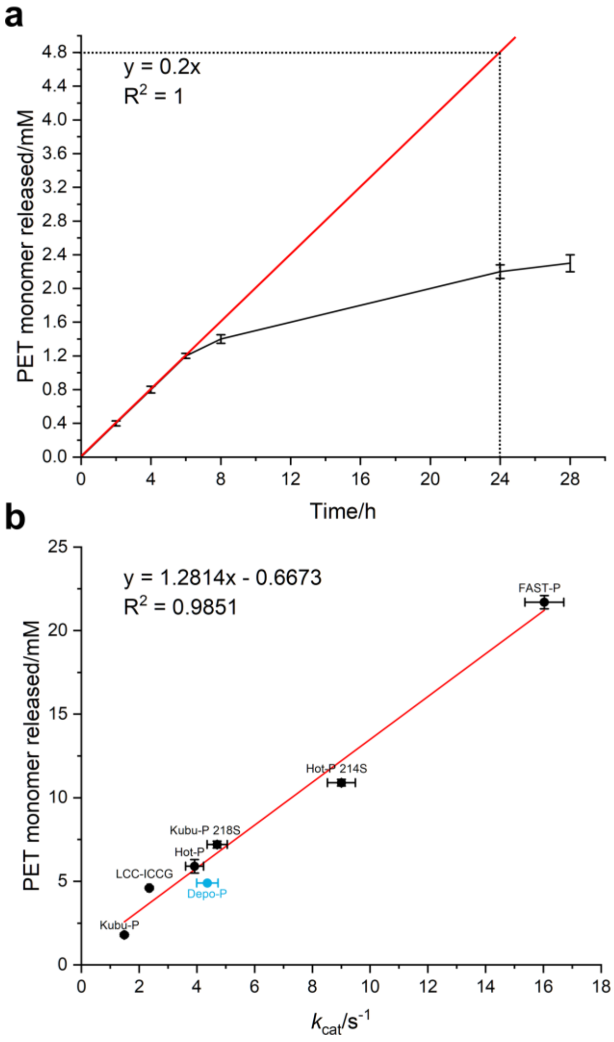

**Figure S16: Analysis of PET degradation by Depo-PETase on gf-PET film at 50 °C. a,** Extrapolation of PET monomer concentration to the 24 h time point based on the first three data points. The regression equation (y = 0.2x, $R^2$ = 1) indicates 4.8 mM monomer released in 24 h. **b,** Correlation between the PET monomers released from gf-PET film after 24 h at 50 °C and the **PETra**-derived $k_{cat}$ values at 30 °C. The correlation was calculated using PET hydrolases other than Depo-PETase (black points). The linear regression (y = 1.2814x - 0.6673, $R^2$ = 0.9851) predicts 4.9 mM PET monomers released for Depo-PETase (blue point), in close agreement with the estimated value in panel **a**. Error bars indicate the standard deviation of three independent replicates.

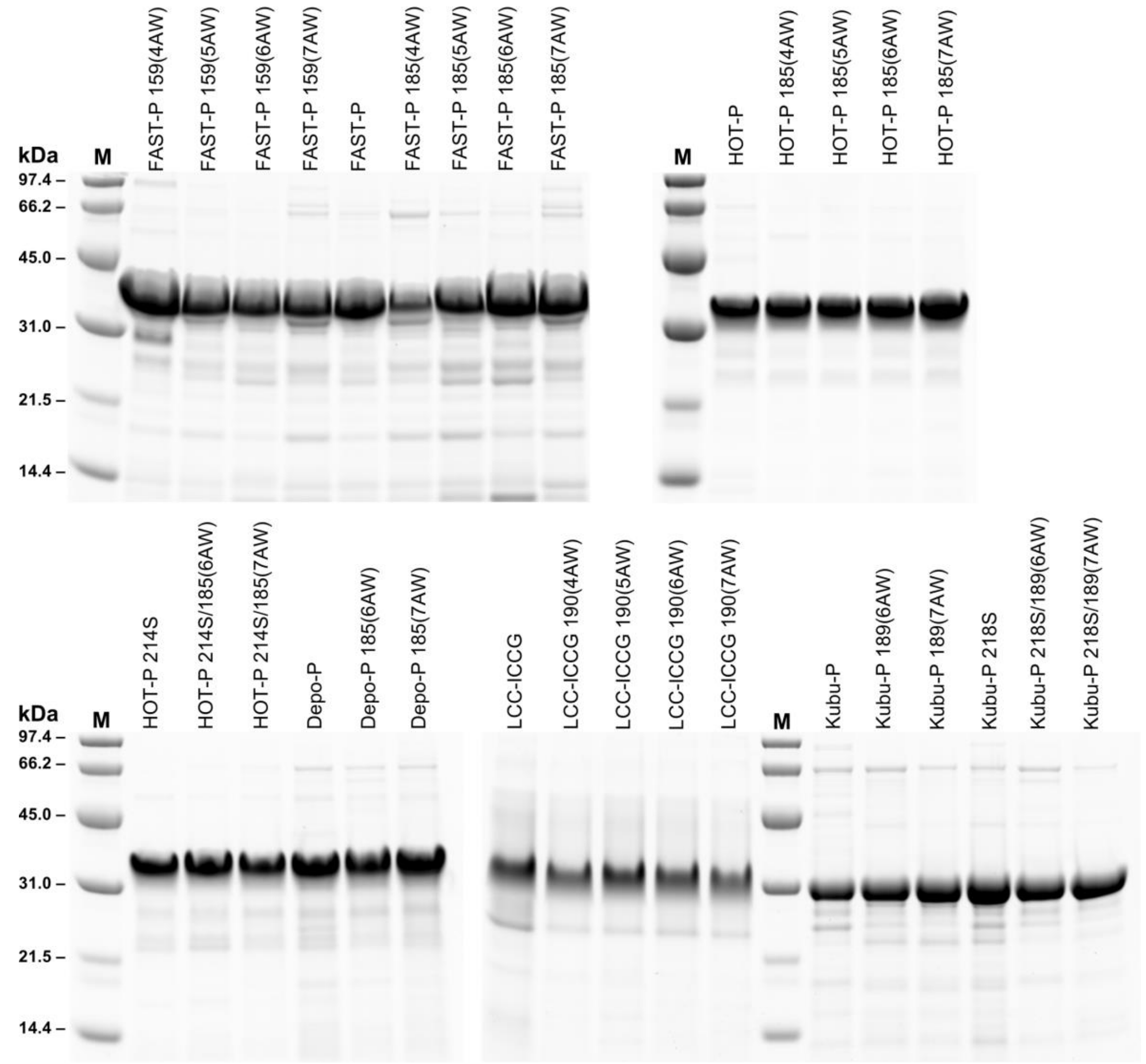

**Figure S17:** Reducing SDS-PAGE analysis of purified proteins following Ni–NTA affinity chromatography. Lane M contains molecular weight markers.

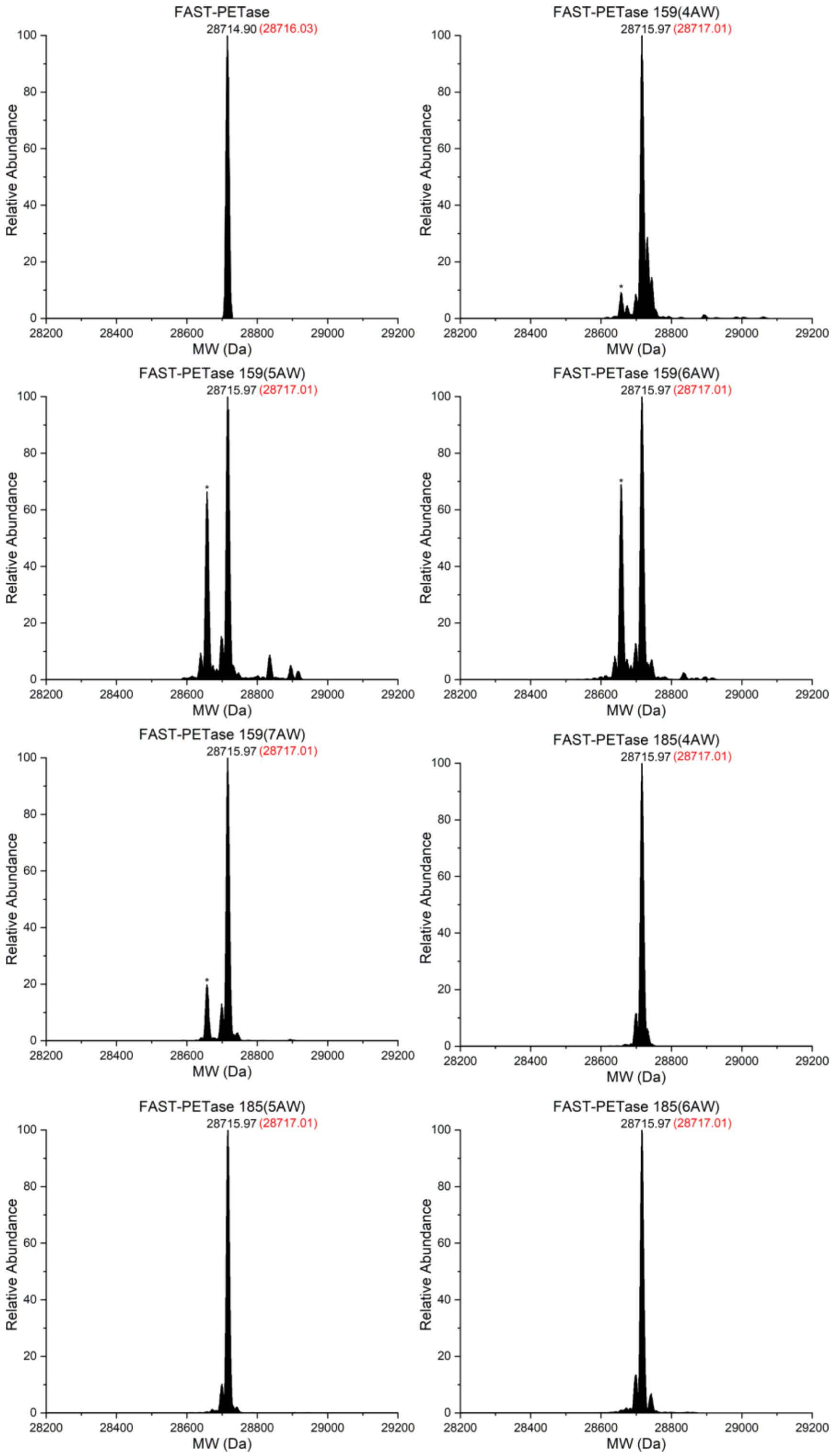
FAST-PETase
28714.90 (28716.03)
FAST-PETase 159(4AW)
28715.97 (28717.01)
FAST-PETase 159(5AW)
28715.97 (28717.01)
FAST-PETase 159(6AW)
28715.97 (28717.01)
FAST-PETase 159(7AW)
28715.97 (28717.01)
FAST-PETase 185(4AW)
28715.97 (28717.01)
FAST-PETase 185(5AW)
28715.97 (28717.01)
FAST-PETase 185(6AW)
28715.97 (28717.01)
Relative Abundance
MW (Da)
0
20
40
60
80
100
28200
28400
28600
28800
29000
29200
*

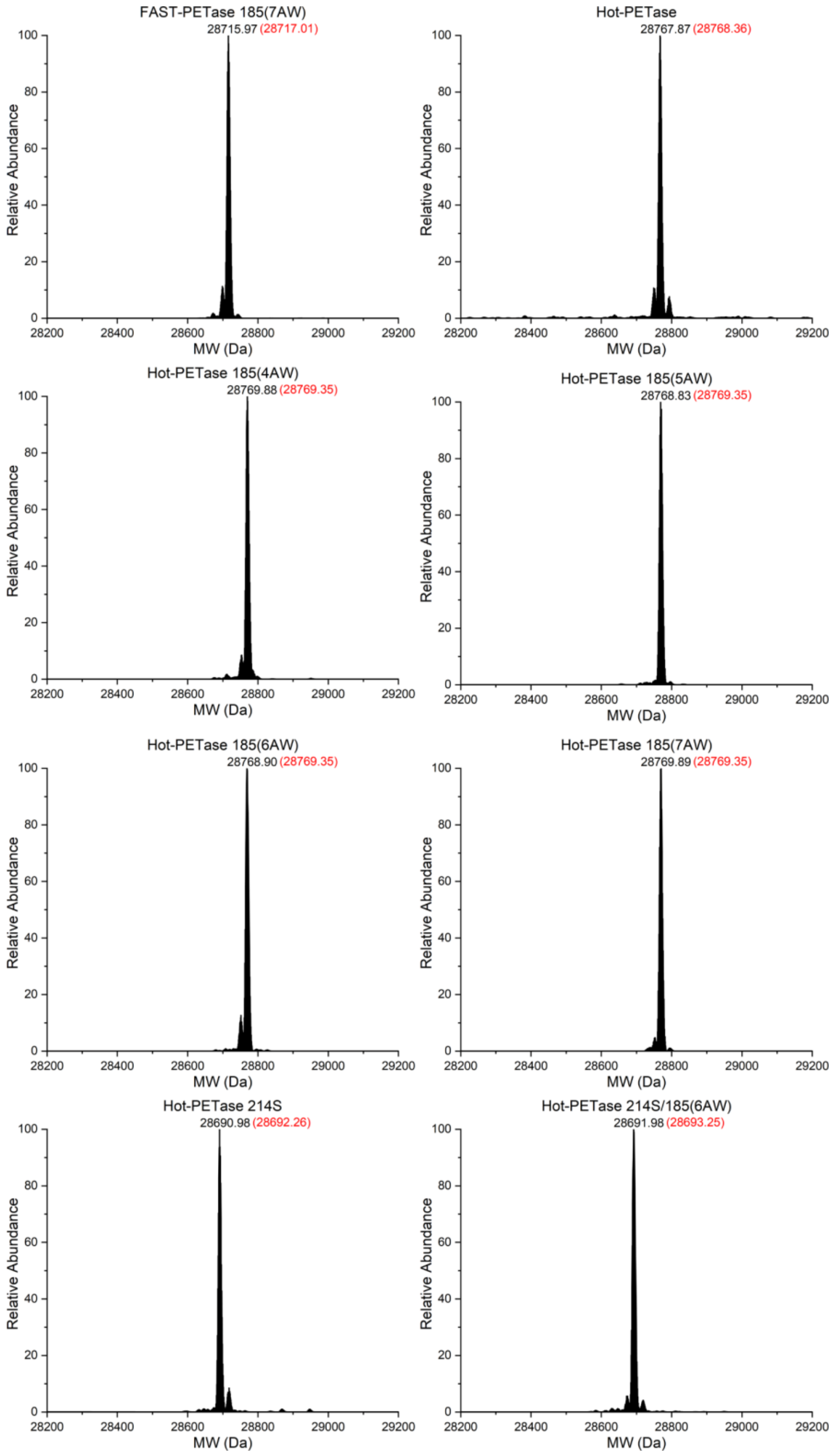
FAST-PETase 185(7AW)
28715.97 (28717.01)
Hot-PETase
28767.87 (28768.36)
Hot-PETase 185(4AW)
28769.88 (28769.35)
Hot-PETase 185(5AW)
28768.83 (28769.35)
Hot-PETase 185(6AW)
28768.90 (28769.35)
Hot-PETase 185(7AW)
28769.89 (28769.35)
Hot-PETase 214S
28690.98 (28692.26)
Hot-PETase 214S/185(6AW)
28691.98 (28693.25)
Relative Abundance
MW (Da)

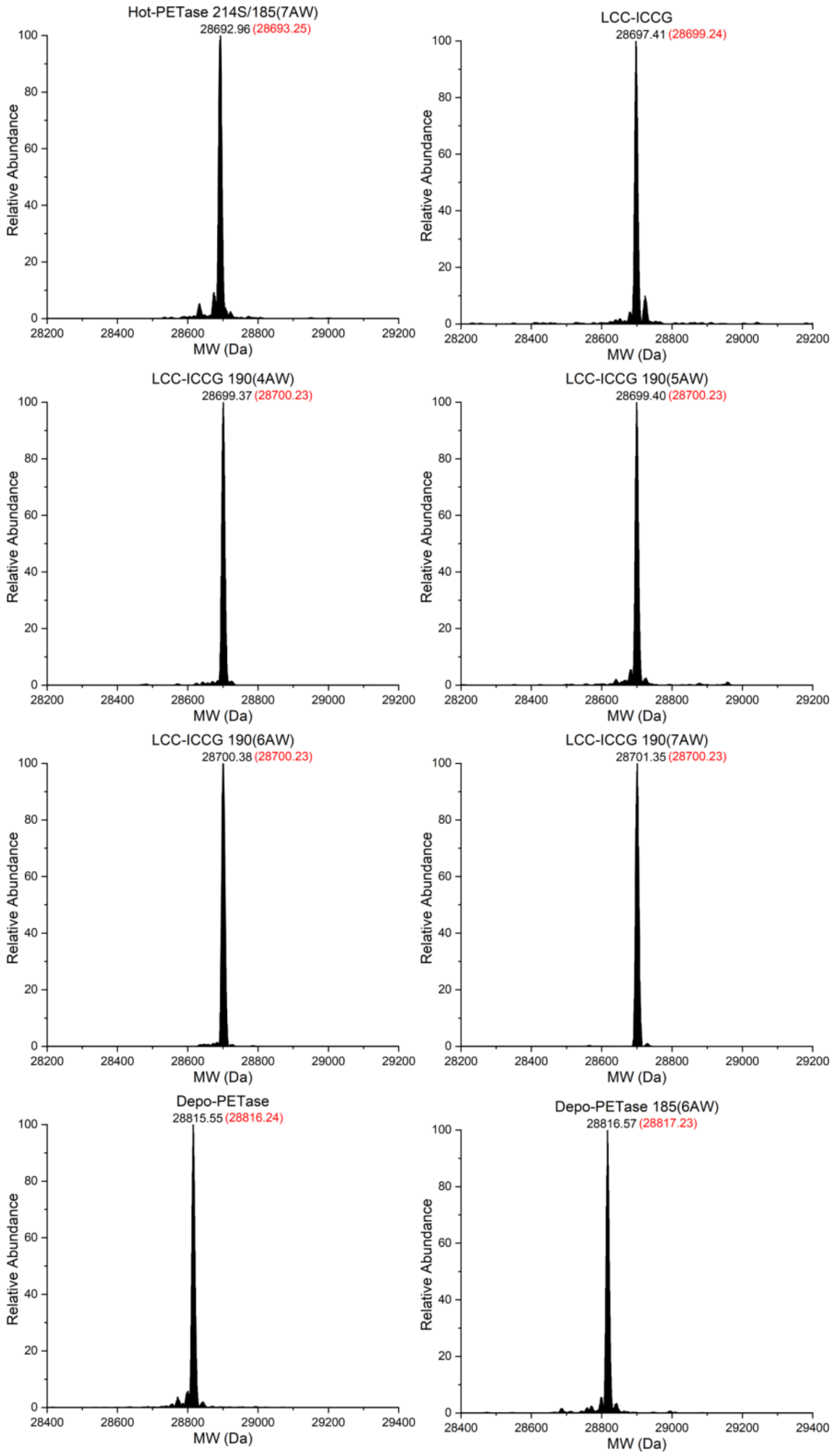
Hot-PETase 214S/185(7AW)
28692.96 (28693.25)
LCC-ICCG
28697.41 (28699.24)
LCC-ICCG 190(4AW)
28699.37 (28700.23)
LCC-ICCG 190(5AW)
28699.40 (28700.23)
LCC-ICCG 190(6AW)
28700.38 (28700.23)
LCC-ICCG 190(7AW)
28701.35 (28700.23)
Depo-PETase
28815.55 (28816.24)
Depo-PETase 185(6AW)
28816.57 (28817.23)
Relative Abundance
MW (Da)

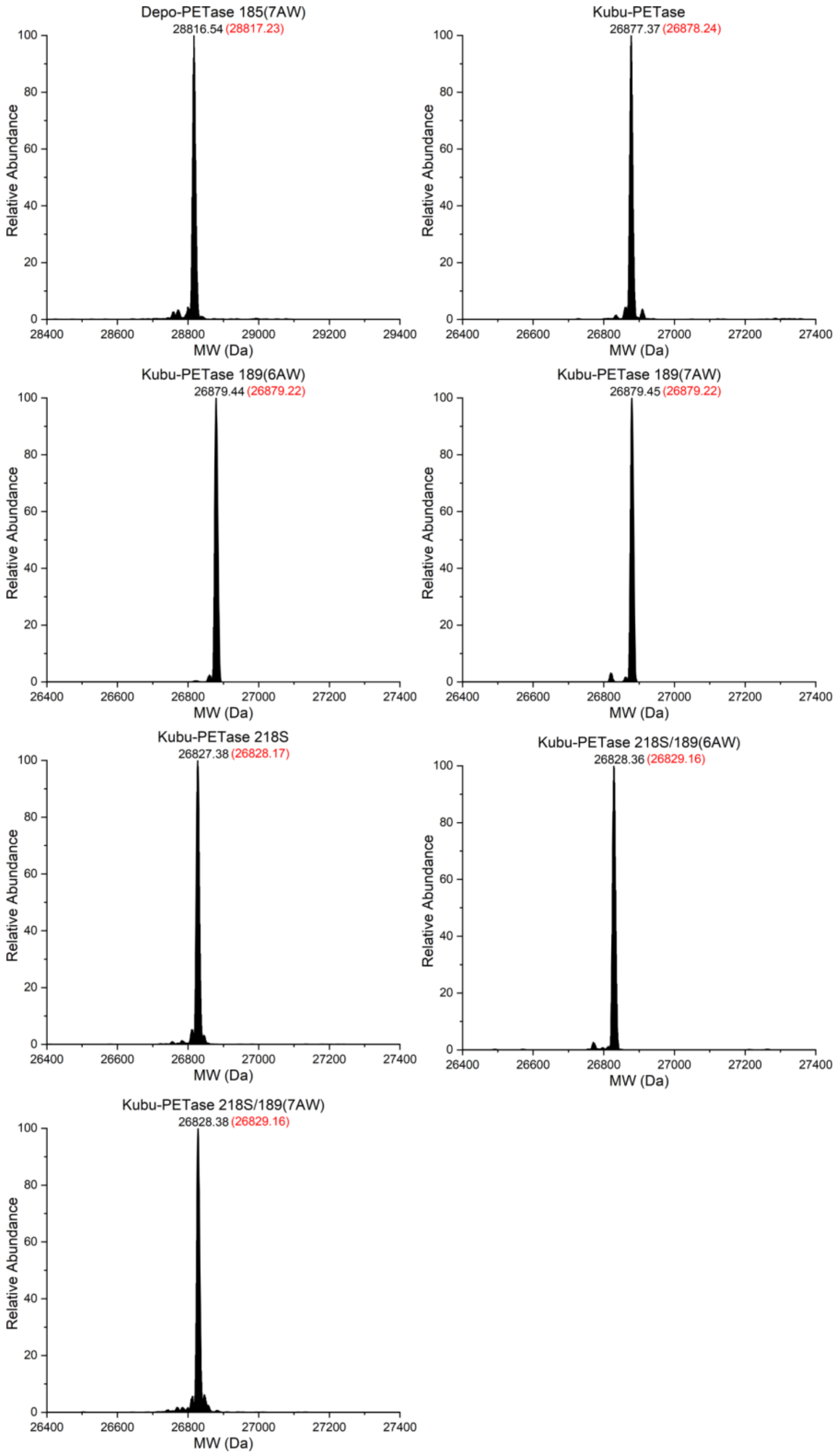

**Figure S18:** Intact protein mass spectrometry analysis of purified proteins. The observed and expected masses of the full-length proteins are shown in black and red, respectively. All LCC-ICCG and Kubu-PETase variants display loss of the N-terminal methionine. The expected masses were calculated assuming formation of two disulfide bonds in FAST-PETase, Depo-PETase, and LCC-ICCG and three disulfide bonds in Hot-PETase and Kubu-PETase variants. The peak corresponding to the near-cognate suppression of the amber stop codon by the glutaminyl-tRNA is indicated with an asterisk.

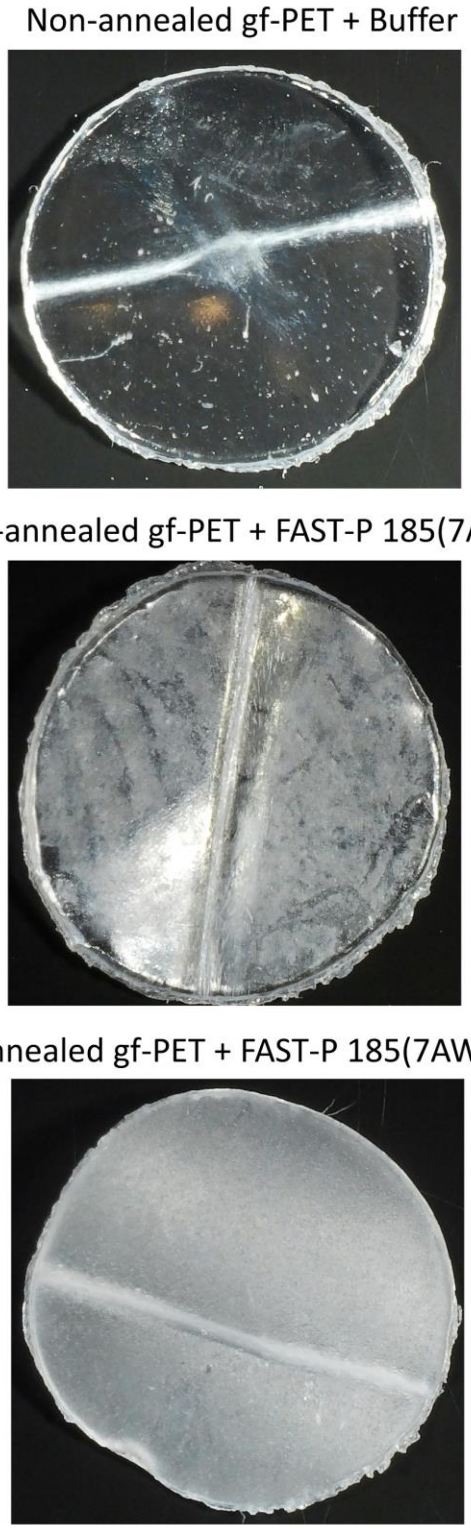

**Figure S19. Standardisation of gf-PET film by annealing for reproducible enzymatic degradation.** Commercial gf-PET films exhibit enthalpy relaxation due to polymer aging, which results in inconsistent degradation due to non-uniform crystallinity ($X_C$), leaving high $X_C$ regions undegraded (appearing clear). This inconsistency is eliminated by annealing the gf-PET discs to achieve a uniform $X_C$ of 8.2%,[60] which ensures homogeneous degradation. The assay shown was performed using 100 nM FAST-PETase 185(7AW) in 100 mM potassium phosphate buffer (pH 8.0) at 50 °C for 24 h.

### Supplementary Tables

**Table S1:** List of chemicals and their suppliers used in this study.

| Compound | Supplier | Catalogue number | Cost (USD/mol)[a] |
|---|---|---|---|
| 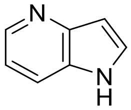 4-Azaindole | Ambeed (Illinois, USA) | A152186 | 330 |
| 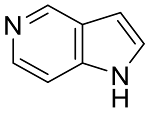 5-Azaindole | Ambeed (Illinois, USA) | A143044 | 220 |
| 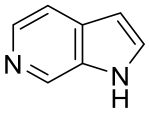 6-Azaindole | Ambeed (Illinois, USA) | A216265 | 320 |
| 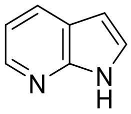 7-Azaindole | Ambeed (Illinois, USA) | A156097 | 30 |
| 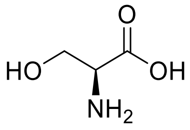 L-Serine | Merck KGaA (Darmstadt, Germany) | S4500 | |
| 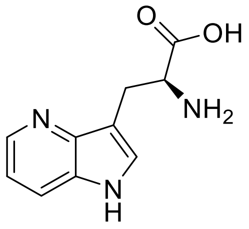 L-4Azatryptophan | Ambeed (Illinois, USA) | A264672 | 440,000 |
| 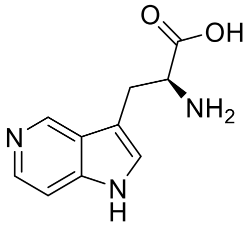 L-5Azatryptophan | Ambeed (Illinois, USA) | A332640 | 480,000 |
| 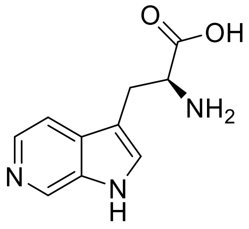 L-6Azatryptophan | Ambeed (Illinois, USA) | A861240 | 270,000 |

| | | | |
|---|---|---|---|
| 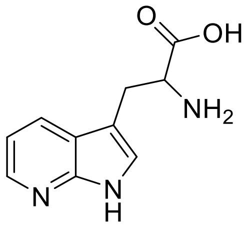 DL-7Azatryptophan | Ambeed (Illinois, USA) | A448683 | 70,000 |
| 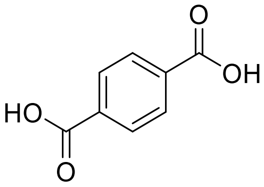 TPA | Ambeed (Illinois, USA) | A125601 | |
| 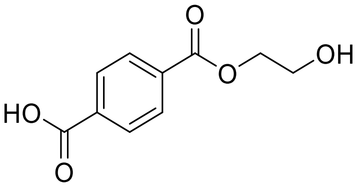 MHET | Ambeed (Illinois, USA) | A875019 | |
| 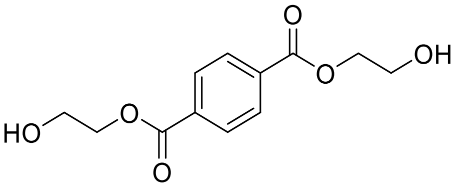 BHET | Merck KGaA (Darmstadt, Germany) | 465151 | |
| 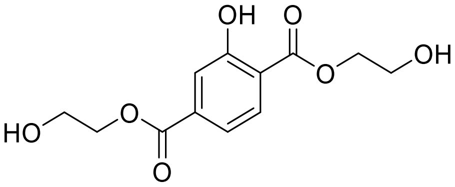 BHET-OH | Ambeed (Illinois, USA) | A2085224 | |
| Goodfellow PET film, thickness 0.25 mm, size 300 × 300 mm, condition transparent | Merck KGaA (Darmstadt, Germany) | GF25214475 | |

[a] Cost is calculated based of the largest available size (November 2025), normalised per mole.

**Table S2:** Photophysical properties of azatryptophan isomers measured in PBS buffer at 25 °C.

| Isomer | $\lambda_{ex}$max (nm) | $\lambda_{em}$max (nm) | Extinction coefficient[a] ($M^{-1}.cm^{-1}$) | Quantum yield (%)[b] | Brightness[c] ($M^{-1}$ $cm^{-1}$) |
|---|---|---|---|---|---|
| 4AW | 290 | 432 | 7364 ± 90 (at 290 nm)<br>6209 ± 76 (at 280 nm) | < 1 | - |
| 5AW | 280 | 427 | 2407 ± 11 (at 280 nm) | 2.7 ± 0.1 | 65 |
| 6AW | 325 | 402 | 2611 ± 47 (at 325 nm)<br>1116 ± 20 (at 280 nm) | 52.8 ± 1.2 | 1379 |
| 7AW | 290 | 404 | 7541 ± 56 (at 290 nm)<br>6453 ± 48 (at 280 nm) | 1.9 ± 0.1 | 143 |

[a] Molar extinction coefficients (ε) were measured at the wavelength of maximum absorption ($\lambda_{ex}$max) for each compound. In addition, molar extinction coefficients at 280 nm, $\varepsilon_{280}$, are reported, as $\varepsilon_{280}$ measurements are commonly used for determining protein concentrations using predictions based on amino acid composition.[61] As a control, the $\varepsilon_{280}$ value of L-tryptophan measured on the same instruments was 5516 ± 74 $M^{-1}.cm^{-1}$, in close agreement with the literature value of 5500 $M^{-1}.cm^{-1}$.[62]

[b] Quantum yields were determined at $\lambda_{ex}$max for the respective isomer, using the protocol of Würth et al. with either tryptophan (280–290 nm) or quinine sulfate (325 nm) as the reference standard.[63]

[c] Calculated as the product of molar extinction coefficient and quantum yield.

**Table S3:** Mutations in the G1PylRS variants selected for activity with 4AW, 5AW, and 6AW.

| RS Variants | Mutation Sites | | | | | | |
|---|---|---|---|---|---|---|---|
| *Mm*PylRS wild-type[a] | L305 | Y306 | N346 | V348 | Y384 | V401 | W417 |
| G1PylRS wild-type | L124 | Y125 | N165 | V167 | Y204 | A221 | W237 |
| 4AW08[b] | L | Y | A | F | W | H | T |
| 4AW32 | L | Y | A | F | W | T | C |
| 4AW34 | L | Y | A | F | W | T | C |
| 4AW57 | C | F | G | V | W | S | Y |
| 5AW02 | L | Y | D | F | W | S | S |
| 5AW21 | L | F | D | F | W | T | S |
| 5AW32[b] | L | Y | D | F | W | S | S |
| 6AW13[b] | T | L | D | A | W | S | W |
| 6AW14 | T | L | D | A | W | S | W |
| 6AW21 | E | M | S | A | Y | S | W |
| 6AW23 | T | L | D | A | W | S | W |
| 6AW28 | T | L | D | A | W | S | W |
| 6AW44 | T | L | D | A | W | S | W |

[a] Corresponding sites in the *Methanosarcina mazei* PylRS enzyme provided for comparison.

[b] The mutants 4AW08, 5AW32, and 6AW13 were used for all following applications and named 4AWRS, 5AWRS, and 6AWRS, respectively.

**Table S4: PETra-derived kinetic parameters for the PET hydrolases tested in this study.**

| Enzyme variant | $V_{max}$/mM·s$^{-1}$ | $K_m$/M | $k_{cat}$/s$^{-1}$ | Catalytic Efficiency/M$^{-1}$·s$^{-1}$ |
|---|---|---|---|---|
| FAST-PETase | 1.60 ± 0.07 | 2.10 ± 0.15 | 16.03 ± 0.67 | 7700 ± 400 |
| FAST-PETase 185(6AW) | 1.09 ± 0.02 | 1.93 ± 0.09 | 10.94 ± 0.23 | 5700 ± 100 |
| FAST-PETase 185(7AW) | 2.03 ± 0.07 | 2.23 ± 0.14 | 20.34 ± 0.65 | 9200 ± 300 |
| HOT-PETase | 0.39 ± 0.03 | 3.59 ± 0.25 | 3.92 ± 0.31 | 1100 ± 0 |
| HOT-PETase 214S | 0.90 ± 0.05 | 1.56 ± 0.12 | 9.01 ± 0.48 | 5800 ± 100 |
| HOT-PETase 185(6AW) | 0.42 ± 0.02 | 3.59 ± 0.26 | 4.23 ± 0.24 | 1200 ± 0 |
| HOT-PETase 185(7AW) | 0.45 ± 0.02 | 3.26 ± 0.17 | 4.46 ± 0.18 | 1400 ± 100 |
| HOT-PETase 214S/185(6AW) | 0.85 ± 0.07 | 1.62 ± 0.12 | 8.54 ± 0.69 | 5300 ± 300 |
| HOT-PETase 214S/185(7AW) | 0.76 ± 0.05 | 1.77 ± 0.11 | 7.59 ± 0.49 | 4300 ± 0 |
| Kubu-PETase | 0.15 ± 0.00 | 1.11 ± 0.08 | 1.48 ± 0.01 | 1300 ± 100 |
| Kubu-PETase 189(6AW) | 0.18 ± 0.01 | 1.37 ± 0.04 | 1.75 ± 0.05 | 1300 ± 0 |
| Kubu-PETase 189(7AW) | 0.21 ± 0.01 | 1.06 ± 0.10 | 2.10 ± 0.10 | 2000 ± 100 |
| Kubu-PETase 218S | 0.47 ± 0.03 | 0.75 ± 0.08 | 4.70 ± 0.34 | 6300 ± 200 |
| Kubu-PETase 218S/189(6AW) | 0.52 ± 0.02 | 0.86 ± 0.05 | 5.20 ± 0.18 | 6100 ± 400 |
| Kubu-PETase 218S/189(7AW) | 0.43 ± 0.01 | 0.63 ± 0.04 | 4.29 ± 0.08 | 6800 ± 300 |
| LCC-ICCG | 0.23 ± 0.01 | 4.43 ± 0.10 | 2.35 ± 0.06 | 500 ± 0 |
| LCC-ICCG 190(6AW) | 0.25 ± 0.02 | 3.80 ± 0.33 | 2.52 ± 0.22 | 700 ± 0 |
| LCC-ICCG 190(7AW) | 0.22 ± 0.02 | 2.88 ± 0.49 | 2.23 ± 0.22 | 800 ± 100 |
| Depo-PETase | 0.44 ± 0.04 | 3.50 ± 0.33 | 4.36 ± 0.37 | 1300 ± 100 |

**Table S5:** Melting temperatures (Tm) of the PET hydrolases used in this study and their azaPETases determined by differential scanning fluorimetry[a].

| Enzyme | Measured Tm/°C | Reported Tm/°C |
|---|---|---|
| LCC-ICCG | 96.3 ± 0.1 | 94.0[22] |
| LCC-ICCG 190(4AW) | 92.6 ± 0.2 | |
| LCC-ICCG 190(5AW) | 93.9 ± 0.1 | |
| LCC-ICCG 190(6AW) | 96.0 ± 0.1 | |
| LCC-ICCG 190(7AW) | 96.8 ± 0.1 | |
| FAST-PETase | 65.4 ± 0.5 | 67.1[19] |
| FAST-PETase 185(4AW) | 63.6 ± 0.1 | |
| FAST-PETase 185(5AW) | 62.8 ± 0.1 | |
| FAST-PETase 185(6AW) | 64.6 ± 0.2 | |
| FAST-PETase 185(7AW) | 65.4 ± 0.3 | |
| FAST-PETase 159(4AW) | 69.6 ± 0.1 | |
| FAST-PETase 159(5AW) | 61.8 ± 0.1 | |
| FAST-PETase 159(6AW) | 63.9 ± 0.1 | |
| FAST-PETase 159(7AW) | 60.8 ± 0.1 | |
| Hot-PETase | 83.0 ± 0.2 | 82.5[21] |
| Hot-PETase 185(4AW) | 83.5 ± 0.1 | |
| Hot-PETase 185(5AW) | 83.5 ± 0.1 | |
| Hot-PETase 185(6AW) | 83.9 ± 0.1 | |
| Hot-PETase 185(7AW) | 84.6 ± 0.1 | |
| Hot-PETase 214S | 78.8 ± 0.1 | |
| Hot-PETase 214S/185(6AW) | 78.7 ± 0.2 | |
| Hot-PETase 214S/185(7AW) | 79.0 ± 0.0 | |
| Depo-PETase | 68.9 ± 0.5 | 69.4[20] |
| Kubu-PETase | > 99.0[b] | > 99.0[23] |
| Kubu-PETase 189(6AW) | > 99.0[b] | |
| Kubu-PETase 189(7AW) | > 99.0[b] | |
| Kubu-PETase 218S | 92.0 ± 0.4[b] | |
| Kubu-PETase 218S/189(6AW) | 92.4 ± 0.2[b] | |
| Kubu-PETase 218S/189(7AW) | 92.4 ± 0.1[b] | |

[a] Tm values were measured using the Applied Biosystems Protein Thermal Shift kit on a QuantStudio™ 3 Real-Time PCR System and data were analysed by the Protein Thermal Shift™ Software v1.4 (Thermo Fisher Scientific Inc., USA). Data are shown as the mean of independent triplicates ± standard deviation.

[b] Tm values were determined by circular dichroism spectroscopy on a Chirascan Circular Dichroism Spectrometer (Applied Photophysics Ltd, UK).

**Table S6:** DNA and corresponding amino acid sequences of the proteins used in this study.

| Protein | DNA sequence | Amino acid sequence[a] |
|---|---|---|
| *Tm*9D8* TrpB-opt | ATGAAAGGCTACTTCGGTCCGTACGGTGGCCAGTACGTGCCAGAAAT<br>CCTGATGGGAGCTCTGGAAGAACTGGAAGCTGCGTACGAAGGCATCA<br>TGAAAGATGAGTCTTTCTGGAAAGAATTCAATGACCTGCTGCGCGAT<br>TATGCGGGTCGTCCGACTCCGCTGTACTTCGCACGTCGTCTGTCCGA<br>AAAATACGGTGCTCGCGTATATCTGAAACGTGAAGACCTGCTGCATA<br>CTGGTGCGCATAAAATCAATAACGCTATCGGCCAGGTTCTGCTGGCA<br>AAACTGATGGGCAAAACCCGTATCATTGCTGAAACGGGTGCTGGTCA<br>GCACGGCGTAGCAACTGCTACCGCAGCAGCGCTGTTCGGTATGGAAT<br>GTGTAATCTATATGGGCGAAGAAGACACGATCCGCCAGAAACTGAAC<br>GTTGAACGTATGAAACTGCTGGGTGCTAAAGTTGTACCGGTAAAATC<br>CGGTAGCCGTACCCTGAAAGACGCAATTGATGAAGCTCTGCGTGACT<br>GGATTACCAACCTGCAAACCACCTATTACGTGTTTGGCTCTGTGGTT<br>GGTCCGCATCCATATCCGATTATCGTACGTAACTTCCAAAAGGTTAT<br>CGGCGAAGAGACCAAAAAACAGATTCCGGAAAAAGAAGGCCGTCTGC<br>CGGACTACATCGTTGCATGCGTTAGCGGTGGTTCTAACGCTGCCGGT<br>ATCTTCTATCCGTTTATCGATTCTGGTGTGAAGCTGATCGGCGTAGA<br>AGCCGGTGGCGAAGGTCTGGAAACCGGTAAACATGCGGCTTCTCTGC<br>TGAAAGGTAAAATCGGCTACTTACACGGTTCTAAGACGTTCGTTCTG<br>CAGGATGACTGGGGCCAAGTTCAGGTGAGCCACTCCGTCTCCGCTGG<br>CCTGGACTACTCCGGTGTCGGTCCGGAACACGCCTATTGGCGTGAGA<br>CCGGTAAAGTGCTGTACGATGCTGTGACCGATGAAGAAGCTCTGGAC<br>GCATTCATCGAACTGTCTCGCCTGGAAGGCATCATCCCAGCCCTGGA<br>GTCTTCTCACGCACTGGCTTATCTGAAGAAGATCAACATCAAGGGTA<br>AAGTTGTGGTGGTTAATCTGTCTGGTCGTGGTGACAAGGATCTGGAA<br>TCTGTACTGAACCACCCGTATGTTCGCGAACGCATCCGCCTCGAGCA<br>TCACCACCATCATCACTAA | MKGYFGPYGGQYVPEILMGALEE<br>LEAAYEGIMKDESFWKEFNDLLR<br>DYAGRPTPLYFARRLSEKYGARV<br>YLKREDLLHTGAHKINNAIGQVL<br>LAKLMGKTRIIAETGAGQHGVAT<br>ATAAALFGMECVIYMGEEDTIRQ<br>KLNVERMKLLGAKVVPVKSGSRT<br>LKDAIDEALRDWITNLQTTYYVF<br>GSVVGPHPYPIIVRNFQKVIGEE<br>TKKQIPEKEGRLPDYIVACVSGG<br>SNAAGIFYPFIDSGVKLIGVEAG<br>GEGLETGKHAASLLKGKIGYLHG<br>SKTFVLQDDWGQVQVSHSVSAGL<br>DYSGVGPEHAYWRETGKVLYDAV<br>TDEEALDAFIELSRLEGIIPALE<br>SSHALAYLKKINIKGKVVVVNLS<br>GRGDKDLESVLNHPYVRERIRLE<br>HHHHHH |
| 4AWRS | ATGGTGGTGAAATTTACCGATAGCCAGATTCAGCATCTGATGGAATA<br>TGGTGATAATGATTGGAGCGAAGCCGAATTTGAAGATGCAGCAGCAC<br>GTGATAAAGAATTTAGCAGCCAGTTTAGCAAACTGAAAAGCGCCAAT<br>GATAAAGGCCTGAAAGATGTTATTGCAAATCCGCGTAATGATCTGAC<br>CGATCTGGAAAACAAAATTCGCGAAAAACTGGCAGCCCGTGGTTTTA<br>TTGAAGTTCATACCCCGATTTTTGTGAGCAAAAGCGCACTGGCAAAA<br>ATGACCATTACCGAAGATCATCCGCTGTTCAAACAGGTGTTTTGGAT<br>TGATGATAAACGTGCACTGCGTCCGATGCATGCAATGAATCTGTATA<br>AAGTTATGCGTGAACTGCGCGATCATACCAAAGGTCCGGTTAAAATC<br>TTTGAAATTGGTAGCTGCTTTCGCAAAGAAAGCAAAAGCAGTACCCA<br>TCTGGAAGAATTTACCATGCTGGCCCTGTTCGAAATGGGTCCTGATG<br>GTGATCCGATGGAACATCTGAAAATGTATATTGGCGATATCATGGAT<br>GCCGTTGGTGTTGAATATACCACCAGTCGTGAAGAATCAGATGTTTG<br>GGTTGAAACCCTGGACGTGGAAATTAATGGCACCGAAGTTGCAAGCG<br>GTCATGTTGGTCCGCATAAACTGGATCCGGCACATGATGTGCATGAA<br>CCGACGGCAGGTATTGGTTTTGGTCTGGAACGTCTGCTGATGCTGAA<br>AAATGGTAAAAGCAATGCACGCAAAACCGGCAAAAGTATTACCTATC<br>TGAATGGCTACAAACTGGATTAA | MVVKFTDSQIQHLMEYGDNDWSE<br>AEFEDAAARDKEFSSQFSKLKSA<br>NDKGLKDVIANPRNDLTDLENKI<br>REKLAARGFIEVHTPIFVSKSAL<br>AKMTITEDHPLFKQVFWIDDKRA<br>LRPMHAMNLYKVMRELRDHTKGP<br>VKIFEIGSCFRKESKSSTHLEEF<br>TMLALFEMGPDGDPMEHLKMYIG<br>DIMDAVGVEYTTSREESDVWVET<br>LDVEINGTEVASGHVGPHKLDPA<br>HDVHEPTAGIGFGLERLLMLKNG<br>KSNARKTGKSITYLNGYKLD |

| | | |
|---|---|---|
| 5AWRS | ATGGTGGTGAAATTTACCGATAGCCAGATTCAGCATCTGATGGAATA<br>TGGTGATAATGATTGGAGCGAAGCCGAATTTGAAGATGCAGCAGCAC<br>GTGATAAAGAATTTAGCAGCCAGTTTAGCAAACTGAAAAGCGCCAAT<br>GATAAAGGCCTGAAAGATGTTATTGCAAATCCGCGTAATGATCTGAC<br>CGATCTGGAAAACAAAATTCGCGAAAAACTGGCAGCCCGTGGTTTTA<br>TTGAAGTTCATACCCCGATTTTTGTGAGCAAAAGCGCACTGGCAAAA<br>ATGACCATTACCGAAGATCATCCGCTGTTCAAACAGGTGTTTTGGAT<br>TGATGATAAACGTGCACTGCGTCCGATGCATGCAATGAATCTGTATA<br>AAGTTATGCGTGAACTGCGCGATCATACCAAAGGTCCGGTTAAAATC<br>TTTGAAATTGGTAGCTGCTTTCGCAAAGAAAGCAAAAGCAGTACCCA<br>TCTGGAAGAATTTACCATGCTGGACCTGTTCGAAATGGGTCCTGATG<br>GTGATCCGATGGAACATCTGAAAATGTATATTGGCGATATCATGGAT<br>GCCGTTGGTGTTGAATATACCACCAGTCGTGAAGAATCAGATGTTTG<br>GGTTGAAACCCTGGACGTGGAAATTAATGGCACCGAAGTTGCAAGCG<br>GTTCTGTTGGTCCGCATAAACTGGATCCGGCACATGATGTGCATGAA<br>CCGAGTGCAGGTATTGGTTTTGGTCTGGAACGTCTGCTGATGCTGAA<br>AAATGGTAAAAGCAATGCACGCAAAACCGGCAAAAGTATTACCTATC<br>TGAATGGCTACAAACTGGATTAA | MVVKFTDSQIQHLMEYGDNDWSE<br>AEFEDAAARDKEFSSQFSKLKSA<br>NDKGLKDVIANPRNDLTDLENKI<br>REKLAARGFIEVHTPIFVSKSAL<br>AKMTITEDHPLFKQVFWIDDKRA<br>LRPMHAMNLYKVMRELRDHTKGP<br>VKIFEIGSCFRKESKSSTHLEEF<br>TMLDLFEMGPDGDPMEHLKMYIG<br>DIMDAVGVEYTTSREESDVWVET<br>LDVEINGTEVASGSVGPHKLDPA<br>HDVHEPSAGIGFGLERLLMLKNG<br>KSNARKTGKSITYLNGYKLD |
| 6AWRS | ATGGTGGTGAAATTTACCGATAGCCAGATTCAGCATCTGATGGAATA<br>TGGTGATAATGATTGGAGCGAAGCCGAATTTGAAGATGCAGCAGCAC<br>GTGATAAAGAATTTAGCAGCCAGTTTAGCAAACTGAAAAGCGCCAAT<br>GATAAAGGCCTGAAAGATGTTATTGCAAATCCGCGTAATGATCTGAC<br>CGATCTGGAAAACAAAATTCGCGAAAAACTGGCAGCCCGTGGTTTTA<br>TTGAAGTTCATACCCCGATTTTTGTGAGCAAAAGCGCACTGGCAAAA<br>ATGACCATTACCGAAGATCATCCGCTGTTCAAACAGGTGTTTTGGAT<br>TGATGATAAACGTGCACTGCGTCCGATGCATGCAATGAATACGCTTA<br>AAGTTATGCGTGAACTGCGCGATCATACCAAAGGTCCGGTTAAAATC<br>TTTGAAATTGGTAGCTGCTTTCGCAAAGAAAGCAAAAGCAGTACCCA<br>TCTGGAAGAATTTACCATGCTGGACCTGGCAGAAATGGGTCCTGATG<br>GTGATCCGATGGAACATCTGAAAATGTATATTGGCGATATCATGGAT<br>GCCGTTGGTGTTGAATATACCACCAGTCGTGAAGAATCAGATGTTTG<br>GGTTGAAACCCTGGACGTGGAAATTAATGGCACCGAAGTTGCAAGCG<br>GTAGTGTTGGTCCGCATAAACTGGATCCGGCACATGATGTGCATGAA<br>CCGTGGGCAGGTATTGGTTTTGGTCTGGAACGTCTGCTGATGCTGAA<br>AAATGGTAAAAGCAATGCACGCAAAACCGGCAAAAGTATTACCTATC<br>TGAATGGCTACAAACTGGATTAA | MVVKFTDSQIQHLMEYGDNDWSE<br>AEFEDAAARDKEFSSQFSKLKSA<br>NDKGLKDVIANPRNDLTDLENKI<br>REKLAARGFIEVHTPIFVSKSAL<br>AKMTITEDHPLFKQVFWIDDKRA<br>LRPMHAMNTLKVMRELRDHTKGP<br>VKIFEIGSCFRKESKSSTHLEEF<br>TMLDLAEMGPDGDPMEHLKMYIG<br>DIMDAVGVEYTTSREESDVWVET<br>LDVEINGTEVASGSVGPHKLDPA<br>HDVHEPWAGIGFGLERLLMLKNG<br>KSNARKTGKSITYLNGYKLD |
| 7AWRS[11] | ATGGTGGTGAAATTTACCGATAGCCAGATTCAGCATCTGATGGAATA<br>TGGTGATAATGATTGGAGCGAAGCCGAATTTGAAGATGCAGCAGCAC<br>GTGATAAAGAATTTAGCAGCCAGTTTAGCAAACTGAAAAGCGCCAAT<br>GATAAAGGCCTGAAAGATGTTATTGCAAATCCGCGTAATGATCTGAC<br>CGATCTGGAAAACAAAATTCGCGAAAAACTGGCAGCCCGTGGTTTTA<br>TTGAAGTTCATACCCCGATTTTTGTGAGCAAAAGCGCACTGGCAAAA<br>ATGACCATTACCGAAGATCATCCGCTGTTCAAACAGGTGTTTTGGAT<br>TGATGATAAACGTGCACTGCGTCCGATGCATGCAATGAATATGATGA<br>AAGTTATGCGTGAACTGCGCGATCATACCAAAGGTCCGGTTAAAATC<br>TTTGAAATTGGTAGCTGCTTTCGCAAAGAAAGCAAAAGCAGTACCCA<br>TCTGGAAGAATTTACCATGCTGGCCCTGGGAGAAATGGGTCCTGATG<br>GTGATCCGATGGAACATCTGAAAATGTATATTGGCGATATCATGGAT<br>GCCGTTGGTGTTGAATATACCACCAGTCGTGAAGAATCAGATGTTTA<br>TGTTGAAACCCTGGACGTGGAAATTAATGGCACCGAAGTTGCAAGCG<br>GTGGGGTTGGTCCGCATAAACTGGATCCGGCACATGATGTGCATGAA<br>CCGCATGCAGGTATTGGTTTTGGTCTGGAACGTCTGCTGATGCTGAA<br>AAATGGTAAAAGCAATGCACGCAAAACCGGCAAAAGTATTACCTATC<br>TGAATGGCTACAAACTGGATTAA | MVVKFTDSQIQHLMEYGDNDWSE<br>AEFEDAAARDKEFSSQFSKLKSA<br>NDKGLKDVIANPRNDLTDLENKI<br>REKLAARGFIEVHTPIFVSKSAL<br>AKMTITEDHPLFKQVFWIDDKRA<br>LRPMHAMNMMKVMRELRDHTKGP<br>VKIFEIGSCFRKESKSSTHLEEF<br>TMLALGEMGPDGDPMEHLKMYIG<br>DIMDAVGVEYTTSREESDVYVET<br>LDVEINGTEVASGGVGPHKLDPA<br>HDVHEPHAGIGFGLERLLMLKNG<br>KSNARKTGKSITYLNGYKLD |

| | | |
|---|---|---|
| His$_6$-TAG-RFP | ATGCACCACCATCACCATCACTAGGCCAGTAGTGAAGACGTTATCAA<br>GGAGTTTATGCGTTTCAAAGTACGTATGGAGGGTAGTGTTAACGGAC<br>ACGAATTTGAGATCGAGGGAGAGGGGGAAGGTCGTCCTTACGAGGGA<br>ACTCAAACGGCCAAATTAAAGGTGACCAAAGGTGGGCCCTTGCCATT<br>CGCGTGGGACATCTTGTCACCCCAGTTCCAGTACGGGTCGAAGGCAT<br>ACGTAAAACACCCAGCGGACATTCCTGACTATCTTAAGTTATCTTTC<br>CCGGAAGGTTTTAAATGGGAACGCGTGATGAACTTTGAGGATGGGGG<br>GGTTGTTACGGTGACACAAGACTCCTCATTGCAAGATGGAGAGTTTA<br>TCTATAAAGTCAAACTTCGCGGCACCAATTTTCCATCTGACGGTCCT<br>GTAATGCAGAAAAAAACAATGGGCTGGGAAGCCTCCACAGAACGTAT<br>GTACCCCGAAGATGGAGCTTTAAAGGGCGAAATTAAAATGCGCTTAA<br>AACTTAAAGACGGCGGCCATTACGACGCCGAAGTGAAAACGACGTAT<br>ATGGCTAAGAAACCCGTCCAGCTTCCGGGAGCCTATAAAACTGACAT<br>CAAACTGGATATTACATCACACAACGAAGATTATACTATTGTCGAAC<br>AGTACGAACGCGCCGAAGGCCGCCATTCAACGGGAGCATAA | MHHHHHHXASSEDVIKEFMRFKV<br>RMEGSVNGHEFEIEGEGEGRPYE<br>GTQTAKLKVTKGGPLPFAWDILS<br>PQFQYGSKAYVKHPADIPDYLKL<br>SFPEGFKWERVMNFEDGGVVTVT<br>QDSSLQDGEFIYKVKLRGTNFPS<br>DGPVMQKKTMGWEASTERMYPED<br>GALKGEIKMRLKLKDGGHYDAEV<br>KTTYMAKKPVQLPGAYKTDIKLD<br>ITSHNEDYTIVEQYERAEGRHST<br>GA |
| NT*-<br>ENLYFQGDX | ATGCATCATCATCATCACCACAGCCATACCACACCGTGGACCAATCC<br>TGGTCTGGCAGAAAACTTTATGAATAGCTTTATGCAGGGTCTGAGCA<br>GCATGCCTGGTTTTACCGCAAGCCAGCTGGACAAAATGAGCACCATT<br>GCACAGAGCATGGTTCAGAGCATTCAGAGCCTGGCAGCACAGGGTCG<br>TACCAGTCCGAATGATCTGCAGGCACTGAATATGGCATTTGCAAGCA<br>GCATGGCAGAAATTGCAGCAAGCGAAGAAGGTGGCGGTAGCCTGAGC<br>ACCAAAACCAGCAGCATTGCAAGCGCAATGAGCAATGCATTTCTGCA<br>GACAACCGGTGTTGTTAATCAGCCGTTTATTAACGAAATTACCCAGC<br>TGGTTAGCATGTTTGCACAGGCAGGTATGAATGATGTTAGCGCAGAA<br>AACCTGTACTTTCAAGGCGATTAGTAA | MHHHHHHSHTTPWTNPGLAENFM<br>NSFMQGLSSMPGFTASQLDKMST<br>IAQSMVQSIQSLAAQGRTSPNDL<br>QALNMAFASSMAEIAASEEGGGS<br>LSTKTSSIASAMSNAFLQTTGVV<br>NQPFINEITQLVSMFAQAGMNDV<br>SAENLYFQGDX |
| FAST-PETase | ATGCAGACCAATCCGTATGCACGTGGTCCGAATCCGACCGCAGCAAG<br>CCTGGAAGCAAGCGCAGGTCCGTTTACCGTTCGTAGCTTTACCGTTA<br>GCCGTCCGAGCGGTTATGGTGCAGGCACCGTTTATTATCCGACCAAT<br>GCCGGTGGCACCGTTGGTGCAATTGCCATTGTTCCGGGTTATACCGC<br>ACGTCAGAGCAGCATTAAATGGTGGGGTCCGCGTCTGGCAAGCCATG<br>GTTTTGTTGTTATTACCATTGATACCAACAGCACCCTGGATCAGCCG<br>GAAAGCCGTAGCAGCCAGCAGATGGCAGCACTGCGTCAGGTTGCCAG<br>CCTGAATGGCACCAGCAGCAGCCCGATTTATGGTAAAGTTGATACAG<br>CACGTATGGGTGTTATGGGTTGGAGCATGGGTGGTGGTGGTAGCCTG<br>ATTAGCGCAGCAAATAATCCGAGCCTGAAAGCAGCAGCACCGCAGGC<br>TCCGTGGCATAGCAGCACCAATTTTAGCAGCGTTACCGTTCCGACAC<br>TGATTTTTGCATGTGAAAATGATAGCATTGCACCGGTTAATAGCAGC<br>GCACTGCCGATCTATGATAGTATGAGCCAGAATGCAAAACAGTTTCT<br>GGAAATTAAAGGTGGCAGCCATAGCTGTGCAAATAGCGGTAATAGCA<br>ATCAGGCACTGATCGGTAAAAAGGGTGTTGCATGGATGAAACGCTTT<br>ATGGATAATGATACCCGCTATAGCACCTTTGCCTGCGAAAATCCGAA<br>TAGTACCGCAGTTAGCGATTTTCGTACCGCAAATTGTAGCCTGGAAC<br>ATCATCACCATCATCATTAA | MQTNPYARGPNPTAASLEASAGP<br>FTVRSFTVSRPSGYGAGTVYYPT<br>NAGGTVGAIAIVPGYTARQSSIK<br>WWGPRLASHGFVVITIDTNSTLD<br>QPESRSSQQMAALRQVASLNGTS<br>SSPIYGKVDTARMGVMGWSMGGG<br>GSLISAANNPSLKAAAPQAPWHS<br>STNFSSVTVPTLIFACENDSIAP<br>VNSSALPIYDSMSQNAKQFLEIK<br>GGSHSCANSGNSNQALIGKKGVA<br>WMKRFMDNDTRYSTFACENPNST<br>AVSDFRTANCSLEHHHHHH |
| FAST-PETase<br>159TAG | ATGCAGACCAATCCGTATGCACGTGGTCCGAATCCGACCGCAGCAAG<br>CCTGGAAGCAAGCGCAGGTCCGTTTACCGTTCGTAGCTTTACCGTTA<br>GCCGTCCGAGCGGTTATGGTGCAGGCACCGTTTATTATCCGACCAAT<br>GCCGGTGGCACCGTTGGTGCAATTGCCATTGTTCCGGGTTATACCGC<br>ACGTCAGAGCAGCATTAAATGGTGGGGTCCGCGTCTGGCAAGCCATG<br>GTTTTGTTGTTATTACCATTGATACCAACAGCACCCTGGATCAGCCG<br>GAAAGCCGTAGCAGCCAGCAGATGGCAGCACTGCGTCAGGTTGCCAG<br>CCTGAATGGCACCAGCAGCAGCCCGATTTATGGTAAAGTTGATACAG<br>CACGTATGGGTGTTATGGGTTAGAGCATGGGTGGTGGTGGTAGCCTG<br>ATTAGCGCAGCAAATAATCCGAGCCTGAAAGCAGCAGCACCGCAGGC<br>TCCGTGGCATAGCAGCACCAATTTTAGCAGCGTTACCGTTCCGACAC<br>TGATTTTTGCATGTGAAAATGATAGCATTGCACCGGTTAATAGCAGC<br>GCACTGCCGATCTATGATAGTATGAGCCAGAATGCAAAACAGTTTCT<br>GGAAATTAAAGGTGGCAGCCATAGCTGTGCAAATAGCGGTAATAGCA<br>ATCAGGCACTGATCGGTAAAAAGGGTGTTGCATGGATGAAACGCTTT<br>ATGGATAATGATACCCGCTATAGCACCTTTGCCTGCGAAAATCCGAA<br>TAGTACCGCAGTTAGCGATTTTCGTACCGCAAATTGTAGCCTGGAAC<br>ATCATCACCATCATCATTAA | MQTNPYARGPNPTAASLEASAGP<br>FTVRSFTVSRPSGYGAGTVYYPT<br>NAGGTVGAIAIVPGYTARQSSIK<br>WWGPRLASHGFVVITIDTNSTLD<br>QPESRSSQQMAALRQVASLNGTS<br>SSPIYGKVDTARMGVMGXSMGGG<br>GSLISAANNPSLKAAAPQAPWHS<br>STNFSSVTVPTLIFACENDSIAP<br>VNSSALPIYDSMSQNAKQFLEIK<br>GGSHSCANSGNSNQALIGKKGVA<br>WMKRFMDNDTRYSTFACENPNST<br>AVSDFRTANCSLEHHHHHH |

| | | |
|---|---|---|
| FAST-PETase<br>185TAG | ATGCAGACCAATCCGTATGCACGTGGTCCGAATCCGACCGCAGCAAG<br>CCTGGAAGCAAGCGCAGGTCCGTTTACCGTTCGTAGCTTTACCGTTA<br>GCCGTCCGAGCGGTTATGGTGCAGGCACCGTTTATTATCCGACCAAT<br>GCCGGTGGCACCGTTGGTGCAATTGCCATTGTTCCGGGTTATACCGC<br>ACGTCAGAGCAGCATTAAATGGTGGGGTCCGCGTCTGGCAAGCCATG<br>GTTTTGTTGTTATTACCATTGATACCAACAGCACCCTGGATCAGCCG<br>GAAAGCCGTAGCAGCCAGCAGATGGCAGCACTGCGTCAGGTTGCCAG<br>CCTGAATGGCACCAGCAGCAGCCCGATTTATGGTAAAGTTGATACAG<br>CACGTATGGGTGTTATGGGTTGGAGCATGGGTGGTGGTAGCCTG<br>ATTAGCGCAGCAAATAATCCGAGCCTGAAAGCAGCAGCACCGCAGGC<br>TCCGTAGCATAGCAGCACCAATTTTAGCAGCGTTACCGTTCCGACAC<br>TGATTTTTGCATGTGAAAATGATAGCATTGCACCGGTTAATAGCAGC<br>GCACTGCCGATCTATGATAGTATGAGCCAGAATGCAAAACAGTTTCT<br>GGAAATTAAAGGTGGCAGCCATAGCTGTGCAAATAGCGGTAATAGCA<br>ATCAGGCACTGATCGGTAAAAAGGGTGTTGCATGGATGAAACGCTTT<br>ATGGATAATGATACCCGCTATAGCACCTTTGCCTGCGAAAATCCGAA<br>TAGTACCGCAGTTAGCGATTTTCGTACCGCAAATTGTAGCCTGGAAC<br>ATCATCACCATCATCATTAA | MQTNPYARGPNPTAASLEASAGP<br>FTVRSFTVSRPSGYGAGTVYYPT<br>NAGGTVGAIAIVPGYTARQSSIK<br>WWGPRLASHGFVVITIDTNSTLD<br>QPESRSSQQMAALRQVASLNGTS<br>SSPIYGKVDTARMGVMGWSMGGG<br>GSLISAANNPSLKAAAPQAPXHS<br>STNFSSVTVPTLIFACENDSIAP<br>VNSSALPIYDSMSQNAKQFLEIK<br>GGSHSCANSGNSNQALIGKKGVA<br>WMKRFMDNDTRYSTFACENPNST<br>AVSDFRTANCSLEHHHHHH |
| Depo-PETase | ATGCAGACCAATCCGTATGCACGTGGTCCGAATCCGACCGCAGCAAG<br>CCTGGAAGCAAGCGCAGGTCCGTTTACCGTTCGTAGCTTTACCGTTA<br>GCCGTCCGAGCGGTTATGGTGCAGGCACCGTTTATTATCCGACCAAT<br>GCCGGTGGCACCGTTGGTGCAATTGCCATTGTTCCGGGTTATATTGC<br>ACGTCAGAGCAGCATTAAATGGTGGGGTCCGCGTCTGGCAAGCCATG<br>GTTTTGTTGTTATTACCATTGATACCAACAGCACCCTGGATCAGCCG<br>AGTAGCCGTAGCAGCCAGCAGATGGCAGCACTGCGTCAGGTTGCCAG<br>CCTGAATGGCACCAGCAGCAGCCCGATTTATGGTAAAGTTGATACAG<br>CACGTATGGGTGTTATGGGTTGGAGCATGGGTGGTGGTAGCCTG<br>ATTAGCGCAGCAAATAATCCGAGCCTGAAAGCAGCAGCACCGCAGGC<br>TCCGTGGCATAGCAGCACCAATTTTAGCAGCGTTACCGTTCCGACAC<br>TGATTTTTGCATGTGAAAATGATAGCATTGCACCGGTTAATAGCAGC<br>GCACTGCCGATCTATAACAGTATGAGCCGCAATGCAAAACAGTTTCT<br>GGAAATTAAAGGTGGCAGCCATAGCTGTGCAAATAGCGGTAATAGCG<br>ATCAGGCACTGATCGGTAAAAAGGGTGTTGCATGGATGAAATATTTT<br>ATGGATAATGATACCCGCTATAGCACCTTTGCCTGCGAAAATCCGAA<br>TAGTACCCGCGTTAGCGATTTTCGTACCGCAAATTGTCCGCTGGAAC<br>ATCATCACCATCATCATTAA | MQTNPYARGPNPTAASLEASAGP<br>FTVRSFTVSRPSGYGAGTVYYPT<br>NAGGTVGAIAIVPGYIARQSSIK<br>WWGPRLASHGFVVITIDTNSTLD<br>QPSSRSSQQMAALRQVASLNGTS<br>SSPIYGKVDTARMGVMGWSMGGG<br>GSLISAANNPSLKAAAPQAPWHS<br>STNFSSVTVPTLIFACENDSIAP<br>VNSSALPIYNSMSRNAKQFLEIK<br>GGSHSCANSGNSDQALIGKKGVA<br>WMKYFMDNDTRYSTFACENPNST<br>RVSDFRTANCPLEHHHHHH |
| Depo-PETase<br>185TAG | ATGCAGACCAATCCGTATGCACGTGGTCCGAATCCGACCGCAGCAAG<br>CCTGGAAGCAAGCGCAGGTCCGTTTACCGTTCGTAGCTTTACCGTTA<br>GCCGTCCGAGCGGTTATGGTGCAGGCACCGTTTATTATCCGACCAAT<br>GCCGGTGGCACCGTTGGTGCAATTGCCATTGTTCCGGGTTATATTGC<br>ACGTCAGAGCAGCATTAAATGGTGGGGTCCGCGTCTGGCAAGCCATG<br>GTTTTGTTGTTATTACCATTGATACCAACAGCACCCTGGATCAGCCG<br>AGCAGCCGTAGCAGTCAGCAGATGGCAGCACTGCGTCAGGTTGCCAG<br>CCTGAATGGCACCAGCAGCAGCCCGATTTATGGTAAAGTTGATACCG<br>CACGTATGGGTGTTATGGGTTGGAGCATGGGTGGTGGTAGCCTG<br>ATTAGCGCAGCAAATAATCCGAGCCTGAAAGCAGCAGCACCGCAGGC<br>TCCGTAGCATAGCAGCACCAATTTTAGCAGCGTTACCGTTCCGACAC<br>TGATTTTTGCATGTGAAAATGATAGCATTGCACCGGTTAATAGCAGC<br>GCACTGCCGATTTATAACAGCATGAGCCGTAATGCAAAACAGTTTCT<br>GGAAATTAAAGGTGGCAGCCATAGCTGTGCAAATAGCGGTAATAGCG<br>ATCAGGCACTGATTGGTAAAAAGGGTGTTGCATGGATGAAATACTTC<br>ATGGATAATGATACCCGCTATAGCACCTTTGCCTGCGAAAATCCGAA<br>TAGCACCCGTGTTAGCGATTTTCGTACCGCAAATTGTCCGCTGGAAC<br>ATCATCACCATCATCATTAA | MQTNPYARGPNPTAASLEASAGP<br>FTVRSFTVSRPSGYGAGTVYYPT<br>NAGGTVGAIAIVPGYIARQSSIK<br>WWGPRLASHGFVVITIDTNSTLD<br>QPSSRSSQQMAALRQVASLNGTS<br>SSPIYGKVDTARMGVMGWSMGGG<br>GSLISAANNPSLKAAAPQAPXHS<br>STNFSSVTVPTLIFACENDSIAP<br>VNSSALPIYNSMSRNAKQFLEIK<br>GGSHSCANSGNSDQALIGKKGVA<br>WMKYFMDNDTRYSTFACENPNST<br>RVSDFRTANCPLEHHHHHH |
| Hot-PETase | ATGCAGACCAATCCGTATGCACGTGGTCCGAATCCGACCGCAGCAAG<br>CCTGGAAGCAAGCGCAGGTCCGTTTACCGTTCGTAGCTTTACCGTTG<br>CACGTCCGGTTGGTTATGGTGCAGGCACCGTTTATTATCCGACCAAT<br>GCCGGTGGCACCGTTGGTGCAATTGCCATTGTTCCGGGTTATACCGC<br>AACACAGAGCAGCATTAATTGGTGGGGTCCGCGTCTGGCAAGCCATG<br>GTTTTGTTGTTATTACCATTGATACCAACAGCACCCTGGATAAACCG<br>GAAAGCCGTAGCAGCCAGCAGATGGCAGCACTGCGTCAGGTTGCCAG<br>CCTGAATGGCACCAGCAGCAGCCCGATTTATGGTAAAGTTGATACCG<br>CACGTGGTGGTGTTATGGGTTGGAGCATGGGTGGTGGTAGCCTG | MQTNPYARGPNPTAASLEASAGP<br>FTVRSFTVARPVGYGAGTVYYPT<br>NAGGTVGAIAIVPGYTATQSSIN<br>WWGPRLASHGFVVITIDTNSTLD<br>KPESRSSQQMAALRQVASLNGTS<br>SSPIYGKVDTARGGVMGWSMGGG<br>GSLISAANNPSLKAAVMAPWHS<br>STNFSSVTVPTLIFACENDRIAP<br>VKEYALPIYDSMSLNAKQFLEIC |

| | | |
|---|---|---|
| | ATTAGCGCAGCAAATAATCCGAGCCTGAAAGCAGCAGCAGTTATGGC<br>ACCGTGGCATAGCAGCACCAATTTTAGCAGCGTTACCGTTCCGACAC<br>TGATTTTTGCATGTGAAAATGATCGTATCGCACCGGTTAAAGAATAT<br>GCACTGCCGATCTATGATAGCATGAGTCTGAATGCAAAACAGTTCCT<br>GGAAATTTGTGGTGGTTCACATAGCTGTGCATGTAGCGGTAATAGCA<br>ATCAGGCACTGATCGGTATGAAAGGTGTTGCATGGATGAAACGCTTT<br>ATGGATAATGATACCCGCTATAGCCAGTTTGCCTGCGAAAATCCGAA<br>TAGCACCGCAGTTTGTGATTTTCGTACCGCAAATTGTAGCCTGGAAC<br>ATCATCACCATCATCATTAA | GGSHSCACSGNSNQALIGMKGVA<br>WMKRFMDNDTRYSQFACENPNST<br>AVCDFRTANCSLEHHHHHH |
| Hot-PETase 185TAG | ATGCAGACCAATCCGTATGCACGTGGTCCGAATCCGACCGCAGCAAG<br>CCTGGAAGCAAGCGCAGGTCCGTTTACCGTTCGTAGCTTTACCGTTG<br>CACGTCCGGTTGGTTATGGTGCAGGCACCGTTTATTATCCGACCAAT<br>GCCGGTGGCACCGTTGGTGCAATTGCCATTGTTCCGGGTTATACCGC<br>AACACAGAGCAGCATTAATTGGTGGGGTCCGCGTCTGGCAAGCCATG<br>GTTTTGTTGTTATTACCATTGATACCAACAGCACCCTGGATAAACCG<br>GAAAGCCGTAGCAGCCAGCAGATGGCAGCACTGCGTCAGGTTGCCAG<br>CCTGAATGGCACCAGCAGCAGCCCGATTTATGGTAAAGTTGATACCG<br>CACGTGGTGGTGTTATGGGTTGGAGCATGGGTGGTGGTGGTAGCCTG<br>ATTAGCGCAGCAAATAATCCGAGCCTGAAAGCAGCAGCAGTTATGGC<br>ACCGTAGCATAGCAGCACCAATTTTAGCAGCGTTACCGTTCCGACAC<br>TGATTTTTGCATGTGAAAATGATCGTATCGCACCGGTTAAAGAATAT<br>GCACTGCCGATCTATGATAGCATGAGTCTGAATGCAAAACAGTTCCT<br>GGAAATTTGTGGTGGTTCACATAGCTGTGCATGTAGCGGTAATAGCA<br>ATCAGGCACTGATCGGTATGAAAGGTGTTGCATGGATGAAACGCTTT<br>ATGGATAATGATACCCGCTATAGCCAGTTTGCCTGCGAAAATCCGAA<br>TAGCACCGCAGTTTGTGATTTTCGTACCGCAAATTGTAGCCTGGAAC<br>ATCATCACCATCATCATTAA | MQTNPYARGPNPTAASLEASAGP<br>FTVRSFTVARPVGYGAGTVYYPT<br>NAGGTVGAIAIVPGYTATQSSIN<br>WWGPRLASHGFVVITIDTNSTLD<br>KPESRSSQQMAALRQVASLNGTS<br>SSPIYGKVDTARGGVMGWSMGGG<br>GSLISAANNPSLKAAAVMAPXHS<br>STNFSSVTVPTLIFACENDRIAP<br>VKEYALPIYDSMSLNAKQFLEIC<br>GGSHSCACSGNSNQALIGMKGVA<br>WMKRFMDNDTRYSQFACENPNST<br>AVCDFRTANCSLEHHHHHH |
| Hot-PETase 214S | ATGCAGACCAATCCGTATGCACGTGGTCCGAATCCGACCGCAGCAAG<br>CCTGGAAGCAAGCGCAGGTCCGTTTACCGTTCGTAGCTTTACCGTTG<br>CACGTCCGGTTGGTTATGGTGCAGGCACCGTTTATTATCCGACCAAT<br>GCCGGTGGCACCGTTGGTGCAATTGCCATTGTTCCGGGTTATACCGC<br>AACACAGAGCAGCATTAATTGGTGGGGTCCGCGTCTGGCAAGCCATG<br>GTTTTGTTGTTATTACCATTGATACCAACAGCACCCTGGATAAACCG<br>GAAAGCCGTAGCAGCCAGCAGATGGCAGCACTGCGTCAGGTTGCCAG<br>CCTGAATGGCACCAGCAGCAGCCCGATTTATGGTAAAGTTGATACCG<br>CACGTGGTGGTGTTATGGGTTGGAGCATGGGTGGTGGTGGTAGCCTG<br>ATTAGCGCAGCAAATAATCCGAGCCTGAAAGCAGCAGCAGTTATGGC<br>ACCGTGGCATAGCAGCACCAATTTTAGCAGCGTTACCGTTCCGACAC<br>TGATTTTTGCATGTGAAAATGATCGTATCGCACCGGTTAAAGAAAGC<br>GCACTGCCGATCTATGATAGCATGAGTCTGAATGCAAAACAGTTCCT<br>GGAAATTTGTGGTGGTTCACATAGCTGTGCATGTAGCGGTAATAGCA<br>ATCAGGCACTGATCGGTATGAAAGGTGTTGCATGGATGAAACGCTTT<br>ATGGATAATGATACCCGCTATAGCCAGTTTGCCTGCGAAAATCCGAA<br>TAGCACCGCAGTTTGTGATTTTCGTACCGCAAATTGTAGCCTGGAAC<br>ATCATCACCATCATCATTAA | MQTNPYARGPNPTAASLEASAGP<br>FTVRSFTVARPVGYGAGTVYYPT<br>NAGGTVGAIAIVPGYTATQSSIN<br>WWGPRLASHGFVVITIDTNSTLD<br>KPESRSSQQMAALRQVASLNGTS<br>SSPIYGKVDTARGGVMGWSMGGG<br>GSLISAANNPSLKAAAVMAPWHS<br>STNFSSVTVPTLIFACENDRIAP<br>VKESALPIYDSMSLNAKQFLEIC<br>GGSHSCACSGNSNQALIGMKGVA<br>WMKRFMDNDTRYSQFACENPNST<br>AVCDFRTANCSLEHHHHHH |
| Hot-PETase 214S/185TAG | ATGCAGACCAATCCGTATGCACGTGGTCCGAATCCGACCGCAGCAAG<br>CCTGGAAGCAAGCGCAGGTCCGTTTACCGTTCGTAGCTTTACCGTTG<br>CACGTCCGGTTGGTTATGGTGCAGGCACCGTTTATTATCCGACCAAT<br>GCCGGTGGCACCGTTGGTGCAATTGCCATTGTTCCGGGTTATACCGC<br>AACACAGAGCAGCATTAATTGGTGGGGTCCGCGTCTGGCAAGCCATG<br>GTTTTGTTGTTATTACCATTGATACCAACAGCACCCTGGATAAACCG<br>GAAAGCCGTAGCAGCCAGCAGATGGCAGCACTGCGTCAGGTTGCCAG<br>CCTGAATGGCACCAGCAGCAGCCCGATTTATGGTAAAGTTGATACCG<br>CACGTGGTGGTGTTATGGGTTGGAGCATGGGTGGTGGTGGTAGCCTG<br>ATTAGCGCAGCAAATAATCCGAGCCTGAAAGCAGCAGCAGTTATGGC<br>ACCGTAGCATAGCAGCACCAATTTTAGCAGCGTTACCGTTCCGACAC<br>TGATTTTTGCATGTGAAAATGATCGTATCGCACCGGTTAAAGAAAGC<br>GCACTGCCGATCTATGATAGCATGAGTCTGAATGCAAAACAGTTCCT<br>GGAAATTTGTGGTGGTTCACATAGCTGTGCATGTAGCGGTAATAGCA<br>ATCAGGCACTGATCGGTATGAAAGGTGTTGCATGGATGAAACGCTTT<br>ATGGATAATGATACCCGCTATAGCCAGTTTGCCTGCGAAAATCCGAA<br>TAGCACCGCAGTTTGTGATTTTCGTACCGCAAATTGTAGCCTGGAAC<br>ATCATCACCATCATCATTAA | MQTNPYARGPNPTAASLEASAGP<br>FTVRSFTVARPVGYGAGTVYYPT<br>NAGGTVGAIAIVPGYTATQSSIN<br>WWGPRLASHGFVVITIDTNSTLD<br>KPESRSSQQMAALRQVASLNGTS<br>SSPIYGKVDTARGGVMGWSMGGG<br>GSLISAANNPSLKAAAVMAPXHS<br>STNFSSVTVPTLIFACENDRIAP<br>VKESALPIYDSMSLNAKQFLEIC<br>GGSHSCACSGNSNQALIGMKGVA<br>WMKRFMDNDTRYSQFACENPNST<br>AVCDFRTANCSLEHHHHHH |

| | | |
|---|---|---|
| LCC-ICCG | ATGAGCAATCCGTATCAGCGTGGTCCGAATCCGACACGTAGCGCACT | MSNPYQRGPNPTRSALTADGPFS |
| | GACCGCAGATGGTCCGTTTAGCGTTGCAACCTATACCGTTAGCCGTC | VATYTVSRLSVSGFGGGVIYYPT |
| | TGAGCGTTAGCGGTTTTGGTGGTGGTGTTATCTATTATCCGACCGGC | GTSLTFGGIAMSPGYTADASSLA |
| | ACCAGCCTGACCTTTGGTGGTATTGCAATGAGTCCGGGTTATACAGC | WLGRRLASHGFVVLVINTNSRFD |
| | AGATGCAAGCAGCCTGGCATGGCTGGGTCGTCGTCTGGCAAGCCATG | GPDSRASQLSAALNYLRTSSPSA |
| | GTTTTGTTGTTCTGGTGATTAATACCAACAGCCGTTTTGATGGTCCG | VRARLDANRLAVAGHSMGGGGTL |
| | GATAGCCGTGCAAGCCAGCTGAGCGCAGCACTGAATTATCTGCGTAC | RIAEQNPSLKAAVPLTPWHTDKT |
| | CAGCAGTCCGAGCGCAGTTCGTGCACGTCTGGATGCAAATCGTCTGG | FNTSVPVLIVGAEADTVAPVSQH |
| | CCGTTGCAGGTCATAGCATGGGTGGCGGTGGCACCCTGCGTATTGCA | AIPFYQNLPSTTPKVYVELCNAS |
| | GAACAGAATCCGAGCCTGAAAGCAGCAGTTCCGCTGACACCGTGGCA | HIAPNSNNAAISVYTISWMKLWV |
| | TACCGATAAAACCTTTAATACCAGCGTTCCGGTTCTGATTGTTGGTG | DNDTRYRQFLCNVNDPALCDFRT |
| | CAGAAGCAGATACCGTTGCACCGGTTAGCCAGCATGCAATTCCGTTT | NNRHCQLEHHHHHH |
| | TATCAGAATCTGCCGAGCACCACACCGAAAGTTTATGTTGAACTGTG | |
| | TAATGCCAGCCATATTGCACCGAATAGCAATAATGCAGCCATTAGCG | |
| | TTTATACCATCAGCTGGATGAAACTGTGGGTTGATAATGATACCCGT | |
| | TATCGTCAGTTTCTGTGCAATGTTAATGATCCGGCACTGTGTGATTT | |
| | TCGTACCAATAATCGTCATTGTCAGCTGGAACATCATCACCACCATC | |
| | ATTAA | |
| LCC-ICCG 190TAG | ATGAGCAATCCGTATCAGCGTGGTCCGAATCCGACACGTAGCGCACT | MSNPYQRGPNPTRSALTADGPFS |
| | GACCGCAGATGGTCCGTTTAGCGTTGCAACCTATACCGTTAGCCGTC | VATYTVSRLSVSGFGGGVIYYPT |
| | TGAGCGTTAGCGGTTTTGGTGGTGGTGTTATCTATTATCCGACCGGC | GTSLTFGGIAMSPGYTADASSLA |
| | ACCAGCCTGACCTTTGGTGGTATTGCAATGAGTCCGGGTTATACAGC | WLGRRLASHGFVVLVINTNSRFD |
| | AGATGCAAGCAGCCTGGCATGGCTGGGTCGTCGTCTGGCAAGCCATG | GPDSRASQLSAALNYLRTSSPSA |
| | GTTTTGTTGTTCTGGTGATTAATACCAACAGCCGTTTTGATGGTCCG | VRARLDANRLAVAGHSMGGGGTL |
| | GATAGCCGTGCAAGCCAGCTGAGCGCAGCACTGAATTATCTGCGTAC | RIAEQNPSLKAAVPLTPXHTDKT |
| | CAGCAGTCCGAGCGCAGTTCGTGCACGTCTGGATGCAAATCGTCTGG | FNTSVPVLIVGAEADTVAPVSQH |
| | CCGTTGCAGGTCATAGCATGGGTGGCGGTGGCACCCTGCGTATTGCA | AIPFYQNLPSTTPKVYVELCNAS |
| | GAACAGAATCCGAGCCTGAAAGCAGCAGTTCCGCTGACACCGTAGCA | HIAPNSNNAAISVYTISWMKLWV |
| | TACCGATAAAACCTTTAATACCAGCGTTCCGGTTCTGATTGTTGGTG | DNDTRYRQFLCNVNDPALCDFRT |
| | CAGAAGCAGATACCGTTGCACCGGTTAGCCAGCATGCAATTCCGTTT | NNRHCQLEHHHHHH |
| | TATCAGAATCTGCCGAGCACCACACCGAAAGTTTATGTTGAACTGTG | |
| | TAATGCCAGCCATATTGCACCGAATAGCAATAATGCAGCCATTAGCG | |
| | TTTATACCATCAGCTGGATGAAACTGTGGGTTGATAATGATACCCGT | |
| | TATCGTCAGTTTCTGTGCAATGTTAATGATCCGGCACTGTGTGATTT | |
| | TCGTACCAATAATCGTCATTGTCAGCTGGAACATCATCACCACCATC | |
| | ATTAA | |
| Kubu-PETase | ATGGCAGATCAGGTTGGTCAGGCACCGACCGCAGCAAATATTACCGG | MADQVGQAPTAANITGDGSFATA |
| | TGATGGTAGCTTTGCAACCGCAAGCGCACCGATTACCAATCAGACCG | SAPITNQTGFGGGTVYYPTAAGT |
| | GTTTTGGTGGTGGCACCGTTTATTATCCGACAGCAGCAGGCACCTAT | YPVVAVVPGFVSRWSQISWLGPR |
| | CCGGTTGTTGCAGTTGTTCCGGGTTTTGTTAGCCGTTGGAGCCAGAT | VASWGFVVVGADTNSGFDSPSSR |
| | TAGCTGGCTGGGTCCGCGTGTTGCAAGCTGGGGTTTTGTGGTTGTTG | ADQLLAALNWAVNSAPAAVRGKV |
| | GTGCAGATACCAATAGCGGTTTTGATAGCCCGAGCAGCCGTGCAGAT | DGTRRGVAGWSMGGGGTLEALCK |
| | CAGCTGCTGGCAGCACTGAATTGGGCAGTTAATAGCGCACCGGCAGC | DTTGTVKAGIPLAPWHIGQDFSC |
| | AGTTCGTGGTAAAGTTGATGGCACCCGTCGTGGTGTTGCAGGTTGGA | VTKPVFIVGAQNDTIAPPAQHAV |
| | GCATGGGTGGTGGTGGTACACTGGAAGCACTGTGTAAAGATACCACC | PFYNAAAGPKSYLELCGASHFFP |
| | GGCACCGTTAAAGCAGGTATTCCGCTGGCACCGTGGCATATTGGTCA | TTANPTVSRAMVSWLKRFVSSDD |
| | GGATTTTAGCTGTGTTACCAAACCGGTGTTTATTGTGGGTGCACAGA | RFTPFTCGFAGASVCAFRSTACL |
| | ATGATACCATTGCACCGCCTGCACAGCATGCAGTTCCGTTTTATAAC | EHHHHHH |
| | GCAGCAGCCGGTCCGAAAAGCTATCTGGAACTGTGTGGTGCAAGCCA | |
| | TTTCTTTCCGACCACCGCAAATCCGACCGTTAGCCGTGCCATGGTGA | |
| | GCTGGCTGAAACGTTTTGTGAGCAGTGATGATCGTTTTACCCCGTTT | |
| | ACCTGTGGTTTTGCCGGTGCAAGCGTTTGTGCATTTCGTAGCACCGC | |
| | ATGTCTGGAACATCATCACCATCATCATTAA | |
| Kubu-PETase 189TAG | ATGGCAGATCAGGTTGGTCAGGCACCGACCGCAGCAAATATTACCGG | MADQVGQAPTAANITGDGSFATA |
| | TGATGGTAGCTTTGCAACCGCAAGCGCACCGATTACCAATCAGACCG | SAPITNQTGFGGGTVYYPTAAGT |
| | GTTTTGGTGGTGGCACCGTTTATTATCCGACAGCAGCAGGCACCTAT | YPVVAVVPGFVSRWSQISWLGPR |
| | CCGGTTGTTGCAGTTGTTCCGGGTTTTGTTAGCCGTTGGAGCCAGAT | VASWGFVVVGADTNSGFDSPSSR |
| | TAGCTGGCTGGGTCCGCGTGTTGCAAGCTGGGGTTTTGTGGTTGTTG | ADQLLAALNWAVNSAPAAVRGKV |
| | GTGCAGATACCAATAGCGGTTTTGATAGCCCGAGCAGCCGTGCAGAT | DGTRRGVAGWSMGGGGTLEALCK |
| | CAGCTGCTGGCAGCACTGAATTGGGCAGTTAATAGCGCACCGGCAGC | DTTGTVKAGIPLAPXHIGQDFSC |
| | AGTTCGTGGTAAAGTTGATGGCACCCGTCGTGGTGTTGCAGGTTGGA | VTKPVFIVGAQNDTIAPPAQHAV |
| | GCATGGGTGGTGGTGGTACACTGGAAGCACTGTGTAAAGATACCACC | PFYNAAAGPKSYLELCGASHFFP |
| | GGCACCGTTAAAGCAGGTATTCCGCTGGCACCGTAGCATATTGGTCA | TTANPTVSRAMVSWLKRFVSSDD |

| | | |
|---|---|---|
| | GGATTTTAGCTGTGTTACCAAACCGGTGTTTATTGTGGGTGCACAGA<br>ATGATACCATTGCACCGCCTGCACAGCATGCAGTTCCGTTTTATAAC<br>GCAGCAGCCGGTCCGAAAAGCTATCTGGAACTGTGTGGTGCAAGCCA<br>TTTCTTTCCGACCACCGCAAATCCGACCGTTAGCCGTGCCATGGTGA<br>GCTGGCTGAAACGTTTTGTGAGCAGTGATGATCGTTTTACCCCGTTT<br>ACCTGTGGTTTTGCCGGTGCAAGCGTTTGTGCATTTCGTAGCACCGC<br>ATGTCTGGAACATCATCACCATCATCATTAA | RFTPFTCGFAGASVCAFRSTACL<br>EHHHHHH |
| Kubu-PETase 218S | ATGGCAGATCAGGTTGGTCAGGCACCGACCGCAGCAAATATTACCGG<br>TGATGGTAGCTTTGCAACCGCAAGCGCACCGATTACCAATCAGACCG<br>GTTTTGGTGGTGGCACCGTTTATTATCCGACAGCAGCAGGCACCTAT<br>CCGGTTGTTGCAGTTGTTCCGGGTTTTGTTAGCCGTTGGAGCCAGAT<br>TAGCTGGCTGGGTCCGCGTGTTGCAAGCTGGGGTTTTGTGGTTGTTG<br>GTGCAGATACCAATAGCGGTTTTGATAGCCCGAGCAGCCGTGCAGAT<br>CAGCTGCTGGCAGCACTGAATTGGGCAGTTAATAGCGCACCGGCAGC<br>AGTTCGTGGTAAAGTTGATGGCACCCGTCGTGGTGTTGCAGGTTGGA<br>GCATGGGTGGTGGTGGTACACTGGAAGCACTGTGTAAAGATACCACC<br>GGCACCGTTAAAGCAGGTATTCCGCTGGCACCGTGGCATATTGGTCA<br>GGATTTTAGCTGTGTTACCAAACCGGTGTTTATTGTGGGTGCACAGA<br>ATGATACCATTGCACCGCCTGCACAGAGCGCAGTTCCGTTTTATAAC<br>GCAGCAGCCGGTCCGAAAAGCTATCTGGAACTGTGTGGTGCAAGCCA<br>TTTCTTTCCGACCACCGCAAATCCGACCGTTAGCCGTGCCATGGTGA<br>GCTGGCTGAAACGTTTTGTGAGCAGTGATGATCGTTTTACCCCGTTT<br>ACCTGTGGTTTTGCCGGTGCAAGCGTTTGTGCATTTCGTAGCACCGC<br>ATGTCTGGAACATCATCACCATCATCATTAA | MADQVGQAPTAANITGDGSFATA<br>SAPITNQTGFGGGTVYYPTAAGT<br>YPVVAVVPGFVSRWSQISWLGPR<br>VASWGFVVVGADTNSGFDSPSSR<br>ADQLLAALNWAVNSAPAAVRGKV<br>DGTRRGVAGWSMGGGGTLEALCK<br>DTTGTVKAGIPLAPWHIGQDFSC<br>VTKPVFIVGAQNDTIAPPAQSAV<br>PFYNAAAGPKSYLELCGASHFFP<br>TTANPTVSRAMVSWLKRFVSSDD<br>RFTPFTCGFAGASVCAFRSTACL<br>EHHHHHH |
| Kubu-PETase 218S/189TAG | ATGGCAGATCAGGTTGGTCAGGCACCGACCGCAGCAAATATTACCGG<br>TGATGGTAGCTTTGCAACCGCAAGCGCACCGATTACCAATCAGACCG<br>GTTTTGGTGGTGGCACCGTTTATTATCCGACAGCAGCAGGCACCTAT<br>CCGGTTGTTGCAGTTGTTCCGGGTTTTGTTAGCCGTTGGAGCCAGAT<br>TAGCTGGCTGGGTCCGCGTGTTGCAAGCTGGGGTTTTGTGGTTGTTG<br>GTGCAGATACCAATAGCGGTTTTGATAGCCCGAGCAGCCGTGCAGAT<br>CAGCTGCTGGCAGCACTGAATTGGGCAGTTAATAGCGCACCGGCAGC<br>AGTTCGTGGTAAAGTTGATGGCACCCGTCGTGGTGTTGCAGGTTGGA<br>GCATGGGTGGTGGTGGTACACTGGAAGCACTGTGTAAAGATACCACC<br>GGCACCGTTAAAGCAGGTATTCCGCTGGCACCGTAGCATATTGGTCA<br>GGATTTTAGCTGTGTTACCAAACCGGTGTTTATTGTGGGTGCACAGA<br>ATGATACCATTGCACCGCCTGCACAGAGCGCAGTTCCGTTTTATAAC<br>GCAGCAGCCGGTCCGAAAAGCTATCTGGAACTGTGTGGTGCAAGCCA<br>TTTCTTTCCGACCACCGCAAATCCGACCGTTAGCCGTGCCATGGTGA<br>GCTGGCTGAAACGTTTTGTGAGCAGTGATGATCGTTTTACCCCGTTT<br>ACCTGTGGTTTTGCCGGTGCAAGCGTTTGTGCATTTCGTAGCACCGC<br>ATGTCTGGAACATCATCACCATCATCATTAA | MADQVGQAPTAANITGDGSFATA<br>SAPITNQTGFGGGTVYYPTAAGT<br>YPVVAVVPGFVSRWSQISWLGPR<br>VASWGFVVVGADTNSGFDSPSSR<br>ADQLLAALNWAVNSAPAAVRGKV<br>DGTRRGVAGWSMGGGGTLEALCK<br>DTTGTVKAGIPLAPXHIGQDFSC<br>VTKPVFIVGAQNDTIAPPAQSAV<br>PFYNAAAGPKSYLELCGASHFFP<br>TTANPTVSRAMVSWLKRFVSSDD<br>RFTPFTCGFAGASVCAFRSTACL<br>EHHHHHH |

[a] X indicates the position of azatryptophan.
